# Illuminating the uncharacterized regulatory genome of *E. coli* with massively parallel reporters

**DOI:** 10.1101/2025.05.13.653802

**Authors:** Tom Röschinger, Heun Jin Lee, Rosalind Wenshan Pan, Grace Solini, Kian Faizi, Baiyi Quan, Tsui-Fen Chou, Madhav Mani, Stephen Quake, Rob Phillips

## Abstract

All cells respond to changes in both their internal milieu and the environment around them through the regulation of their genes. Despite decades of effort, there remain huge gaps in our knowledge of both the function of many genes (the so-called y-ome) and how they adapt to changing environments via regulation. Here we describe a joint experimental and theoretical dissection of the regulation of a broad array of more than 100 biologically interesting genes in *E. coli* across 39 diverse environments, enabling us to identify the binding sites and transcription factors that mediate regulatory control. Using a combination of mutagenesis, massively parallel reporter assays, mass spectrometry, and tools from information theory and statistical physics, we go from complete ignorance of a promoter’s environment-dependent regulatory architecture to a quantitative description of its binding sites, candidate transcription factors that bind them where identifiable, and the conditions under which they act. As proof of principle of the biological insights to be gained from such a study, we chose a combination of genes from the y-ome, toxin-antitoxin pairs, and genes hypothesized to be part of regulatory modules; we discovered a host of new insights into their underlying regulatory landscape and resulting biological function. We highlight discoveries for y-ome genes, including transcription start sites and transcription factor binding sites at base-pair resolution, and their dependence on growth conditions.

## 2 Introduction

The discovery in the early 1960s that there are genes whose job it is to control other genes and how that control is exercised through environmental influences was heralded as “the second secret of life,” [1, 2] vastly expanding the original conception of the gene itself. It has now been more than sixty years since Jacob and Monod ushered in their repressor-operator model and the allied discovery of allosteric regulation [3, 4]. The regulatory landscape of the genome requires building a bridge between its base pairs, the molecules that bind to them, and its environmental context. However, despite a prodigious effort in the case of *Escherichia coli* [5–12], one of biology’s best studied model organisms, for roughly 60% of its genes, databases such as RegulonDB lack strong evidence for transcriptional regulation [13]. Further, roughly 40% of its genes are either partially characterized or not characterized at all. These genes of unknown function have been christened the y-ome [14–16].

The goal of our work is to systematically address these questions by providing a simultaneous promoter-by-promoter and high-throughput quantitative dissection of a variety of biologically interesting genes, as well as genes whose function or context have not yet been discovered. We place particular emphasis on the all-important question of how genes are regulated in response to a myriad of different environmental conditions, as transcription factors often change their activity based on internal or external signals, a set of interactions coined the “allosterome” [17]. Here, using a combination of mutagenesis, massively parallel reporter assays (MPRAs), mass spectrometry and statistical physics, we can go from complete ignorance of a promoter’s regulatory architecture to a quantitative description of its binding sites, the transcription factors that bind them, and the conditions under which they act (Figure 1). Our work builds on and is inspired by many brilliant studies using MPRAs [18–24].

**Figure 1.**
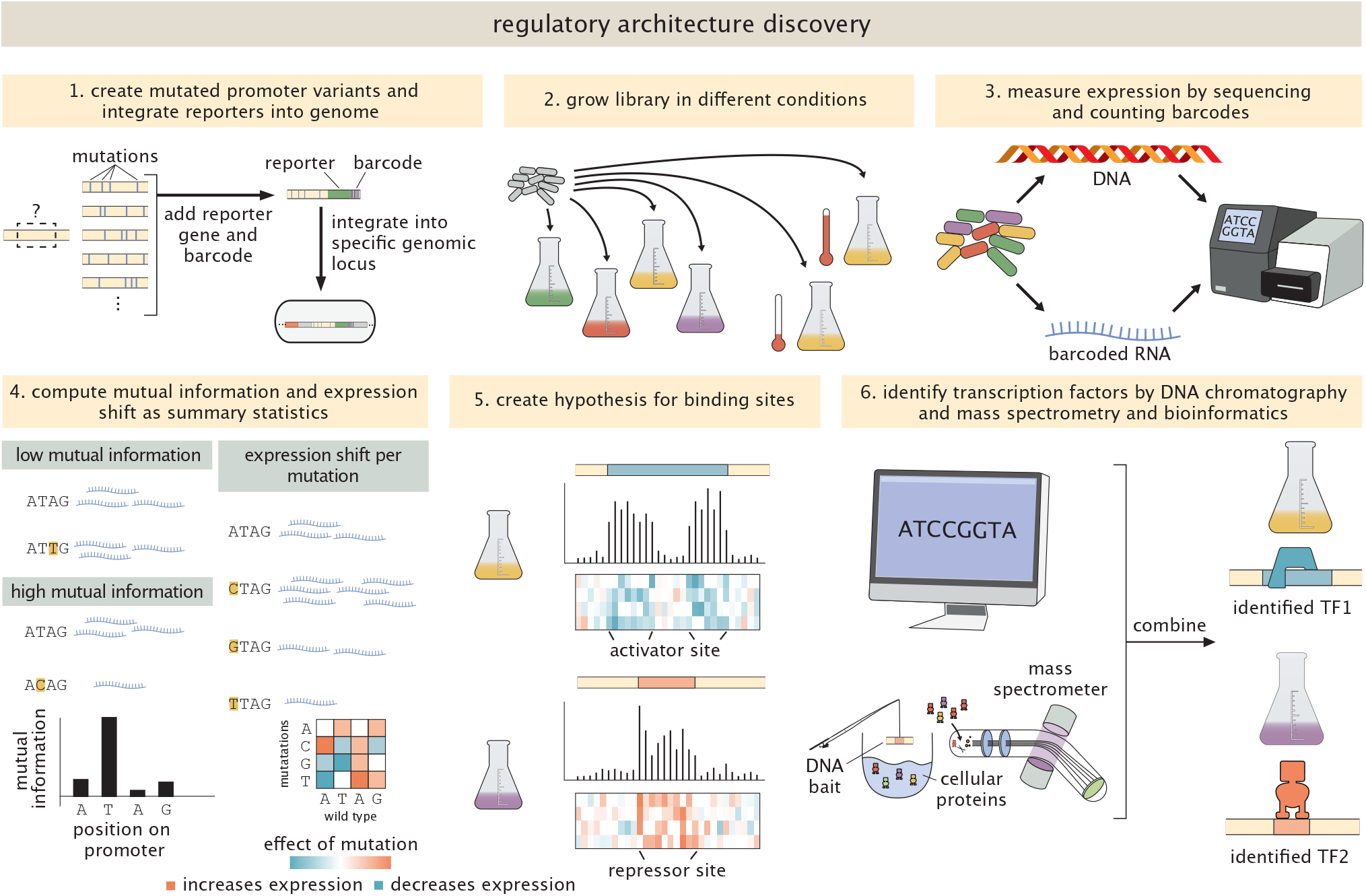
Discovering regulatory architectures. Experimental procedure for characterizing a previously uncharacterized gene. The six elements shown here provide a schematic of the steps needed to go from an uncharacterized promoter to one in which we have a well-defined, environmentally-dependent model of the binding sites, the allied transcription factors, and the energy matrix (shown under the information footprints) describing their binding interaction.

In particular, the present work is founded upon the MPRA studies known as Sort-Seq [6, 18, 19] and Reg-Seq [8], which share the philosophy of using mutated promoters and gene expression measurements in conjunction with information theory to generate high-throughput hypotheses for binding site locations. Together with these binding site hypotheses, both experimental and computational approaches are then used to determine which transcription factors bind them. For reasons we will explain in detail later in the paper, we note that the problem of figuring out which transcription factors bind to which binding sites is very challenging. Here, we developed and exploited a next generation version of Reg-Seq, including a streamlined protocol and genome integrated reporters, to study the regulatory architecture of more than 100 promoters in 39 different growth conditions, while paving the way to study the entire regulatory genome of an organism in one experiment. A detailed discussion of related methods and literature can be found in the Supplementary Information section 1.

To achieve such a condition-dependent dissection of the regulatory landscape, we adopt the protocol shown in Figure 1, which is a modified protocol developed previously [8] and described in detail in the supplementary information. Briefly, the procedure entails first using massively parallel reporter assays in conjunction with information theory and simple probabilistic models to generate hypotheses for the locations of binding sites. These binding sites are then used as a fishing hook for DNA chromatography and mass spectrometry to identify bound transcription factors or used for computational motif scanning [25] in which we compare our putative binding sites to databases of known binding sites. We note, however, that identifying the responsible transcription factor is the most challenging step in this pipeline. Mass spectrometry does not recover all binders, particularly when binding is strongly condition-dependent, and in several cases the responsible factor could not be determined despite multiple experimental approaches. In addition to identifying which transcription factors bind these putative binding sites, the goal is to ultimately infer a nucleotide-by-nucleotide binding energy matrix (for a deep analysis of this approach see [26]) as we have done in the past [6, 8, 18, 27], which predicts a promoter’s input-output function based on statistical mechanics [27–31].

As a proof of principle, we focused on 104 genes that struck us as particularly exciting, exploring their promoters across 39 growth conditions. Several genes have multiple known or predicted transcription start sites, yielding 119 promoter constructs in total. The genes span six categories. First, 16 “gold standards” with well-characterized regulatory architectures from previous experiments [6, 8], many of them famous case studies. Second, 18 genes identified as having high protein-abundance variance across conditions in a seminal quantitative proteome study [32], of which 9 had no annotated function at the time and are part of the y-ome [15]. Third, 13 additional y-ome genes lacking any functional description in EcoCyc [16]: identifying transcription factor binding sites at these promoters offers a starting point for inferring the pathways these genes participate in. Fourth, 18 genes from toxin/antitoxin systems, whose tightly controlled regulation responds to a range of cellular stresses [33]. Fifth, two newly identified iModulons [34] putatively controlled by YmfT and YgeV — both of which contain further y-ome genes — offering an opportunity to investigate how genes respond to environments as a collective. Sixth, 6 genes participating in feedforward loop motifs [35–38], a starting point for studying how environmental perturbations propagate through gene networks. The full list of genes is provided in Table S5. The diversity of regulatory discoveries that emerge from these experiments — from binding site verification to novel transcription factor identification to new transcription start sites — is summarized in Figure 2, which serves as a roadmap for the results that follow. Our ultimate objective is to extend this systematic analysis to every gene in *E. coli* (and later other organisms such as *P. aeruginosa*) under a broad array of environmental conditions.

**Figure 2.**
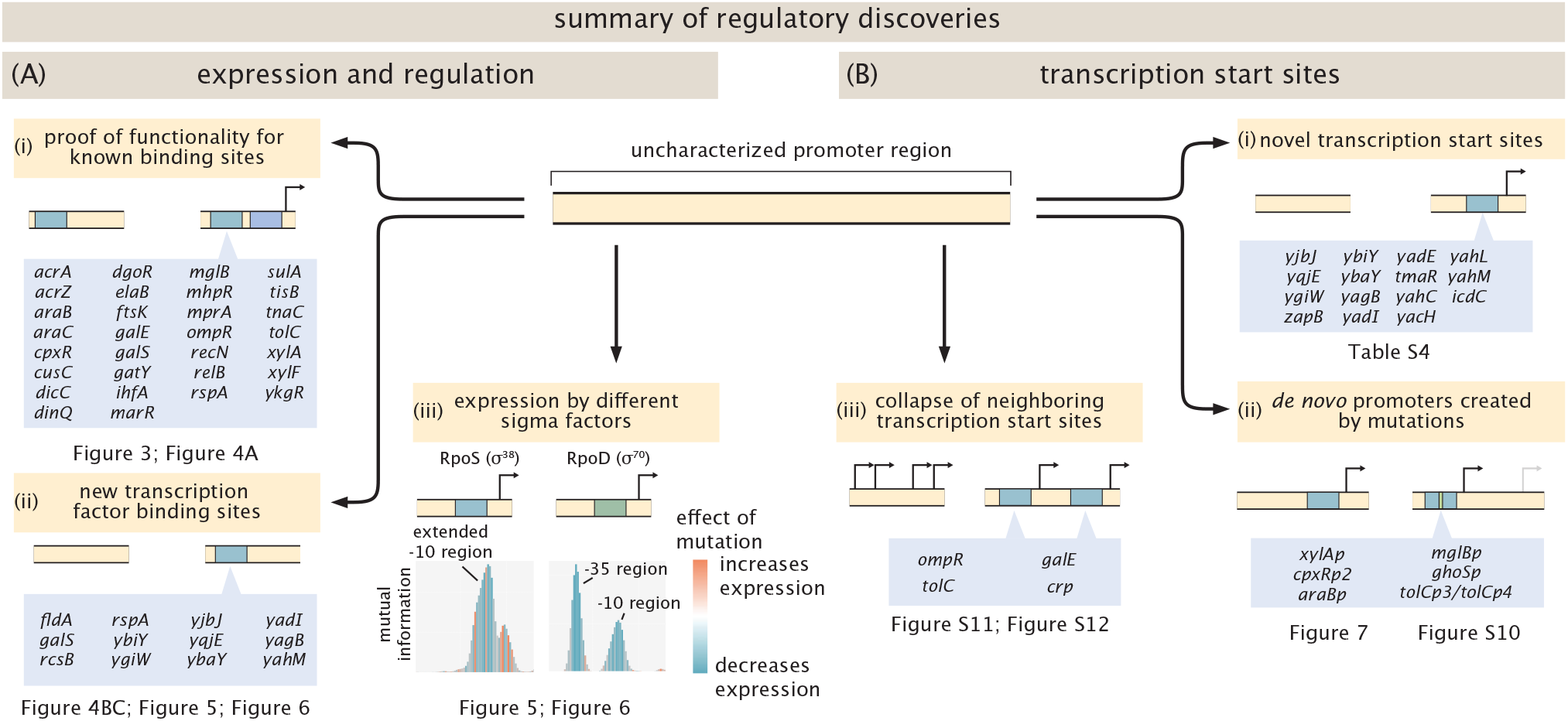
Types of discoveries made in Reg-Seq experiments. (A) The primary goal of Reg-Seq experiments is the discovery and verification of transcription factor binding sites. (i) Sites annotated from different methods are tested for functionality. (ii) Novel transcription factor binding sites are discovered. (iii) Expression by different sigma factors dependent on condition. (B) In addition to transcription factor binding sites, Reg-Seq experiments deliver results for transcription start sites as well. (i) Novel transcription start sites can be identified for genes without previously identified promoters. (ii) Mutations in existing promoters can lead to new, distinct transcription start sites, especially in conditions where the annotated site is not active. (iii) Neighboring annotated promoters can collapse to be a single transcription start site.

## 3 Results

We use a modified Reg-Seq [8] protocol with the long-term goal of enabling systematic discovery of regulatory architectures across large fractions of the genome. In brief, a pool of mutated variants was generated for each promoter. Each variant is cloned upstream of a reporter gene and a randomized DNA barcode. The entire library is genome integrated using a modified ORBIT protocol [39], which allowed us to perform genome integration at such scale. Genome integration eliminates plasmid copynumber variation and the need for antibiotic selection, while preserving library diversity (median 13 barcodes per variant). A detailed description of the library cloning procedure can be found in the SI Section S2. The full list of conditions, including media composition, supplements, and harvest times, is given in SI Section S2.5.3. All conditions were performed in at least two biological replicates. Per-gene results, including footprints, sequences, and condition-by-condition expression data, are collected in an interactive compendium SI file 1.

To identify binding sites from Reg-Seq data, we use mutual information footprints and expression shift matrices. Information footprints have been used with great success in previous experiments [8, 18], and a detailed exploration of information footprints can be found in the SI Section S3.

### Multidimensional discovery from Reg-Seq experiments

Our experiments yielded a richer set of discoveries than binding site verification alone. Across 119 promoters and 39 conditions, we identified 49 previously annotated binding sites as footprints, and discovered 13 novel binding sites, 5 of which we were able to assign to a specific transcription factor. Beyond binding sites, we identified 18 new transcription start sites for genes with no previously annotated promoter or only weak prior evidence, and found 6 cases where single mutations create entirely new promoters *de novo*, three of which are described in detail below and three more in the supplementary information. In a handful of cases, multiple annotated transcription start sites collapsed to a single active site. Figure 2 organizes these discoveries by type, and serves as a roadmap for the sections that follow, each of which highlights specific examples in more detail. A complete per-promoter accounting is provided in the supplementary information.

First, annotated transcription factor binding sites are tested for functionality (Figure 2A(i)). Binding of transcription factors alone does not imply that expression of a gene is altered, as has been shown in the case of PhoB [40]. Here, we investigate if modifying the binding site, and therefore, in most cases the binding energy, leads to a measurable change in expression from the reporter gene. If we observe the binding site in the information footprint, it verifies binding and regulation of the corresponding promoter. There are different reasons for certain binding sites to not be identified by our method. For one, some transcription factors require specific conditions that we did not test in our experiments, such as TorR, which is activated by the TorS/TorT two-component system in response to TMAO (trimethylamine N-oxide) [41]. A systematic class of misses are sites that require distant sites in the genome for function, such as DNA looping. Since reporter constructs are 160 bases long around the transcription start site, distant sites are not included in the reporter construct. A per-promoter accounting of every annotated binding site, with rationale for each exclusion or miss, is provided in the section “All identified binding sites” below and in SI Section S4. Here, missed binding sites are opportunities to learn how to improve the method in further iterations.

Beyond verification of annotated sites, we identify novel transcription factor binding sites (Figure 2A(ii)), discussed in detail in the section “All identified binding sites” below. Another interesting result is the difference in footprints we observe for sigma factors. Transcriptional regulation is not only instigated by transcription factors, but on an even higher level by sigma factors. Their condition dependence and differential interaction with transcription factors govern which sets of genes are expressed. As shown in Figure 2A(iii), we find distinct footprints for *σ*^70^ and *σ*^38^, especially in stationary phase and slow growing conditions.

In addition to the identification of binding sites, Reg-Seq delivers discoveries about transcription start sites. First, using a computational model for *σ*^70^ binding [42], we predicted transcription start sites for genes that did not have previously identified sites in databases (Figure 2B(i)). Most confirmed sites were active specifically in stationary phase, showing information footprints for *σ*^38^, highlighting the lack of annotation for stationary phase, and that many y-genes might be expressed under such conditions. This reflects the substantial overlap between the *σ*^70^ and *σ*^38^ − 10 consensus sequences, which means a *σ*^70^-trained predictor also catches *σ*^38^-driven promoters.

As shown in Figure 2B(iii), few genes we studied had multiple, neighboring transcription start sites annotated in databases, and in some cases the promoters showed no sign of two different initiation events. We also observed promoters created *de novo* by individual mutations (Figure 2B(ii)), which we explore in more detail below.

### Condition specific resolution of binding sites

Transcriptional regulation is very dependent on the environment cells are in. The transcriptional repressor LexA is involved in the cellular stress response to DNA damage, also known as the SOS response [43]. LexA has 42 annotated binding sites in EcoCyc, and it has been observed that LexA and many of its regulatory targets are upregulated when treated with 2.5 mM hydrogen peroxide (H_2_O_2_) for 10 minutes [44]. The reason for this upregulation is that in the presence of hydrogen peroxide, the RecA-ssDNA filament stimulates LexA autoproteolysis, causing its dissociation from its binding sites and subsequent expression of its target genes [45, 46]. There are five genes in our pool that have annotated binding sites for LexA in their promoters: *tisB, sulA, recN, dinQ*, and *ftsK*. As shown in Figure 3, the information footprint shows the binding site for LexA in the promoters *sulAp* and *tisBp* as peaks of mutual information. In the binding sites, mutations mostly increase expression, since binding of LexA is weakened if the site is mutated, and repression is reduced. When hydrogen peroxide is added, the repressor peaks disappear, showing that LexA no longer represses the promoters. To confirm we observe binding by LexA, we performed DNA chromatography and mass spectrometry as described previously [6, 8] (details are shown in SI section S2.7). We used both lysates from cells grown in stationary phase, and from cells that were induced with hydrogen peroxide. Stationary-phase lysate, in which LexA is constitutively present, gives clear enrichment for LexA, confirming specific binding to the bait sequence. Lysate from cells induced with hydrogen peroxide, in which LexA is degraded, gives no enrichment, consistent with the loss of repressor footprint observed in the Reg-Seq data. These results show us how we can identify the specific condition a regulator responds to. LexA control of tisAB had been inferred from SOS-induction patterns [47] and from a sequence-annotated LexA box [48], and ysdA was captured as a strong LexA ChIP-chip target without binding-site resolution [49]; our results provide base-pair-resolution evidence of LexA binding at the tisB promoter and directly demonstrate that this binding represses transcription. When cells are induced with gentamicin, LexA repression at *tisBp* is weakened (Figure S13), suggesting a route by which gentamicin exposure leads to selective *tisB* expression and the previously reported gentamicin tolerance [50]. How *tisB* is selectively derepressed without broader SOS induction remains unclear.

**Figure 3.**
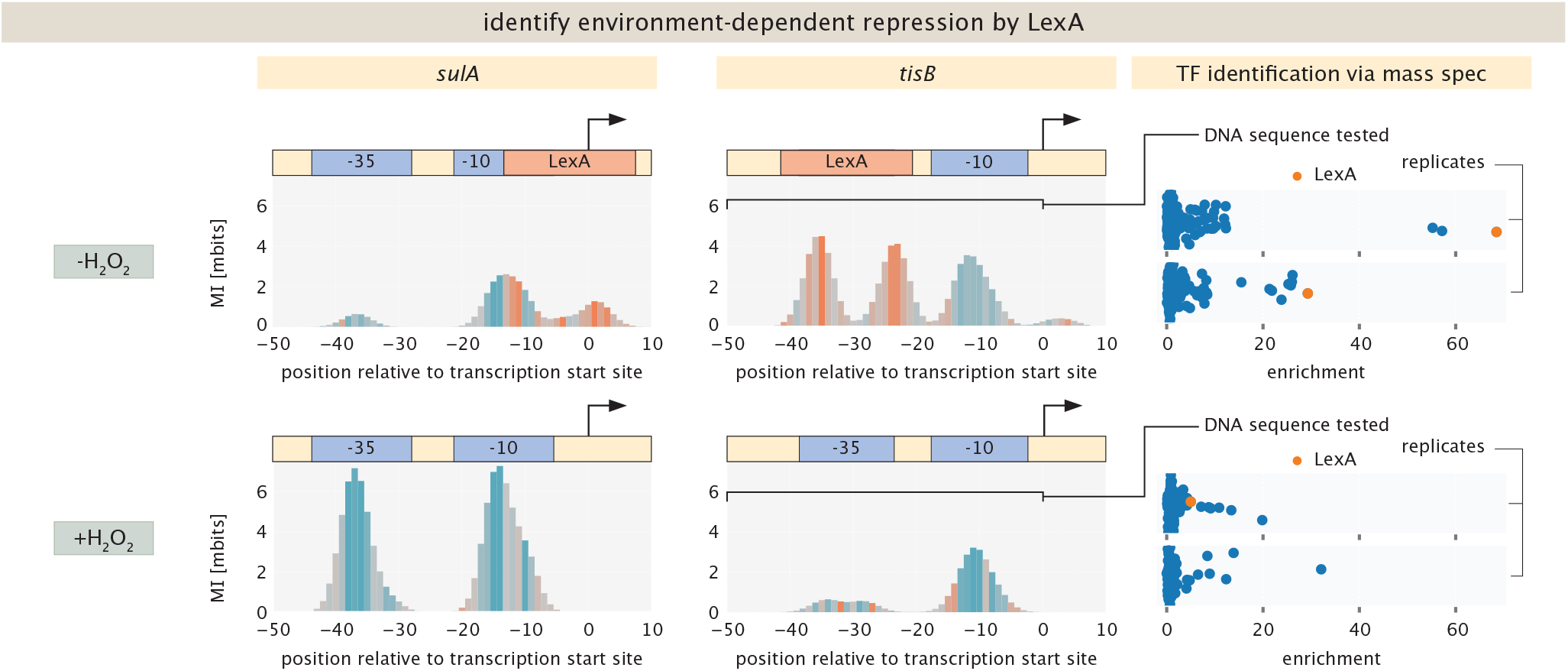
Identification of condition specific regulation by Reg-Seq. Repression by LexA of the *sulAp* and *tisBp* promoters. During exposure to hydrogen peroxide, LexA is catabolized and no repressor signal is observed. LexA is identified as binder in mass-spec experiments.

The CpxR-CpxA two-component system regulates the response to inner-membrane stress [51, 52], with CpxR autoactivating its own expression in the phosphorylated state [53]. We recover the CpxR activator site at the *cpxR* promoter under both gentamicin and copper sulfate (500 μM and 2 mM) induction (Figure S13), and confirm CpxR binding by mass spectrometry under both conditions (Figure S14). We find an analogous footprint at *yqaE*, which carries an annotated CpxR site. The *mtnp* promoter, which also has an annotated CpxR site [54], shows no activation in any condition we tested — either because the inducing condition is absent from our panel, or the site is not functional under the conditions probed.

### All identified binding sites

Figure 4 summarizes all binding sites identified in our experiments, organized by whether they were previously annotated and whether the responsible transcription factor was identified. Across 43 promoters with previously annotated regulation, RegulonDB lists 143 binding-site entries within our 160 bp mutagenesis windows. Of these, 40 cannot be tested in our assay — either because the inducing condition is absent from our panel, the cognate TF binds degenerately (Fis, HU, CspA), the cooperative partner falls outside the window (e.g. AraC looping), or the annotation overlaps an unresolvable cluster (ompFp). Of the 43 promoters reconciled, 2 contain no testable annotations — every in-window site is either degenerate (Fis/HU/CspA) or depends on a cooperative partner outside the reporter window — leaving 41 promoters and 103 testable annotations at which recovery can be assessed. We recover 44 (43%) as clear footprints and 5 more as weak footprints, for a combined recovery of 49/103 (48%). At the level of promoters, 18/41 are fully recovered, 15 are partially recovered, and 8 yield no recovered sites. A per-promoter accounting of every annotation, with the rationale for each exclusion or miss, is provided in Supplementary Note 1.

**Figure 4.**
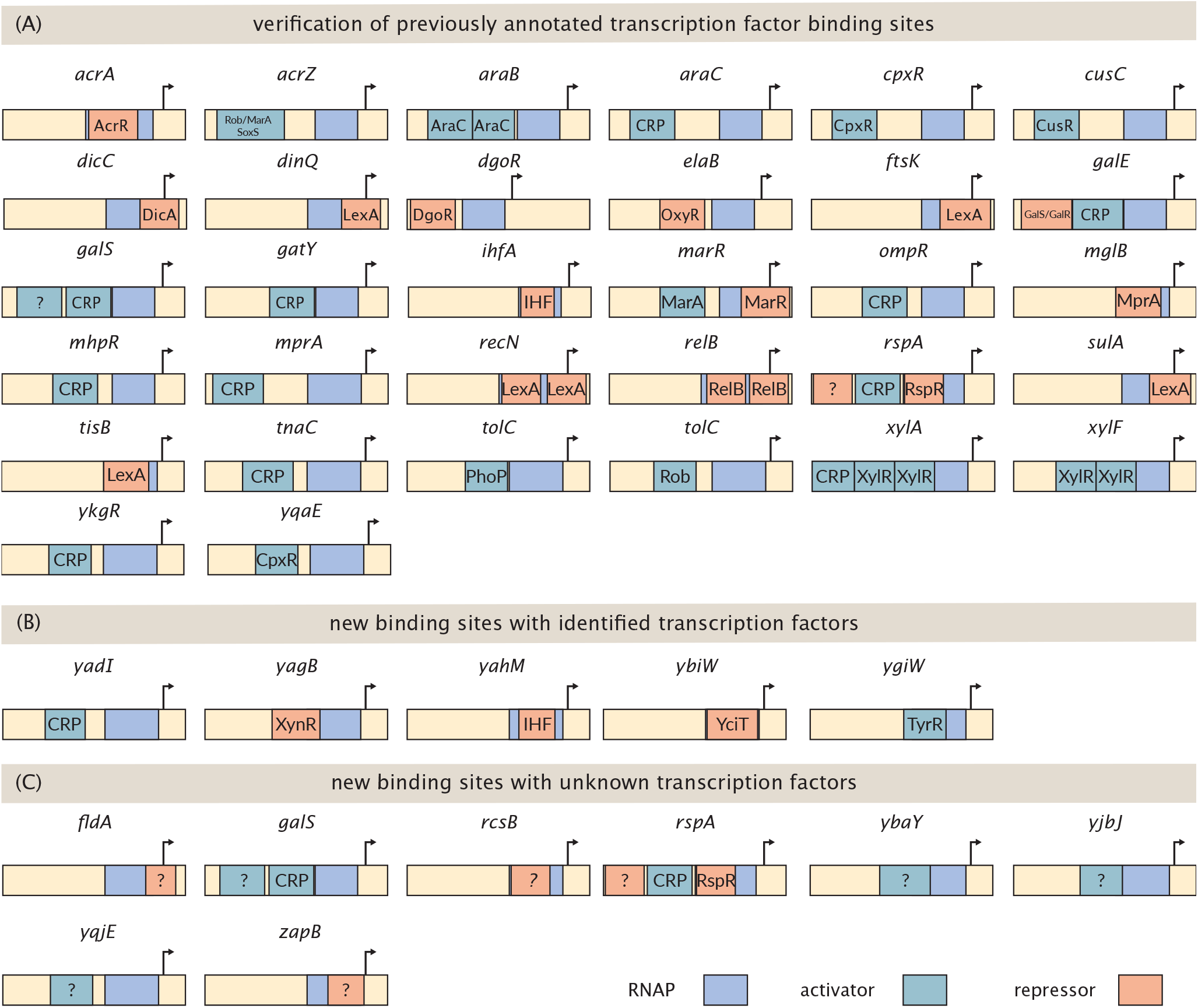
All identified binding sites. (A) Previously annotated binding sites that were verified in our experiments. (B) New binding sites for which we identified the transcription factors. (C) New binding sites where the transcription factors were not identified.

The 54 misses are not uniform in interpretation. A substantial fraction are W-confidence annotations supported only by sequence similarity to a TF consensus (e.g. CRP at aceBp, MarA at yncEp, OxyR at znuCp), where the absence of a Reg-Seq footprint argues that the predicted site is not a physiologically active regulatory interaction. Others occur at promoters whose activity in our reporter is too low to resolve repressor footprints, or under conditions where the cognate TF may not reach the activity regime required for binding (e.g. OxyR under transient H_2_O_2_ exposure). A third class reflects cooperative regulatory architectures whose partners fall outside the 160 bp reporter window and so cannot be reconstituted in our constructs. Each miss thus points to a specific follow-up: re-examination of the annotation, an expanded condition panel, or an extended construct design.

As shown in Figure 4(B), we identified five new binding sites where we identified the transcription factors binding as well. The gene *ygiW* lacks an annotated transcription start site. It has been suggested to be involved in the response to hydrogen peroxide [55] and is upregulated in stationary phase [32, 56]. We predicted a promoter region, and found activation in stationary phase. Mass spectrometry experiments show enrichment for the transcription factor TyrR (Figure S14). We did not find activation from this promoter after addition of hydrogen peroxide, suggesting that regulation of this gene does not respond to this condition, or there is another promoter region for this gene which responds specifically to this condition.

Another y-ome gene that we considered is the *ybiYW* operon, which until now has had no functional annotation. Using our broad suite of environmental conditions, we found an active transcription start site when cells are grown anaerobically both with and without supplemented nitrate, as shown in Figure S14. We also found expression when minimal media is supplemented with glucose and subinhibitory concentrations of ampicillin, but only in one replicate, which could indicate that in this specific experiment cells entered anaerobic growth conditions. As shown in Figure S14, DNA chromatography and mass spectrometry with lysate from cells grown in minimal media with glucose shows high enrichment for the transcription factor YciT, suggesting YciT binds in the vicinity of the −5 site relative to the predicted transcription start site. The position of the binding site, overlapping the RNAP footprint, is consistent with YciT acting as a repressor — contrary to its annotation as an activator in RegulonDB but in agreement with transcriptomic evidence that *ybiY* and *ybiW* are upregulated in a Δ*yciT* strain [9]. Combined with the anaerobic activation of the promoter, the simplest model is that YciT represses *ybiYW* under aerobic conditions, with repression relieved under anaerobiosis. This finding suggests that YciT and YbiY-YbiW are involved in the cell’s response to anaerobic conditions, which has not been reported before. As shown in Figure S14, the information footprint suggests binding of an activator around the −20 region, but we found no candidate transcription factors through the mass spectrometry experiments or computational motif scanning. It should be noted that cells grown aerobically were used for DNA chromatography, as producing the needed amount of cell lysate anaerobically was technically out of scope. These results agree with previous work where YciT was found to bind upstream of *ybiY* [9]. The predicted binding site was about 100 bases more upstream than the binding site we identified, and regulation was assumed to depend on osmotic shock, which we cannot confirm here.

The gene *yagB* lies within the CP4-6 cryptic prophage element and is transcribed as part of the *yagB* /*insX* /*yagA* unit, divergent from *yagEF*. The shared intergenic region contains a binding site for the IclR-family repressor XynR (YagI), which was shown to repress both divergent transcription units [57]. The *yagEF* genes encode enzymes for D-xylonate catabolism, and XynR activity is modulated by xylonate as an effector ligand. The information footprints in Figure S14 reveal a single broad peak at approximately position −40 relative to the predicted transcription start site, containing both blue and orange signal. This mixed signal is consistent with overlapping contacts from *σ*^*S*^ (blue; mutations decrease expression) and a repressor (orange; mutations increase expression), indicating that the two proteins bind the same DNA region. The position of the *σ*^*S*^ contact places the actual transcription start site approximately 25 bp upstream of the computational prediction. Mass spectrometry identified two proteins enriched at this promoter, as shown in Figure S14. The first is XynR, consistent with the binding site identified [57]. The second is H-NS, the nucleoid-associated protein known to silence horizontally acquired DNA elements in *E. coli*, including cryptic prophages [58, 59]. H-NS enrichment at a prophage-encoded promoter is expected given this global silencing role. However, we do not observe a distinct H-NS footprint separate from the XynR/sigma factor peak, and we cannot distinguish whether H-NS binding overlaps entirely with XynR or simply produces no measurable expression effect.

The gene *yahM* is completely uncharacterized. While no transcription start site is annotated in EcoCyc, one was identified in high-throughput experiments [60]. Here we confirm the transcription start site with an offset of about +15 bases, and identify binding of IHF as repressor, which shows especially in information footprints from growth on arabinose, as shown in Figures 4(B) and S14. Both IHF subunits (IhfA and IhfB) are recovered by mass spectrometry, consistent with binding of the intact IHF heterodimer.

For one further y-ome gene, *yadI*, we identified a transcription start site and a binding site for the global regulator CRP. Because CRP is one of the most-studied regulators in *E. coli* and the discovery illustrates what Reg-Seq can do at uncharacterized loci, we describe this case in detail in the next section.

We further identified 8 binding sites where we did not identify the corresponding transcription factors. The information footprint for the *fldA* shows a repressor binding site in phenazine methosulfate around the transcription start site, as shown in Figure S15. In the promoter for *galS* we find another activator binding site around −95, which shows especially in stationary phase or growth on acetate or galactose. A repressor binding site is visible in the *rcsBp2* promoter, most visible for growth on acetate, as shown in Figure S15. Additional repressor or activator footprints without an identified transcription factor were observed at the *zapBp* and *rspAp* promoters; details are in Figure S15.

The identified binding sites for *yjbJ, ybaY* and *yqjE* are discussed in more detail below.

### Identification of a CRP binding site

CRP is one of the most ubiquitous regulators in *E. coli*, with more than 300 annotated binding sites around the genome. There are 13 promoters with known CRP-binding sites in our pool. Of these, 8 group together in the expression profile with a large increase in expression for growth in, e.g., acetate, arabinose, and LB. Three of the exceptions, *araBp, xylAp*, and *xylFp*, are promoters of genes involved in the metabolism of specific carbon sources (arabinose and xylose), and we observe increased expression in these conditions. For one of the genes without an annotated function or a transcription start site, *yadI*, we found a putative promoter that groups with CRP-activated promoters. We find an active transcription start site, and a putative activator binding site in a few conditions, as shown in Figure 5(A), especially in stationary phase, as well as minimal media with acetate and galactose. Mass spectrometry did not identify CRP at *yadIp* or control CRP promoters (SI Section S5.2). To assess whether the sequence at the activator peak is consistent with a CRP binding site, we constructed a PWM from 269 CRP binding sites with strong or confirmed evidence in RegulonDB. The best-scoring 20-mer within the activator peak region (GTTTTGACGGCTATCACCCT, positions −45 to −26, dyad axis at −34.5) received a log-odds score of 8.69, placing it at the 36th percentile of known CRP sites (Figure 5(B), left). This is well within the range of confirmed CRP targets (median = 10.38, range −17.6 to 19.1). As independent corroboration, we examined a CRP ChIP-exo dataset, where genome-wide CRP binding was measured for growth on glycerol minimal medium [61]. The *yadI* locus is included among the CRP ChIP-exo peak regions (Figure 5B, middle), and the signal shows the TSS-centered protection pattern that is described as characteristic of CRP-activated promoters [61], as opposed to the motif-centered profiles seen at repressor sites such as ArcA and Fur. This is consistent with active CRP-dependent transcription at this locus. Three lines of evidence — internally consistent Reg-Seq footprints (binding site, sequence match to the CRP PWM, and a confirmed 10 contact at the predicted TSS), and independent CRP ChIP-exo binding [61] — establish *yadI* as a CRP-activated gene.

**Figure 5.**
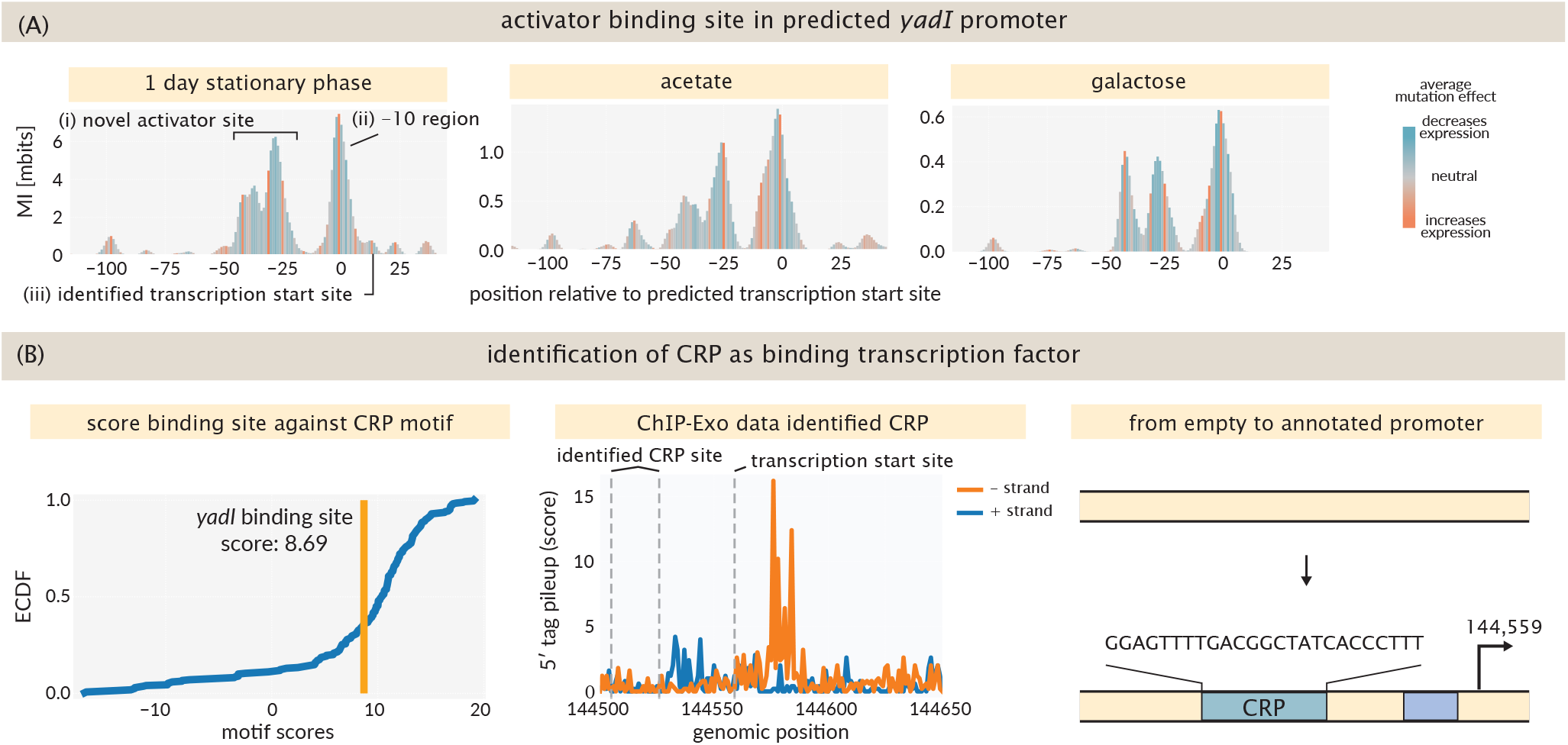
Discovery of novel CRP binding site. (A) Footprints for the predicted promoter for *yadI* show an activator site in slow growth conditions like stationary phase, galactose and acetate. (B) Identification of CRP as binding transcription factor. Evaluation of the CRP binding motif on all known CRP binding sites, as well as the identified binding site in the promoter for *yadI*. Chip-Exo data identifies *yadI* as hit, confirming activation of *yadI* by CRP.

As shown in Figure 5(B, right), the *yadI* case illustrates what Reg-Seq can do at uncharacterized loci. From a single experiment, we identify the transcription factor (CRP, supported by PWM scoring and ChIP-exo), the binding site position (−34.5), the promoter architecture (class II-like by the ChIP-exo protection pattern [61], with CRP overlapping the −35 region), the sigma factor contact (single-peak, extended −10), and the condition-specificity (stationary phase, galactose, and acetate, but not pentose sugars). The y-ome comprises roughly one-third of *E. coli* genes, and the regulatory logic of these genes is almost entirely unknown. Results like this demonstrate that Reg-Seq can systematically fill in this gap.

### Multiple y-genes respond strongly to osmotic shock

A group of six promoters in our pool is activated when cells are washed and grown for 1 h in LB supplemented with 750 mM sodium chloride. Three of these are y-genes without an annotated transcription start site prior to our experiments. The gene *yjbJ* was known to be upregulated in this condition [62], and *ybaY* undergoes supercoiling-dependent transcription associated with the osmotic stress response, which Cheung et al. [63] found to act through RpoS. We also found *yqjE, kbp, ecnB*, and *elaB* to be upregulated. Bulk RNA-Seq data from PRECISE-1K [34] support these observations: in M9 supplemented with 0.3 M NaCl, *yjbJ* ranks seventh by induction across all genes (log_2_fc > 6), and the other genes from this cluster are upregulated as well (*ybaY* : 3.7, *yqjC-yqjE* : 3.1, *kbp*: 2.4, *elaB* : 2.0, *ecnB* : 0.8). Of the 50 most strongly induced genes, 16 are of unknown function.

As shown in Figure 6(A), we identified strong activator binding sites in the predicted promoters for *yjbJ, ybaY* and *yqjE*. The peaks for *yjbJ* and *ybaY* show striking similarity, including 12 of 20 matching bases with a stretch of 7 consecutive matches (Figure 6(B)). Mass spectrometry experiments did not yield a candidate transcription factor, despite using various environments for cell lysate preparation and *in vitro* binding, illustrating the difficulty of this approach when binding is strongly condition dependent.

**Figure 6.**
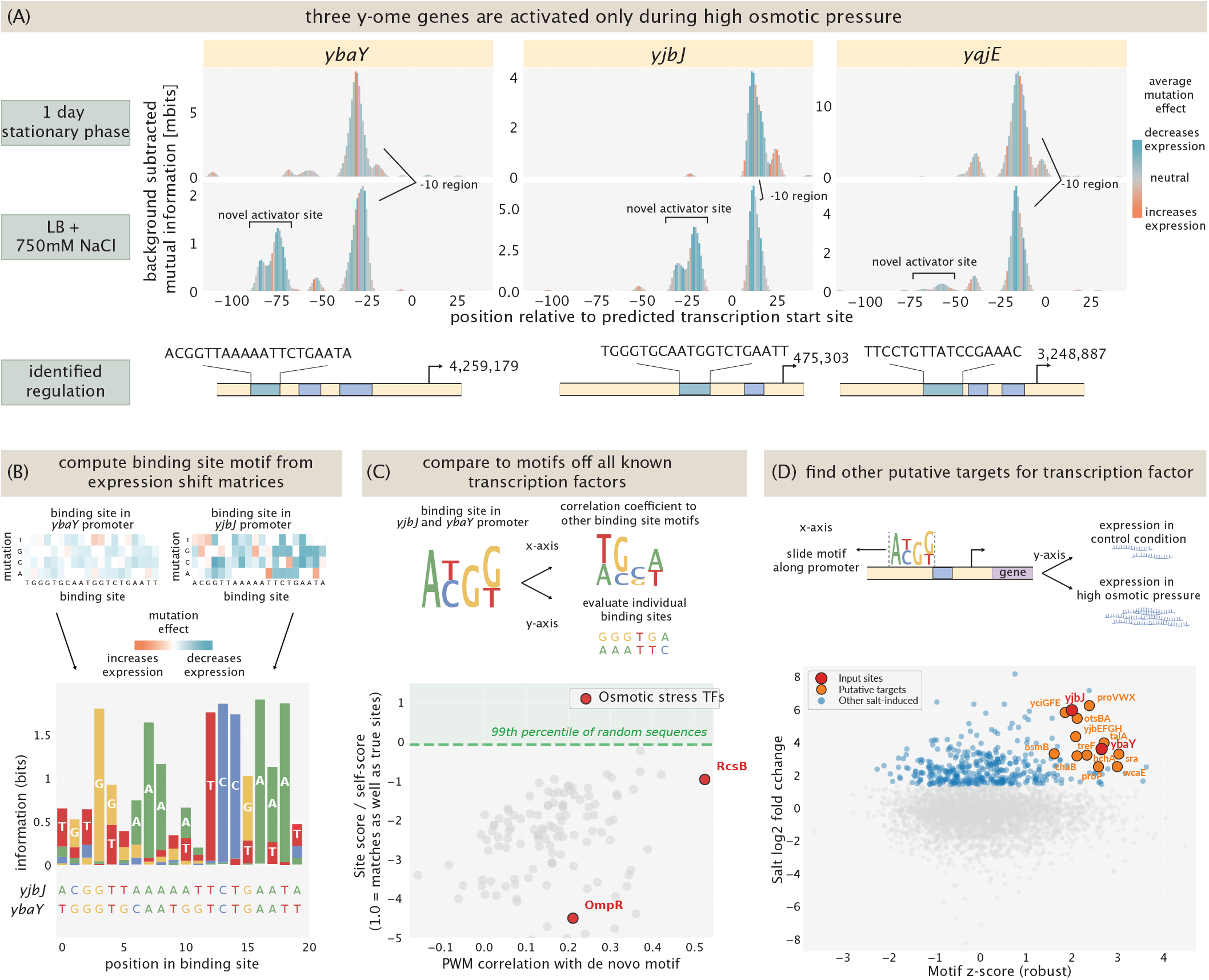
Unknown transcription factor activates y-ome genes under osmotic stress. (A) Information footprints in stationary phase (1 day) and under osmotic shock for *ybaY, yjbJ*, and *yqjE* promoters. An activator peak (blue) appears specifically under osmotic stress. Binding site sequences and transcription start sites for the three y-ome promoters are indicated. (B) De novo motif constructed from expression-weighted Reg-Seq data at the activator binding sites of *yjbJ* and *ybaY*. Letter heights reflect information content; wild-type binding site sequences for all three promoters are shown below. (C) Comparison of the de novo motif against position weight matrices for 145 annotated *E. coli* transcription factors. The x-axis shows the best Pearson correlation; the y-axis shows the normalized site score (1.0 would indicate a match as strong as the transcription factor’s own motif). Red points mark transcription factors known to be active under osmotic stress. The dashed green line indicates the 99th percentile of scores from 1000 random genomic sequences. (D) Promoter motif score (robust z-score) vs. salt-induced gene expression (log_2_ fold change from PRECISE-1K [34]). Red points mark the two input sites used to build the PWM. Orange points mark putative additional targets with both high motif scores and functional connections to osmotic stress. Many of the most salt-induced genes (upper left) have low motif scores, indicating regulation by other systems. *tesB*, which appears divergently transcribed from *ybaY*, scores identically to *ybaY* because the scan detects the same physical binding site from the opposite strand; it is excluded from the candidate list.

As a computational alternative, we constructed a position weight matrix (PWM) from the binding sites in the *yjbJ* and *ybaY* promoters; the *yqjE* site was excluded as less similar with a weaker footprint. Rather than computing the PWM from sequence alignment alone, we use our expression shift matrices to weight each position by the effect of mutations on expression, giving a more direct map from sequence to expression (SI Section S5.1). We compared this de novo motif against PWMs for all 145 transcription factors with at least 3 annotated binding sites in RegulonDB, using two complementary similarity measures: the Pearson correlation between PWMs (maximized over alignment offset and orientation), and a normalized site score in which we evaluate each transcription factor’s known binding sites against our de novo PWM. As shown in Figure 6(C), no transcription factor stands out as a candidate. The identified binding sites are not palindromic, suggesting the binding factor is unlikely to be a homodimer; a heterodimer is one possibility. The right half of our motif matches the canonical right half of an RcsB half-site (TCCTAAAAT) [64], and RcsB scores highest regarding the PWM similarities, but the left half does not match any known RcsB binding partner.

To identify additional candidate targets of this unknown activator, we scanned the 400 bp upstream of every annotated gene in *E. coli* K-12 MG1655 with our de novo PWM (sliding across both strands, retaining the maximum score), and quantified salt-induced expression as the mean log_2_ fold change across all eight pairwise comparisons of four salt and two control replicates in PRECISE-1K. We reasoned that genuine targets should both score high in the motif scan and be salt-upregulated. Neither criterion alone is sufficient: high motif scores occur at promoters of unrelated genes (e.g., *sufB* [65]), and many strongly induced genes are regulated by other known systems. Genes were classified as candidates if they exceeded log_2_fc > 1.5 in PRECISE-1K and had established or plausible roles in osmotic adaptation; their motif scores are shown on the *y*-axis of Figure 6(D), and the full list with scores and fold changes is provided in SI file 2.

Both input sites score well above the genomic background (*ybaY* : *z* = 2.65, *yjbJ* : *z* = 2.00), as expected for sites used in PWM construction. Among the remaining salt-induced genes, the most striking hits are core components of the osmotic stress response. The *proVWX* operon, encoding the high-affinity ABC transporter for glycine betaine and proline [66–68], scores high and is strongly induced; its transcription is osmotically regulated and depends on DNA supercoiling [69], consistent with [63]. The *otsBA* operon, encoding trehalose-6-phosphate synthase and phosphatase [70], and *treF* [62], encoding cytoplasmic trehalase, also score high and are strongly induced. Together these implicate the unknown activator in trehalose homeostasis under osmotic stress, controlling both synthesis and degradation. In each operon, the motif is found upstream of the first gene and internal genes (*proW, otsA*) score near background, consistent with a single binding event driving the entire transcription unit. Additional candidates include *yjbE* (first gene of the *yjbEFGH* exopolysaccharide operon), the H-NS-repressed *yciGFE* operon [62], *talA* (transaldolase, salt-induced at the protein level [62]), and *osmB* (an osmotically inducible outer membrane lipoprotein).

By contrast, several of the most salt-induced genes — *bdm, kdpAB, osmY, ydeO* — score near background, consistent with regulation by other characterized systems (KdpDE [71], RpoS [72], EvgA [73]). This separation suggests the motif identifies a specific regulatory program rather than salt induction generically.

### Transcription Start Sites: Old and New

While *de novo* promoters have been reported as evolving from random sequences [74], we investigated how often this phenomenon occurs based on single mutations in identified promoters. Here, we looked for promoters that had a large increase in expression caused by a single specific mutation, which can be identified from the information footprint by finding single narrow peaks, and expression shifts matrices where only one of the possible three mutations at this position increases expression significantly. As shown in Figure 7(A), three positions in the *σ*^70^ −10 consensus are especially important for binding. Most of the time, mutations creating new transcription start sites move a sequence closer to this consensus sequence. To confirm that the observed mutation increases the binding affinity of *σ*^70^, and leads to a new transcription start site that is distinct from the initially reported or predicted one, we deploy the computational model from LaFleur et al. [42], which predicts a transcription rate from a given promoter sequence, and highlights in which element of the promoter (e.g. −10 region, −35 region, spacer etc.) the mutation resides. We use the model to scan the 160 base pair promoter region for every possible transcription start site, and compare the resulting rates from sequences that carry the mutation of interest to all other sequences. When we find a transcription start site with a strongly increased transcription rate, we identify the transcription start site as a *de novo* promoter.

**Figure 7.**
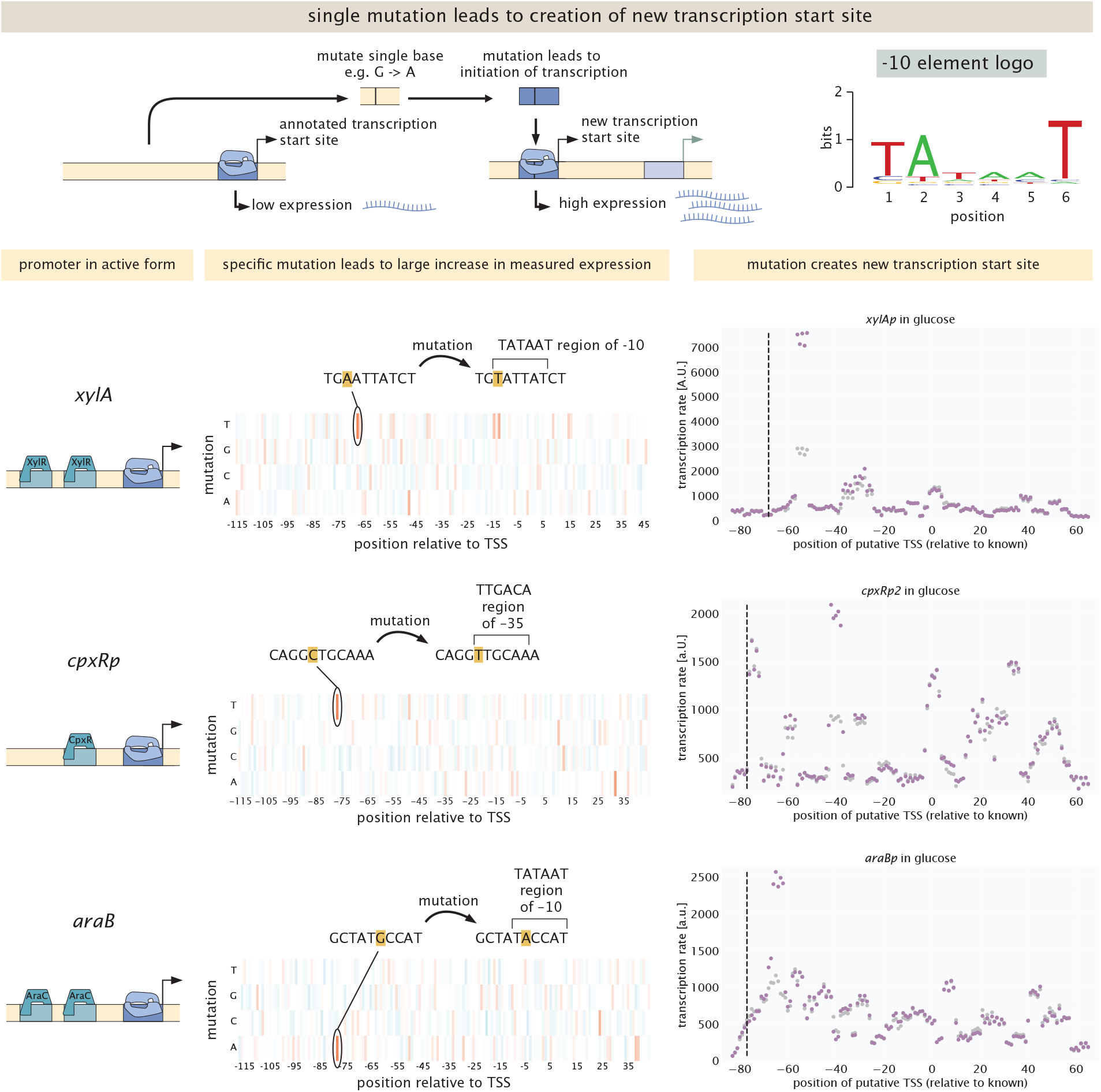
*De novo* promoters. (A) Single mutations away from the annotated transcription start site can lead to a new transcription start site, a *de novo* promoter. (B)*De novo* promoter in the *xylA* promoter, shown here for cells grown in minimal media with glucose as carbon source, a condition where the promoter is not activated. (C)*De novo* promoter in the *cpxR* promoter, shown here for cells grown in minimal media with glucose as carbon source, a condition where the promoter is not activated. (D)*De novo* promoter in the *araBp* promoter, shown here for cells grown in minimal media with glucose as carbon source, a condition where the promoter is not activated.

We find multiple promoters with strong *de novo* promoters (Figure S10), especially promoters that rely on activation by specific activators, such as *xylAp* (activated by XylR in xylose), *cpxRp* (activated by CpxR by e.g. gentamicin) and *araBp* (activated by AraC in arabinose). As shown in Figure 7(B), in the case of *xylAp*, a mutation about 70 base pairs upstream of the annotated transcription start site increases the binding affinity of *σ*^70^ to a different −10 region, creating a new transcription start site which is upstream of the annotated one. The predicted transcription rate at this site is increased two-fold from this single mutation, leading to a measurable level of transcription in most conditions. As shown in Figure 7(C-D), we find similar results for the promoters of *araB* and *cpxR*.

Whether these reflect chance mutations on random sequences [74] or a beneficial feature of activator-dependent promoters poised one mutation away from function remains an open question; the examples in Figure S10 are concentrated at activator-dependent promoters, consistent with the latter, but a genome-wide analysis is needed to settle the question.

While de novo promoters reveal how new transcription start sites can emerge from existing sequence, our experiments also clarify the status of transcription start sites that were already on the books. In several cases, earlier work had hypothesized the existence of multiple distinct transcription start sites at a single promoter. For example, for the *ompR* promoter, as shown in Figure S11(A), inspecting the 5’ ends of mRNAs using primer extension assays [75] revealed multiple distinct mRNA species, suggesting the existence of at least four transcription start sites within a window of 116 bases. We find that only one transcription start site — the one associated with *ompRp1* — is active in any of the 39 conditions tested, as the information footprints exhibit binding peaks only at this site (Figure S11). This is consistent with the multiple 5^′^ ends seen by Huang and Igo arising from post-transcriptional processing rather than independent initiation events, though we cannot exclude that the alternate TSSs fire under conditions absent from our panel.

## 4 Discussion

In this paper, we have tackled the parallel and entangled challenges of discovering regulatory architectures and the environmental conditions that affect them. By using massively parallel reporter assays in conjunction with information theory, we are able to peer into the uncharacterized regions of the *E. coli* regulatory genome, and by carrying out our experiments across a broad array of growth conditions, we get a glimpse of the rich context dependence of those architectures. The version of Reg-Seq used here introduces a streamlined library protocol and genome-integrated reporters via a modified ORBIT method [39], which together make the approach scalable enough to anticipate genome-wide application.

How well does Reg-Seq actually work? Across the 41 assessable promoters, we recover 49 of 103 testable annotations as Reg-Seq footprints. Many missed sites were originally identified *in vitro* or under non-physiological conditions, reflecting the difficulty of harmonizing *in vitro* and *in vivo* methods. Other misses point to real features of the regulatory landscape: transcription factors that bind DNA without a detectable effect on expression, or annotations resting on sequence similarity alone. Beyond verification, our approach led to new environment-dependent regulatory architectures, including a CRP binding site at the y-ome gene *yadI*, repression of *ybiYW* by YciT under aerobic conditions, and a shared activator binding site at *yjbJ* and *ybaY* under osmotic stress.

Some of our gene categories did not yield coherent regulatory pictures. The two iModulons reported by Lamoureux et al. [34] as putatively controlled by YmfT and YgeV, and the six genes chosen as part of feedforward loop motifs, did not produce consistent footprints in the conditions we tested. We treat these as inconclusive rather than negative: a richer condition panel, or direct perturbation of the putative regulators, would be needed to test these predictions against base-pair-resolution footprints. The salt-induced y-genes *yjbJ* and *ybaY* illustrate a different limitation. Despite a clear, reproducible activator footprint, mass spectrometry across multiple lysate preparations failed to identify the responsible transcription factor, and no annotated *E. coli* TF motif matches our de novo PWM. This points to a class of regulators that current chromatography and mass spectrometry workflows cannot reliably recover when binding is strongly condition-dependent.

There is still much work to be done. Increasing the size of the mutated regions in our libraries is mostly limited by the availability and affordability of synthesized oligo pools. A broader range of environmental conditions is needed to conclude what conditions a given gene “cares” about. The transcription factor identification step requires complementary experimental and bioinformatic approaches, and even then is unable to find all transcription factors in all environments; the salt-induced activator above is one concrete example, and developing alternative routes for transcription factor identification — including expanded libraries of purified TFs for direct binding assays, and bioinformatic methods that integrate co-expression with motif scanning — will be essential.

A particularly promising route for transcription factor identification is to combine Reg-Seq with CRISPRi-based knockdown libraries. By measuring Reg-Seq footprints in cells where individual transcription factors have been knocked down via gRNA-targeted dCas9, the loss or weakening of a footprint at a candidate promoter would directly identify the responsible regulator. This is well suited to the cases where mass spectrometry has failed, including the salt-induced activator at *yjbJ* and *ybaY*, since it bypasses the need for direct biochemical capture of the TF. Genome-scale CRISPRi libraries are now well established in *E. coli* [76–78] and can be combined with our barcoded reporter libraries in a single experimental design.

Despite these challenges, the work described here provides a viable route to whole-genome reconstruction of the regulatory landscape of an organism in a single experimental approach and corresponding analytical pipeline.

## 5 Acknowledgements

We are grateful to N. Belliveau, A. Flamholz, H. Garcia, E. Gökman, S. Grill, B. Ireland, B. Jones, F. Jülicher, J. Kinney, G. Salmon, N. Typas, G. Urtecho, V. Vitelli and C. Wiggins for useful discussions, and V. Garcia for help with cell lysis for mass spectrometry experiments. We are grateful to the NIH for support through award numbers DP1OD000217 (Director’s Pioneer Award) and NIH MIRA 1R35 GM118043-01. TR was supported by a fellowship from Boehringer Ingelheim Fonds. This work was supported by Igor A. Antoshechkin and by the Millard and Muriel Jacobs Genetics and Genomics Laboratory at the California Institute of Technology. This research was supported in part by grants from the National Science Foundation (NSF, Grant No. DMS-2235451) and Simons Foundation (Grant No. MP-TMPS-00005320) to the NSF-Simons National Institute for Theory and Mathematics in Biology (NITMB). This project has been made possible in part by the Chan Zuckerberg Initiative DAF (Grant No. DAF2023-329587), an advised fund of the Silicon Valley Community Foundation. This research was supported in part by grants from the NSF (Grant No. PHY1748958) and the Gordon and Betty Moore Foundation (2919.02) to the Kavli Institute for Theoretical Physics (M. M.).

## 6 Data and Code Availability

Sequencing data is available at the SRA database under PRJNA1263894. Mass spectrometry data is available at Caltech Data https://doi.org/10.22002/qtm95-b2j72. Code written to process and analyze data, as well as to generate figures, is available on Github https://github.com/RPGroup-PBoC/reg-seq_env. An interactive dashboard to explore information footprints for every gene in every condition can be found at http://rpdata.caltech.edu/data/interactive_footprints.html.

## Supplemental Information

### S1 Existing methods for dissection of gene regulation in bacteria

A huge effort has been expended in uncovering how genes in *E. coli* are regulated [4, 79–89]. For much of the history of modern molecular biology, genes were usually studied on a one-by-one basis due to the lack of high throughput methods. This led to major success stories including insights into the lysis-lysogeny decision in bacteriophage lambda, discoveries on how bacteria use different carbon sources including lactose, galactose and arabinose, insights into the role of DNA looping and a myriad of other examples. Over the past few decades, a variety of high-throughput methods have re-enlivened the subject by enabling the identification of many binding sites for a single transcription factor in one experiment, or identifying binding sites for all transcription factors without knowing the identity of the transcription factors specifically binding those sites. Here we provide an overview of some examples of previous work to give a sense of where our own efforts fit into this enormous subject, highlighting both the successes and open questions resulting from previous work. We begin by discussing in vitro approaches in which DNA and proteins are interrogated outside of their natural cellular environments. Much has been learned from these approaches. Then, we turn to the analysis of in vivo methods which attempt to capture DNA-protein interactions in the context of living cells.

- SELEX: Systematic evolution of ligands by exponential enrichment (SELEX) was developed in 1990 [90] with the purpose of identifying *in vitro* which DNA sequence or ligand a protein binds to. The method uses a library of synthesized DNA that is incubated with purified proteins. Unbound DNA is removed and bound DNA is eluted from the protein and subsequently amplified. This process is repeated over multiple rounds to find DNA with high binding affinity to the protein. The level of specificity of the DNA obtained in these experiments can be tuned by choosing a different number of cycles. Once a DNA sequence with high affinity for the protein is identified, the genome of interest can be scanned for potential transcription factor binding sites by looking for sequence similarity between the genomic DNA and the SELEX DNA. In the context of *E. coli*, binding sites for hundreds of transcription factors have been identified genome wide [91].
- PBM: Protein-binding microarrays (PBM) use a large array of synthesized DNA oligonucleotides that are fused to a surface. Binding of purified protein to the DNA oligonucleotides is measured by fluorescence microscopy of tagged proteins. PBMs are able to better detect less specific binding sites for TFs than SELEX [92]. PBMs have been used to identify the motifs for more than 1000 transcription factors [93].
- DAP-Seq: One modification to *in vitro* binding assays that has been useful is to choose genomic DNA as template instead of synthesized DNA. This approach is the basis of DNA affinity purification sequencing (DAP-Seq) [94, 95]. One of the advantages of using genomic DNA is that such DNA maintains chemical modifications to the DNA such as methylation and reveals that such methylation can be important for DNA - TF interactions.
- ChIP-Seq: Chromatin immunoprecipitation with sequencing (ChIP-Seq) is one of the most commonly used methods to identify binding sites for TFs in the *in vivo* setting. Identifying binding sites *in vivo* is beneficial as potential co-factors and enzymes modifying the conformation of the TF are present if the correct growth condition is chosen. In the first iterations of the method the resolution of identified binding sites was low, however, the use of endonucleases in ChIPexo-Seq [96] lead to higher accuracy. To pull down TFs that are crosslinked to DNA, the TF needs to be modified with a tag, e.g., His-tag [97], or antibodies against the TF need to be available, which can limit the throughput and often means that only one or a few TFs can be studied at the same time. However, binding sites across the entire genome can be found in a single experiment, giving high throughput on this axis. There have been drastic differences in the number of binding sites identified for certain TFs between ChIP-Seq and SELEX [97]. ChIP-Seq has been used to discover a variety of different DNA-Protein interactions, such as the RpoS regulon in *E. coli* [97], the PhoB regulon [40], nucleoid organization by H-NS and MukBEF [98], genome wide binding of CRP (using DNA microarrays) [99] and binding of Sigma70 [100]. In a recent study, ChIp-Seq was performed on 139 TFs for cells grown in minimal media with glycerol [12].
- DNAse footprinting: In contrast to ChIP-Seq, DNAse footprinting does not require a pull down on a specific TF. This allows for the discovery of binding for all DNA binding proteins across the entire genome at the same time, but also comes with the loss of the identity of the protein binding to each site. This method has been combined with RNAP occupancy studies to verify the function of identified binding sites [11, 101].
- MPRA: Massively-parallel reporter assays are one of the signature recent achievements in highthroughput approaches for dissecting promoter function. The method is based upon creating large libraries of genetic variants and measuring their function in parallel. Urtecho et al. use genome-integrated massively parallel reporter assays to catalog and characterize promoters throughout the *E. coli* genome [60]. This approach has been used impressively in *E. coli* to explore not only binding sites, but also ribosomal binding site sequences [102], etc.
- Sort-Seq: In Sort-Seq [6, 18] binding of transcription factors is identified by mutating bases in the vicinity of a transcription start site and measuring expression of a downstream reporter gene using fluorescence activated cell sorting (FACS), followed by DNA sequencing of the mutated promoter variants. By identifying bases where a mutation leads to a large change in expression putative binding sites are discovered. The identity of the transcription factor binding to these sites is then identified by DNA chromotography and mass spectrometry. This approach not only identifies binding of transcription factors, but also shows that the binding is functional, i.e., binding of the transcription factor effects expression of a gene. It has been found that binding sites that have been identified in vitro do not necessarily imply that the binding has regulatory function [40].

All of these methods have been used to gain insights into the regulatory landscape of *E. coli* and other organisms. Databases such as EcoCyc [103] and RegulonDB [104] contain a large number of annotated binding sites. Our goal is to use data from these methods as well as approaches in our paper to provide a systematic, rigerous and complete description of all promoters in *E. coli* with standardized annotations in databases that can be used for e.g. phylogenetic modeling and building blocks for synthetic biology. In particular, as shown in several of the figures in the paper, in those cases where we are able to find putative binding sites and identify the TFs that bind to those sites, the aim is to be able to go from a promoter of unknown regulatory architecture all the way to an environment-dependent regulatory architecture including energy matrices describing transcription factor binding and statistical mechanical models of the input-output response of the promoter of interest as a function of key regulatory knobs such as DNA-TF binding site strength, TF copy number and effector concentration. Beyond that, the aim is for all of these disparate sources to come together to make excellent databases such as EcoCyc [103] and RegulonDB [104] a more complete and internally consistent source as a basis for rigorous understanding of the physiology and evolution of *E. coli* and that will serve as a template for how to structure such databases for other organisms.

### S2 Experimental Procedures

For the 104 genes in this study, we found 119 promoters (some genes are part of operons with multiple annotated transcription start sites and promoters), leading to 178,619 promoter variants that were ordered and more than 95% (170,167) that were recovered during mapping. Across all promoter variants, 5,316,504 unique barcodes were identified, with a median of 28 barcodes per variant. Because we previously observed that expression from plasmids depends significantly on plasmid copy number, possibly leading to non-physiological expression levels [105], we used genome-integrated versions of our libraries. Following genome integration, 168,952 promoter variants and 2,232,542 barcodes were found, with a median of 13 barcodes per promoter variant. Having multiple unique barcodes per variant is essential for handling possible biases that could be introduced by different sequences during gene expression or library preparation. The entire library was then grown in 39 unique growth conditions, as described in the Methods. Once the cultures reached the desired state, they were harvested and prepared for DNA and RNA sequencing to count abundances of barcodes. DNA barcode counts reveal the number of cells harboring a given promoter mutant; mRNA barcode counts reveal the level of gene expression for that given promoter mutant.

#### S2.1 Oligo Pool Design

##### S2.1.1 Identification of Transcription Start Sites

For each gene chosen for this study, shown in Table S5, we first looked for its promoter on EcoCyc [103]. If the promoter was found, the annotated transcription start site (TSS) was used. If multiple promoters were identified, each promoter was included in the experiment. If no promoter was found, we looked for transcriptionally active sites in the data set from Urtecho et al.[60]. In their work, the genome was fragmented and every fragment was tested for transcription initiation in LB. If we could find a site that was identified as active close to the gene of interest, we chose this site as the origin for computational promoter mutagenesis. If no TSS could be identified for a gene, the model from LaFleur et al. [42] was used to computationally predict a TSS in the intergenic region. The site predicted to be the most active within 500 bp upstream of the coding region was chosen as the TSS. Initially, 119 promoters were chosen, however, 7 promoters (mglBp, hdeAp2, mtnp, ybeDp, cpxRp2, galEp1, and ompFp), had an identical TSS as another promoter annotated in Ecocyc. The duplicated promoters were treated as independent when mutated variants were created, leading to twice the number of variants in the total pool.

##### S2.1.2 Computational Promoter Mutagenesis

Once a TSS is identified, the 160 bp region from 115 bp upstream of the TSS to 45 bp downstream is taken from the genome. Most of the cis regulation has been shown to occur within this window. Based on the approach of Kinney et al. [18], each promoter sequence is randomly mutated at a rate of 0.1 per position. 1500 mutated sequences are created per promoter, following the approach of [8], which creates sufficient mutational coverage in the promoter region. The promoter oligonucleotides are flanked by restriction enzyme sites (SpeI at 5’ and ApaI at 3’) which are used in subsequent cloning steps. The restriction sites are flanked by primer sites that are used to amplify the oligo pool. The primer sequences were chosen from a list of orthogonal primer pairs, designed to be optimal for cloning procedures [106]. Oligo pools were synthesized (TwistBioscience, San Francisco, CA, USA) and used for subsequent cloning steps.

#### S2.2 Library Cloning

##### S2.2.1 Cloning Oligo Pool into a Plasmid Vector

The oligo pool was amplified using a 20 bp forward primer (SC142) and a 40 bp reverse primer (SC143), which consists of a 20 bp primer binding site and a 20 bp overhang. PCRs were run with minimal amplification until faint bands appear on an agarose gel in order to minimize amplification bias, using 10 ng of the oligo pool as template, as recommended by TWIST. The PCR was run for with 10 amplification cycles using a volume of 12.5 μl. PCR products were cleaned and concentrated (DNA Clean & Concentrator-5, ZymoResearch) and used for a second amplification step. The 20 bp overhang on the reverse primer from the first amplification was used as primer site for a reverse primer (SC172), which contains randomized 20 bp barcode, flanked by two restriction enzyme sites (SbfI and SalI, 5’ to 3’ direction). The forward primer is the same as in the first amplification step (SC142). PCR amplification is run again with minimal amplification to minimize amplification bias, which we found to be 8 cycles. PCR products are run on a 2% agarose TAE gel and subsequently extracted and purified (Zymoclean Gel DNA Recovery Kit, ZymoResearch). In the next step, a restriction digest is performed on the outer restriction enzyme sites (SpeI-HF and SalI-HF). Unless noted otherwise, all restriction digests were run for 15 minutes at 37 °C, using 1 μg of DNA as template and using 1 μl of each restriction enzyme. The plasmid vector was digested with different restriction enzymes which create compatible sticky ends (XbaI and XhoI). Most restriction enzyme sites are palindromes, so by choosing different enzymes with compatible ends, we avoid having palindromes flanking the plasmid inserts after ligation. The oligo pool is combined with the plasmid vector using T7 DNA ligase (New England Biolabs, Ipswich, MA, USA) following the supplier’s protocol. Ligation products were cleaned and concentrated (DNA Clean & Concentrator-5, ZymoResearch) and a drop dialysis (MF-Millipore VSWP02500, MilliporeSigma, Burlington, MA, USA) was performed for 1 hour to improve sample purity. Electroporation using *E. coli* pir116 electrocompetent cells (Lucigen, Middleton, WI) was performed at 1.8 kV in 1 mm electroporation cuvettes, followed by 1 hour recovery at 37 °C and 250 rpm in 1 ml LB media (BD Difco). The entire culture was plated on 150 mm LB + kanamycin (50 *μ*g/ml) petri dishes and grown overnight at 37 °C. The following day, plates were scraped and the colonies resuspended. Freezer stocks were prepared using a 1:1 dilution of resuspended colonies and 50% glycerol. Cultures were inoculated with 5 × 10^8^ cells in 200 ml of LB + kanamycin (50 *μ*g/ml) and grown at 37 °C until saturation. Plasmids were extracted (ZymoPURE II Plasmid Maxiprep Kit, ZymoResearch) and used for subsequent sequencing. For details, see Section S2.3. The plasmid library is then used as template in a restriction digest using restriction enzymes ApaI and SbfI-HF. The resulting product was cleaned and concentrated (NEB Monarch) and the DNA concentration was measured on a NanoDrop. Similarly, the RiboJ::GFP element was PCR amplified (primers SC191 and SC192), adding restriction sites as overhangs (ApaI and PstI). For details about the restriction sites, see Table S1. The PCR product was cleaned and concentrated (NEB Monarch) and digested with the respective restriction enzymes. The plasmid library is combined with the RiboJ::sfGFP element using T7 DNA ligase (New England Biolabs, Ipswich, MA, USA) following the supplier’s protocol. Ligation products were cleaned and concentrated (NEB Monarch) and a drop dialysis (MF-Millipore VSWP02500, MilliporeSigma, Burlington, MA, USA) was performed for 1 hour to improve sample purity. Electroporation using *E. coli* pir116 electrocompetent cells (Lucigen, Middleton, WI) was performed at 1.8 kV in 1 mm electroporation cuvettes, followed by 1 hour recovery at 37 °C and 250 rpm in 1 ml LB media. Entire cultures were plated on 150 mm kanamycin (50 *μ*g/ml) + LB petri dishes aweren grown overnight. The following day, plates were scraped and the colonies resuspended. Freezer stocks were prepared using a 1:1 dilution of resuspended colonies and 50% glycerol. Cultures were inoculated with 5 × 10^8^ cells in 200 ml of LB + kanamycin (50 *μ*g/ml) and grown at 37 °C until saturation. Plasmids were extracted (ZymoPURE II Plasmid Maxiprep Kit, ZymoResearch) and used for subsequent genome integrations.

**Table S1.** Restriction sites used. All enzymes were ordered from NEB. If available, high fidelity versions of the enzymes were used.

| Part s | 5' restriction site | 3' restriction site |
| --- | --- | --- |
| Plasmid Vector | XbaI | XhoI |
| RiboJ::GFP | ApaI | PtsI |
| Oligo Pool | SpeI | ApaI |
| Barcoding Primer | SbfI | SalI |

During post-hoc inspection of the integrated reporter, we identified a single-base mutation in the GFP coding sequence that introduces a premature stop codon at codon 4. The mutation was present in the template used for all library constructs, so every reporter in the library carries it. We did not use GFP fluorescence as a readout in this work; Reg-Seq quantifies transcription from the mRNA-to-DNA barcode count ratio, which does not require a functional reporter protein. The mutation is, however, expected to affect translation and therefore potentially mRNA half-life through ribosome occupancy effects. Because the GFP cassette is identical across all constructs, any such effect produces a uniform offset in absolute mRNA abundance that cancels out in the cross-construct (information footprints, expression shift matrices) and cross-condition comparisons that underlie our analyses.

#### S2.3 Barcode Mapping

The plasmid library is used for barcode mapping. Purified plasmid is PCR amplified using forward primer (SC185) outside the promoter region and a reverse primer outside the 20 bp barcode (SC184). The PCR is run with minimal amplification, and the product is gel purified (NEB Monarch). The purified DNA was used as template for a second PCR using a primer (SC196), adding an Illumina P5 adapter to the promoter side, using a primer (SC199), and adding an Illumina P7 adapter to the barcode side. The PCR is again run with minimal amplification and gel purified (NEB Monarch). The product was used for sequencing on a Illumina NextSeq 2000 with a P2 flow cell with pair-end reads using primers SC185 for read 1, SC184 for read 2 and SC201 for the index read. Reads were filtered and merged using custom bash scripts, which are available in the Github repository. After processing, each promoter/barcode pair was identified in each read, and pairs with less than 3 total reads were discarded. An alignment algorithm was used to identify the identity of each sequenced promoter variant. This allowed us to include additional promoter variants that were in the initial oligo pool because of synthesis errors in the production of the oligos. The barcode mapping was used in the analysis of libraries grown in various growth conditions. The code used to perform processing of sequencing data can be found in the associated Github repository. Processing is done with the help of various software modules [107–109]. Custom Python code used for the analysis and visualization of results can be found in the associated Github repository.

An important filtering step is to remove any promoter variant-barcode pairs with less than three reads to remove potential sequencing errors that escaped quality control. In total, we obtained 376655 unique sequences, which is more than double the sequences we had initially designed (178619). However, more than 53% of the identified sequences are associated with only one barcode. If we filter for sequences that are associated with at least 2 barcodes, we are left with 172418 unique sequences, which contain more than 96.5% of our initial pool, highlighting that we were able to maintain the library throughout the first cloning steps. Figure S1 shows how many unique variants were identified per promoter variant. Most promoters are close to the expected number of variants, with some notable exceptions, especially the promoter for *dicC*, which has the lowest number of recovered variants. Here, we looked at the mutation rate across all recovered variants and found an increase around the transcription start site, as shown in Figure S2. The area where the increase in mutation rate is observed coincides with the location of a binding site for DicA. The working hypothesis is that binding of DicA to the plasmid containing its binding sites comes with a significant fitness cost to the cells, probably due to the titration effect [105]. Thus, since mutations are randomly spaced throughout the promoter sequence, only variants with more mutations in the binding site survive, as binding of DicA is reduced.

#### S2.4 Genome Integration

##### S2.4.1 Creation of Landing Pad Strain

We used ORBIT [39] to integrate reporter libraries into the *E. coli* chromosome. ORBIT uses a targeting oligo containing an attB site, and an integration plasmid using an attP site. An additional helper plasmid facilitates the integration of the targeting oligo into the replication fork, followed by recombination of the attB and attP sites catalyzed by a *bxb-1* gene in the helper plasmid. To increase the efficiency of genome integration, we created a landing pad strain that contains an attB site close to the *glmS* gene in the *E. coli* chromosome. Wild type *E. coli* (K12 MG1655) is streaked on a LB plate and grown overnight at 37 °C. A single colony is picked and prepared to make elecotrocompetent cells as follows. The picked colony is grown in 3 ml of LB at 37 °C and shaken at 250 rpm overnight. The overnight culture is diluted 1:1000 into fresh LB (e.g. 200 ml) and grown at 37 °C and 250 rpm until exponential phase, reaching an optical density at 600nm (OD600) of ~ 0.4. The cultures are then immediately put on ice and spun in a centrifuge at 5000 g for 10 min. Following the spin, the supernatant is discarded, and the cells are resuspended in the same volume as the initial culture of deionized water at 4 °C. The cells are spun again at 5000 g for 10 min. This wash step is repeated 3 times with 10% glycerol. After the last wash step,the supernatant is discarded and cells are resuspended in the remaining liquid and distributed into 50 *μ*l aliquots. Aliquots are frozen on dry ice and kept at −80 °C until they are used for electroporation. For electroporation, aliquots are thawed on ice and 1 mm electroporation cuvettes are pre-chilled on ice. 100 ng of helper plasmid (Addgene #205291) is added to a 50 *μ*l cell aliquot and mixed by slowly pipetting up and down. The aliquot is then added to the electroporation cuvette and electroporation is performed at 1.8 kV. The aliquot is recovered with 1 ml of LB media pre-warmed to 37 °C. The culture is recovered for 1 hour at 37 °C and shaken at 250 rpm. After recovery, aliquots at various dilutions are plated on LB + gentamicin (gentamicin sulfate 15 μg/ml). Plates are grown overnight and a single colony is picked to prepare electrocompetent cells and frozen stocks as described above. The cells are electroporated with 2 mM of the targeting oligo (SC219) and an integration plasmid containing *kanR* for selection and the *sacB* gene for counterselection. After recovery, the cultures are plated on LB + s (50 *μ*g/ml). A colony is picked and electrocompetent cells are prepared again as mentioned above. Another electroporation is performed using only the targeting oligo (SC219). This time, cells are plated on LB + 7.5% sucrose for selection of loss of the integrated cassette, leaving only an attB site in the locus. This results in a scarless insertion of the attB site into the chromosome.

##### S2.4.2 Integration of the Library

To perform genome integration, the host strain carrying the helper plasmid is made electrocompetent (follow growing and washing steps described above), and the plasmid library is electroporated into the host strain, using about 100 ng of plasmid per transformation. The cells are recovered in 3 ml of prewarmed LB + 1% arabinose and shaken at 37 °C at 250 rpm for 1 hour. The entire volume is plated on LB + kanamycin plates and colonies are grown over night. The next day, all colonies are scraped, resuspended in LB and diluted to an OD600 of 1. The helper plasmid used for genome integration causes growth deficits, hence, the library needs to be removed of the plasmid. Therefore, the library is inoculated with 0.5 ml of culture at an OD600 of 1 in 200 ml of LB, and grown until exponential phase at 37 °C shaken at 250 rpm. The helper plasmid carries the *sacB* gene, which is used for negative selection in the presence of sucrose. At exponential phase, the culture is plated on LB + 7.5% sucrose agarose plates. Plates are grown overnight, scraped and made into frozen stocks at an OD600 of 1. The frozen stocks are then ready for growth experiments.

#### S2.5 Growth Media and Culture Growth

##### S2.5.1 Base Media

Lysogeny Broth (LB) was prepared from powder (BD Difco, tryptone 10 g/l, yeast extract 5 g/l, sodium chloride 10 g/l), and sterilized by autoclaving. M9 Minimal Media pre-mix without carbon source was prepared in the following way, similar to [32]: to 700 ml of ultrapure water, 200 ml of 5 × base salt solution (BD Difco, containing disodium phosphate (anhydrous) 33.9 g/l, monopotassium phosphate 15 g/l, sodium chloride 2.5 g/l, ammonium chloride 5 g/l, in H_2_O, autoclaved), 10 ml of 100X trace elements (5 g/l EDTA, 0.83 g/l FeCl_3_-6H_2_O, 84 mg/l ZnCl_2_, 19 mg/l CuSO_4_ − 5 H_2_O, 10 mg/l CoCl_2_ − 6H_2_O in H_2_O, 10 mg/l H_3_BO_3_, 1.6 mg/l MnCl_2_ − 4H_2_O, prepared as described in [110]), 1 ml 0.1 M CaCl_2_ solution, in H_2_O, autoclaved, 1 ml 1 M MgSO_4_ solution, in H_2_O, autoclaved and 1 ml of 1000 × thiamine solution (1 mg/ml in water, filter sterilized) were added. The resulting solution was filled up to 1 l with water and filter sterilized. M9 minimal medium was complemented with carbon source by mixing appropriate amounts of carbon-source-free M9 minimal medium and carbon source stock solutions. Carbon source stock solutions were prepared as 20% solutions and filter sterilized.

##### S2.5.2 Cultivation

Overnight cultures were incubated from frozen stock in 200 ml M9 minimal media with 0.5% glucose and grown at 37 °C while shaken at 250 rpm. Cultures were diluted 1:100 into the respective growth media (prewarmed to 37 °C, 200 ml) and grown to exponential phase (OD600 of 0.3). To ensure steady state growth, the cultures were diluted a second time 1:100 into the same growth media and grown again to an OD600 of 0.3, ensuring at least 10 cell divisions in the growth media. At this step, there are four different paths for a culture: 1. It is immediately harvested (called standard growth).; 2. A compound is added to the culture and the culture is harvested at a later specified time (called induction); 3. the culture is moved to water bath of a different temperature and then harvested at a later specified time (called cold or heat shock); or 4. the culture is spun down in four 50 ml aliquots at 3500 rpm for 7 min, washed in a different media twice, and then grown in that media for 1 hour (called shock). Unless otherwise mentioned, glucose was used as carbon source. Each condition was done in duplicate with some conditions being done in triplicate when the initial replicates did not seem to correlate well. To compare how experiments correlate, for each pair of conditions, we computed the Pearson correlation coefficient across all mutual information footprints.

##### S2.5.3 Specific Growth Conditions

###### Glucose

For standard growth, 5 ml of 20% glucose solution added to 200 ml of M9 minimal media pre-mix for a final concentration of 0.5%.

###### Arabinose

For standard growth, 5 ml of 20% arabinose solution added to 200 ml of M9 minimal media pre-mix for a final concentration of 0.5%.

###### Xylose

For standard growth, 5 ml of 20% xylose solution added to 200 ml of M9 minimal media pre-mix for a final concentration of 0.5%.

###### Galactose

For standard growth, 2.3 ml of 20% galactose solution added to 200 ml of M9 minimal media pre-mix for a final concentration of 0.23%.

###### Acetate

For standard growth, 5 ml of 20% sodium acetate solution added to 200 ml of M9 minimal media pre-mix for a final concentration of 0.5%.

###### LB

For standard growth, cultures were grown in LB media.

###### Nitrate

For standard growth, potassium nitrate was added to M9-glucose media for a final concentration of 80 mM.

###### Anaerobic

For anaerobic growth, M9-glucose media was kept in a glove box containing nitrogen for multiple days to equilibrate and remove oxygen from the media. Cultures were inoculated in 20 ml of this media inside the glove box in glass tubes which are sealed with rubber plugs. Tubes were grown in a shaker at 37 °C and shaken at 250 rpm. When the culture reached an OD600 of 0.3, a 1:100 dilution was performed inside the glove box and grown to an OD600 of 0.3 again. Cultures were then harvested.

###### Anaerobic and Nitrate

Potassium nitrate was added to standard M9-glucose media for a final concentration of 80 mM. Cultures were grown in anaerobic conditions as described above.

###### Stationary Phase

Cultures were grown in M9-glucose media for an additional one day (1d) or three days (3d) after reaching an OD600 of 0.3 under standard growth conditions.

###### Sodium Salicylate

1 M sodium salicylate stock was prepared and filter sterilized. For standard growth, 500 μl of the stock was added to 200 ml of M9-glucose media for a final concentration of 2.5 mM. For 1 hour induction, 2 ml of the stock was added to 200 ml M9-glucose media for a final concentration of 10 mM.

###### Ethanol

For standard growth, 5 ml of 200 proof ethanol was added to 200ml of M9-glucose media for a final concentration of 2.5%. For 1 hour induction, 10 ml of 200 proof ethanol was added to 200 ml M9-glucose media for a final concentration of 5%.

###### Ampicillin

For both standard growth and 1 hour induction, ampicillin was added to M9-glucose media for a final concentration of 2 mg/l.

###### Leucine

For 1 hour induction, leucine was added to M9-glucose media for a final concentration of 10 mM.

###### Phenazine Methosulfate

For 1 hour induction, 61 mg of phenazine methosulfate (SigmaAldrich) was added to M9-glucose media for a final concentration of 100 μM.

###### 2,2 Dipyridyl

For 1 hour induction, 156 mg of 2,2 dipyridyl (SigmaAldrich) was added to M9-glucose media for a final concentration of 5 mM.

###### Gentamicin

For 1 hour induction, gentamicin was added to M9-glucose media for a final concentration of 5 mg/l.

###### Copper Sulfate

1M stock of CuSO_4_ was prepared. 1 hour inductions were performed with final concentrations of both 500 μM and 2 mM.

###### Hypochlorous Acid

For 1 hour induction, sodium hypochlorite solution (Sigma-Aldrich #425044) was added to M9-glucose media for a final concentration of 4 mM.

###### Spermidine

For a 1 hour induction, spermidine was added to M9-glucose media for a final concentration of 5 mM.

###### Serine Hydroxamate

For a 30 min induction, serine hydroxamate (Sigma-Aldrich) was added to M9-glucose media for a final concentration of 0.4 mg/ml.

###### H_2_O_2_

For 30 min induction, H_2_O_2_ was added to M9-glucose media for a final concentration of 0.1 mM. For 10 min induction, H_2_O_2_ was added to M9-glucose media for a final concentration of 2.5 mM. **Cold Shock**: Cultures were grown in M9-glucose media to an OD600 of 0.3 and then were immersed in a 10 °C water bath and shaken for 1 hour.

###### Medium Cold Shock

Cultures were grown in M9-glucose media to an OD600 of 0.3 and then were immersed in a 19 °C water bath and shaken for 1 hour.

###### Heat shock

Cultures were grown in M9-glucose media to an OD600 of 0.3 and then were immersed in a 42 °C water bath and shaken for 5 min.

###### Nitrogen Starvation

Minimal media premix was prepared with only 10% NH_4_Cl. Cultures were grown in M9-glucose media to an OD600 of 0.3, then washed and grown for 1 hour in M9-glucose media with reduced NH_4_Cl.

###### Magnesium Starvation

Minimal media premix was prepared where MgSO_4_ was replaced with NaSO_4_ at the same concentration. Cultures were grown in M9-glucose media to an OD600 of 0.3, then washed and grown for 1 hour in M9-glucose media with NaSO_4_.

###### Sulphur Starvation

Minimal media premix was prepared where MgSO_4_ was replaced with MgCl at the same concentration. Cultures were grown in M9-glucose media to an OD600 of 0.3, then washed and grown for 1 hour in M9-glucose media without MgSO_4_.

###### pH2

Minimal media was prepared as usual, but the pH is adjusted to 2.0 using sodium hydroxide. Cultures were grown in M9-glucose media to an OD600 of 0.3, then washed and grown for 1 hour in M9-glucose media with pH 2.0.

###### Glutamic Acid pH 2.5

Minimal media was prepared with 1 mM of glutamic acid at pH 2.5. Cultures were grown in M9-glucose media to an OD600 of 0.3, then washed and grown for 1 hour in M9-glucose media with glutamic acid at pH 2.5.

###### Low Phosphate

Minimal media was prepared with only 10 % of disodium phosphate and monopotassium phosphate. Cultures were grown in M9-glucose media to an OD600 of 0.3, then washed and grown for 1 hour in low phosphate M9-glucose media.

###### Low Osmolarity

Minimal media pre-mix was diluted by a factor of two before adding glucose. Cultures were grown in M9-glucose media to an OD600 of 0.3, then washed and grown for 1 hour in low osmolarity M9-glucose media.

###### High Osmolarity

LB was supplemented with 0.75 M NaCl. Cultures were grown in M9-glucose media to an OD600 of 0.3, then washed and grown for 1 hour in LB with 0.75 M NaCl.

#### S2.6 Barcode Sequencing

Once a culture is ready for harvesting, 750 μl of culture are mixed with 750 μl of freshly prepared 1X Monarch DNA/RNA Protection Reagent (NEB) and pelleted by spinning at 20000 g for 1 minute. The supernatant is discarded, and the pellets are frozen on dry ice. Genomic DNA is extracted from four pellets for each sample using a Monarch Spin gDNA Extraction Kit (NEB), following the manufacturer’s protocol for gram-negative bacteria. RNA was extracted and reverse transcription was performed using a custom protocol. 500 ng of gDNA and 5 μl of cDNA was used as template for library preparation. First, the template is amplified by PCR using primers SC184 and SC88. 12 cycles are run for gDNA and between 20 and 25 cycles for cDNA, depending on the sample. The PCR product was run on a 2% agarose gel and bands were gel purified (Monarch DNA Gel Extraction Kit, New England Biolabs). Then, 5 ng of amplified DNA was used for a second PCR (50 μl volume), using forward primer SC80 and one of 92 reverse primers (SC354-SC445), which add an index for demultiplexing. The PCR is run for 6 cycles and the product is run in a 2% agarose gel, followed by gel extraction. The extracted DNA is used for sequencing. Sequencing runs were performed on a NextSeq 2000. A summary of all the sequencing runs used for this paper is shown in Table S2. Primer SC450 was used for read 1, and primer SC270 for the index read. Sequencing data is filtered for quality and trimmed using *fastp* [108]. Barcodes are extracted and counted from sequencing files using custom Bash scripts, which are available on Github.

**Table S2.** Sequencing runs. Every sequencing run containing data used in this work. Each run has its own code for processing, which can be found in the associated Github repository.

| Date | Content | Run Type | Sequencer |
| --- | --- | --- | --- |
| 05/14/2022 | Mapping of promoter variants and barcodes | Paired-End | Next-Seq 2000 - P2 flowcell |
| 03/27/2023 | Comparison of plasmid reporters and genome integrated reporters | Single-end | MiSeq v2 flowcell |
| 09/07/2023 | Barcode counting in 9 Conditions | Single-end, 50 cycles | MiSeq, v2 flowcell |
| 12/07/2023 | Barcode counting in 27 Conditions | Single-end, 27 cycles | Next-Seq 2000, P3 flowcell |
| 06/21/2024 | Barcode counting in 44 Conditions | Single-end, 27 cycles | Next-Seq 2000, P4 flowcell |
| 09/07/2024 | Barcode counting in 24 Conditions | Single-end, 27 cycles | Next-Seq 2000, P2 flowcell |

#### S2.7 DNA Chromatography and Tandem Mass Spectrometry

##### S2.7.1 Cultivation for Lysate

Similar to the procedure described in S2.5.2, overnight cultures were grown in 5 ml of M9 minimal medium with 0.5% glucose at 37 °C and then diluted 1:100 into the growth media listed in S2.5.3. Cultures were carried out in 2800 ml Fernbach-style flasks filled with 500 ml of media. The total volume of liquid culture for a given growth condition ranged from 1000 to 6000 ml, depending on the number of required DNA chromatography experiments. After a given growth condition duration is completed, the cells were harvested by centrifuging at 8000 g for 30 minutes at 4 °C. Cell pellets were stored at −80 °C or subsequently lysed.

##### S2.7.2 Lysate Preparation

Cell pellets were re-suspended in lysis buffer (70 mM potassium acetate, 50 mM HEPES pH 7.5, 5 mM magnesium acetate, 2.5 mM DTT, and cOmplete Ultra EDTA-free protease inhibitor). Mechanical cell lysis was performed using a high pressure cell disruptor (Constant Systems). Afterwards, to help solubilize membrane proteins, n-dodecyl-*β*-D-Maltoside (DDM) detergent was added to the crude lysate for a final concentration of 1 mg/ml. Lysates were clarified of non-soluble cell debris by centrifuging at 30000 g for 1 hour at 4 °C. The collected supernatants were further concentrated to ~ 100 mg/ml using centrifugal protein concentration filter (Amicon Ultra −15) with a molecular weight cut-off of 3 kDA. Protein concentrations were determined using a fluorometer (Qubit fluorometer) and proprietary dyes that specifically label proteins (Qubit Reagent). The lysates were further cleared of non-specific DNA binding proteins by incubating with a competitor salmon sperm DNA (Invitrogen) at 0.1 mg/ml for 10 minutes at 4 °C. An additional 1 hour incubation at 4 °C is performed by adding sacrificial streptavidin-coated magnetic beads without any attached DNA oligos (Dynabeads MyOne Streptavidin T1) at ~ 3 mg/ml in order to clear proteins that may non-specifically bind to the beads surfaces. Lysates are centrifuged one final time to pellet the sacrificial beads and any remaining insoluble component. The resulting supernatants are stored at −80 °C or aliquoted into volumes of 200 μl for subsequent DNA chromatography experiments.

##### S2.7.3 DNA Chromatography

DNA affinity chromatography is used to isolate a transcription factor of interest from a given cell lysate. The procedure detailed below is similar to the one we have used previously [6, 111]. In brief, DNA oligos that have putative transcription factor binding sites are attached to magnetic beads. These beads with tethered DNA are incubated with cell lysates to “fish out” proteins that bind to the oligos. They are spatially separated from the remaining lysate by magnets, allowing for extraction of the bound proteins.

For each protein detected by mass spectrometry, we quantify specific binding through an enrichment value computed as follows. Within each pulldown sample, we first compute the relative abundance of every protein as its mass-spec counts divided by the total counts across all detected proteins; this normalization removes per-run differences in total peptide load. The enrichment of a given protein at a target bait is then the ratio of its relative abundance in the target pulldown to its relative abundance in a control pulldown run in parallel from the same lysate, where the control bait is a sequence near the *ymjF* promoter with no Reg-Seq footprint (Section 2.S2.7.3.1). An enrichment value of 1 indicates that the protein occupies the same fraction of the recovered proteome on the target as on the control, i.e., no specific binding. Values substantially greater than 1 indicate specific binding to the target sequence.

###### S2.7.3.1 DNA Oligos for Magentic Beads

The binding sequence of an oligo (IDT) is taken from the native *E. coli* genome, where the sequence region is hand-selected to match the putative binding sites determined by RegSeq. For the control sequence, a region near the TSS associated with the promoter ymjF was used, since this sequence had no discernible binding sites, as determined by RegSeq. Each oligo has the 5’ end biotinylated (to ensure attachment to streptavidin-coated magnetic beads) and starts with a cut site sequence for the PstI restriction enzyme (New England Biolabs), which allows for the bound protein to be recovered by a restriction digest.

###### S2.7.3.2 Bead Incubations and Protein Recovery

A batch volume of magnetic beads (Dynabeads MyOne Streptavidin T1) is measured out, according to the total number of DNA chromatography experiments being performed and assuming each individual chromatography experiment requires 160 μl of stock beads per 200 μl of aliquoted lysate. The total volume of beads is washed twice in TE buffer (10 mM Tris-HCl pH 8.0, 1 mM EDTA), washed twice in DW buffer (20 mM Tris-HCl pH 8.0, 2 M NaCl, 0.5 mM EDTA), and re-suspended in annealing buffer (20 mM Tris-HCl pH 8.0, 10 mM MgCl2, 100 mM KCl). The beads are aliquoted according to the number of oligos used. DNA oligos are added to the aliquoted beads to a final concentration of 5 μM and incubated for at least 3 hours at room temperature or overnight at 4 °C. After oligo incubation, beads are washed twice in TE buffer and then twice DW buffer. All wash buffers are supplemented 0.05% TWEEN-20 detergent to minimize bead loss related to sticking to surfaces. After washing, beads are incubated in a blocking buffer (20 mM HEPES pH 7.9, 300 mM KCl, 0.05 mg/ml bovine serum albumin, 0.05 mg/ml glycogen, 2.5 mM DTT, 5 mg/ml polyvinylpyrrolidone, and 0.02% DDM) for 1 hour at room temperature to reduce nonspecific protein binding to the bead surfaces. The beads are then washed three times in lysis buffer. The beads are added to the aliquoted lysates to a final concentration of 5 mg/mL. When applicable, a supplement of the reagent defining a given growth condition (carbon source, antibiotic, chemical stress,…) is added to the lysate to approximate the internal cell environment. For example, for the M9-glucose growth condition, glucose is added to the lysate to a final concentration of 0.5%. See Table S7 for details of all lysate supplements used. The beads and lysate are incubated overnight at 4 °C on a rotating rack. The next day, the beads are washed three times in lysis buffer and once in the reaction buffer (NEB buffer r3.1) for the restriction enzyme PstI. 1000 units of PstI is added to each bead reaction. The beads are incubated for 90 min at room temperature. The supernatant containing the DNA and bound proteins is collected for solution-based protein digestion.

##### S2.7.4 Protein Digestion, Labeling and Desalting for Proteomic Analysis

The samples were subjected to an isobaric-labeled filter-aided sample preparation (iFASP) protocol (PMID: 23692318) with minor changes. Briefly, supernatant from each sample was loaded onto a 10 kDa Amicon filter (Pierce), and washed with 8 M urea in 100 mM HEPES (urea buffer) 3 times. Each washing step includes adding 200 μl of the corresponding solution followed by 14000 g centrifugation for 15 min. After 3 washes with urea buffer, 200 μl of urea buffer containing 5 mM tris(2-carboxyethyl)phosphine was added into each filter to break disulfide bonds. The reaction was incubated for 1 hour at room temperature, and 200 μl of urea buffer containing 20 mM of chloroacetamide was added into each filter to alkylate free thiols. The alkylation reaction was incubated for 15 min at room temperature, and the filters were centrifuged for 14000 g for 15 min. The filters were further washed 3 times with 150 μl of 100 mM of triethylamine bicarbonate (TEAB) in water. After the TEAB washes, 120 μl of 100 mM TEAB containing 1 μg of Trypsin (Pierce) was added into each filter. The enzyme to substrate ratio is estimated be from 1:5 to 1:10. The trypsinization step occurred for 16 hours at 37 °C. After trypsinization, 5 μl of DMSO containing 0.05 mg of TMTpro reagent (Thermo) was added into each filter, and the labeling reaction incubated for 1 hour. 1 μl of 5% hydroxylamine was added into each filter to quench the TMT labeling. The samples were then eluted from the filters by 14000 g centrifugation for 15 min. The filters were further washed 3 times with 50 μl of 0.5 M NaCl in water, and all elutions were pooled together. The pooled sample was dried using a CentriVap concentrator (LabConco), and was desalted with a monospin C18 column (GL Science) according to manufacturer’s instructions. The desalted sample was dried again using a CentriVap concentrator and was stored at −80 °C.

##### S2.7.5 Liquid Chromatography and Tandem Mass Spectrometry

Samples were reconstituted in 20 μl of 2% acetonitrile and 0.2% formic acid in water. The peptide concentration was determined using the Pierce Colorimetric Quantitative Peptide Assay. An aliquot of 500 μg of the peptide was loaded onto a Thermo Vanquish Neo liquid chromatography (LC) system, where the peptides were separated on an Aurora UHPLC Column (25 cm × 75 μm, 1.6 μm C18, AUR2-25075C18A, Ion Opticks). The LC gradient was increased from 2% to 98% of mobile phase B over 130 min. See Table S8 for the gradient settings. The LC processed peptides were analyzed on a Thermo Eclipse Tribrid mass spectrometer using a data-dependent acquisition (DDA) method, where the mass spectrometer selects the most intense peptide precursor ions from the first scan of tandem mass spectrometry (MS1) and then fragments and analyzes the precursors in a second scan (MS2). MS1 scans were acquired with a range of 375–1600 m/z in the Orbitrap at a resolution of 120000. The maximum injection time was 50 ms, and the AGC target was 250. MS2 scans were acquired using the quadrupole isolation mode and the higher-energy collisional dissociation (HCD) activation type in the Iontrap. For these scans, the resolution was 50000, the isolation window was 0.7 m/z and the collision energy was 35%. See Table S9 for the detailed parameters used for the mass spectrometry scans. Peptide identification was performed in Proteome Discoverer 2.5 using the Sequest HT search engine; see Table S10 for the search parameters used.

**Figure S1.**
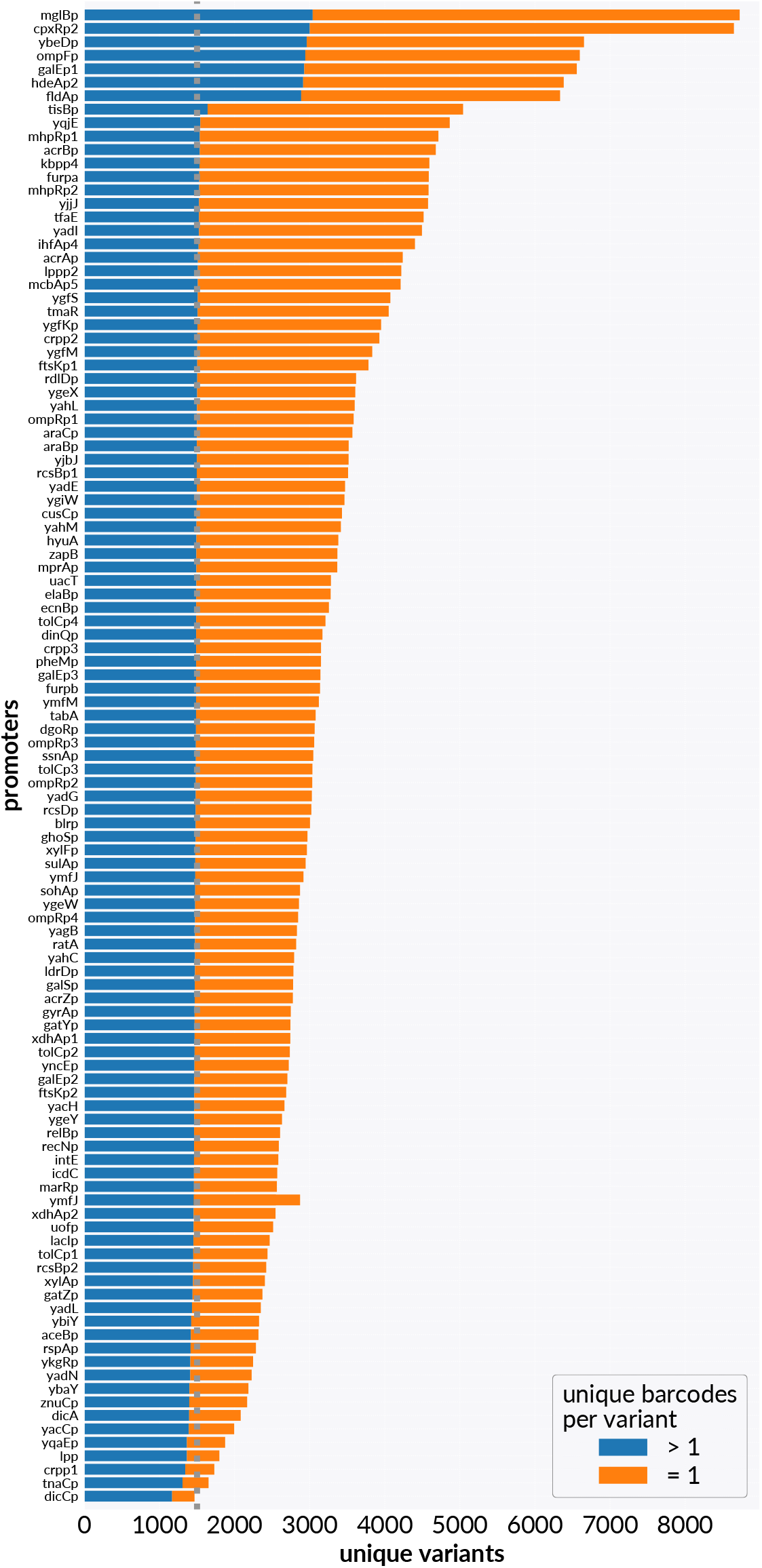
Identified variants for each promoter. Number of unique identified variants of each promoter are shown. Number of variants with one barcode are shown in orange and number of variants with more than one barcode are shown in blue. Number of expected variants (1500) is shown as gray dotted line.

**Figure S2.**
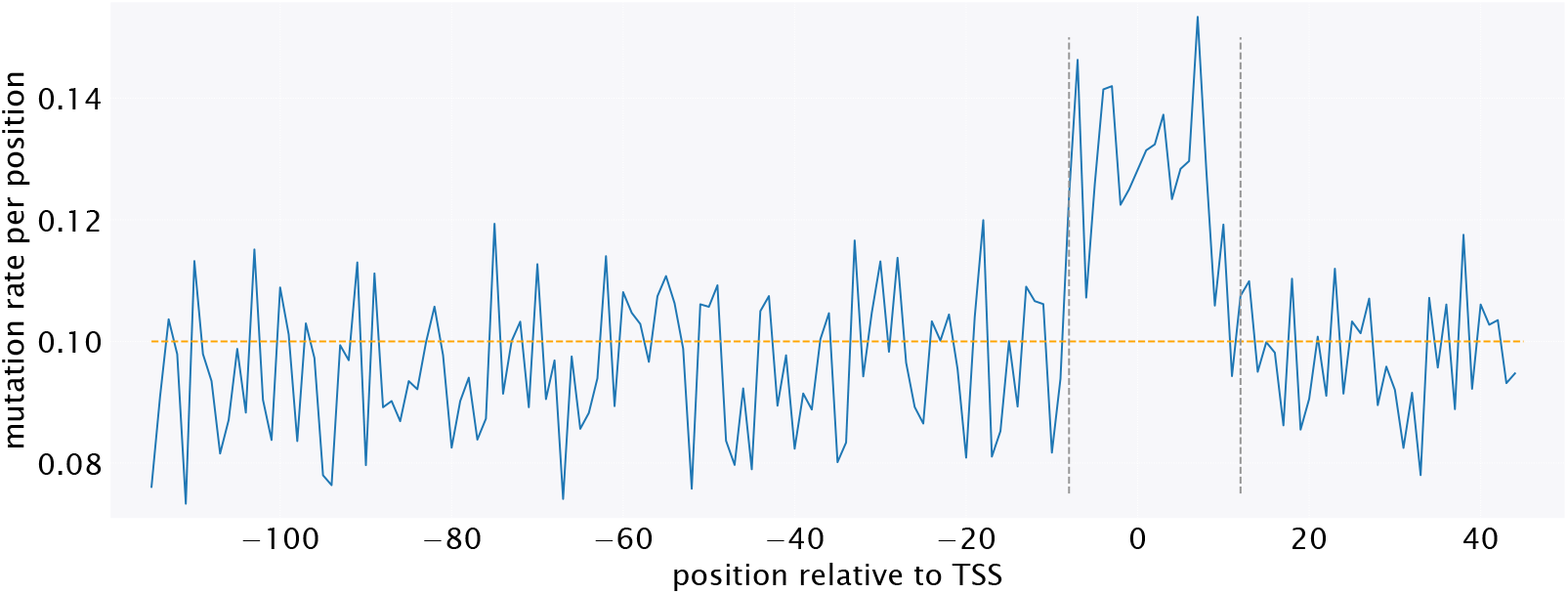
Mutation rate in sequences promoter variants for the *dicCp* promoter. Mutation rate per position across all sequenced promoter variants during promoter-barcode mapping. Sequences are designed to have an average mutation rate of 0.1 per position (orange line). An annotated binding site for DicA is between −8 and +12 (boxed by gray dashed vertical lines.

### S3 What Sets the Scale? Information-Theoretic Foundations of Reg-Seq Analysis

The high throughput nature of Reg-Seq experiments entails a big data processing challenge. Each promoter tested in each condition presents a unique result that needs to be processed and analyzed separately and rigorously. For this purpose we developed a new pipeline to automatically identify binding sites from Reg-Seq data. In this section, we will explore the summary statistics and data processing we use to evaluate and make multidimensional discoveries from Reg-Seq experiments. First, we lay the foundation by taking a deeper dive into mutual information. We organize the discussion around the question of what sets the scale of mutual information one expects to measure in Reg-Seq experiments. What does it mean to get 10^−3^*bits*? Second, we discuss another complementary summary statistic which are expression shift matrices, which give us an even better resolution, as we measure how much expression is changed for each possible mutation at each position.

#### S3.1 Mutual information as a tool to identify binding sites

A proven tool to display the data and show correlations between mutations and their effect on expression from the promoter is the mutual information concept [8, 18]. Mutual information is a metric for characterizing the correlation between two random variables, which measures how much the Shannon entropy of one random variable is reduced by measuring the other,

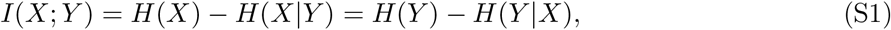

where *H*(*X*) is the entropy of random variable *X*, also called the marginal entropy, and *H*(*X* | *Y*) is the conditional entropy of *X* when *Y* is known. From this equation, we can immediately learn the boundaries of mutual information. Its lower bound is 0, which is the case for two independent variables. Then, no information about *X* is gained by knowing *Y*, and *H*(*X* | *Y*) = *H*(*X*). Since information cannot be lost by measuring one variable, we also know that *H*(*X* | *Y*) ≤ *H*(*X*), thus, mutual information cannot be negative. The upper bound for mutual information is achieved when measuring one random variable determines the other. In this case, the maximum entropy gain is given by the minimum of the marginal entropies of each random variable, max *I*(*X*; *Y*) = min {*H*(*X*), *H*(*Y*)}.

##### S3.1.1 Gene expression toy model

We consider several toy examples to build intuition for how mutual information arises in Reg-Seq experiments. In the first example, we are assuming there is a promoter consisting of a single base, which can be either mutated or wild type. There is a single regulator that can bind to this site, and if the regulator is bound, there is expression from the promoter. We denote the binding energy of the regulator to the wild type sequence to be *h*_*w*_, and the binding energy to the mutated site to be *h*_*m*_. The probability of the regulator binding is then given by

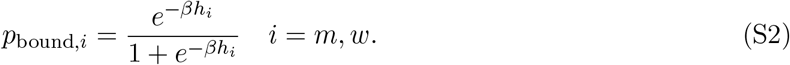

We introduce two random variables. First, the occupancy state *ω*, which can be either bound, *ω* = 1, or unbound, *ω* = 0. Second, the state of the sequence *σ*, which can be either wild type, *σ* = 0, or mutated, *σ* = 1. As shown in Figure S3, at negative binding energies, the probability of finding the base occupied is high, and then decreasing as *h*_*i*_ becomes more positive. Binding energies are given in units of *k*_*B*_*T* and *β* = 1/*k*_*B*_*T*. What does mutual information tell us in this context? It tells us how much we learn about the expression state, is it on or off, by identifying the state of the sequence. Later we will see how we can use this information to identify where binding sites are. For now, in our toy example, we can compute the marginal entropy *H*(*ω*) and the conditional entropy *H*(*ω*|*σ*), which then allows us to compute mutual information in equation (S1). To compute the marginal entropy, we first need to compute the marginal distribution *P*(*ω*). In equation (S2), we have identified the conditional probability of the regulator binding given the state of the sequence, so now we only have to multiply by the probability *P*(*σ*) of finding the base in that state (wild type or mutated). These probabilities are defined by the rate of mutations, which we call *P*(*σ* = 1) = *p*_mut_, and therefore *P*(*σ* = 0) = 1 − *p*_mut_. Together, the marginal probabilities become

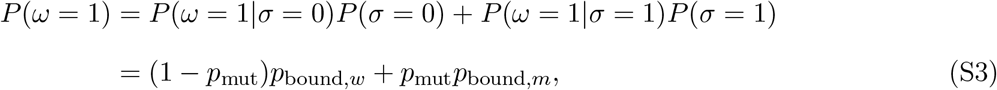

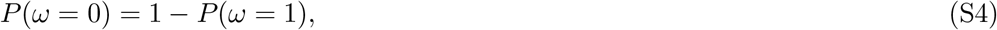

where we can see that finding the site in the bound state is a weighted average of the probability of binding in either the wild type or mutated state, multiplied by the probability of being in such state.

**Figure S3.**
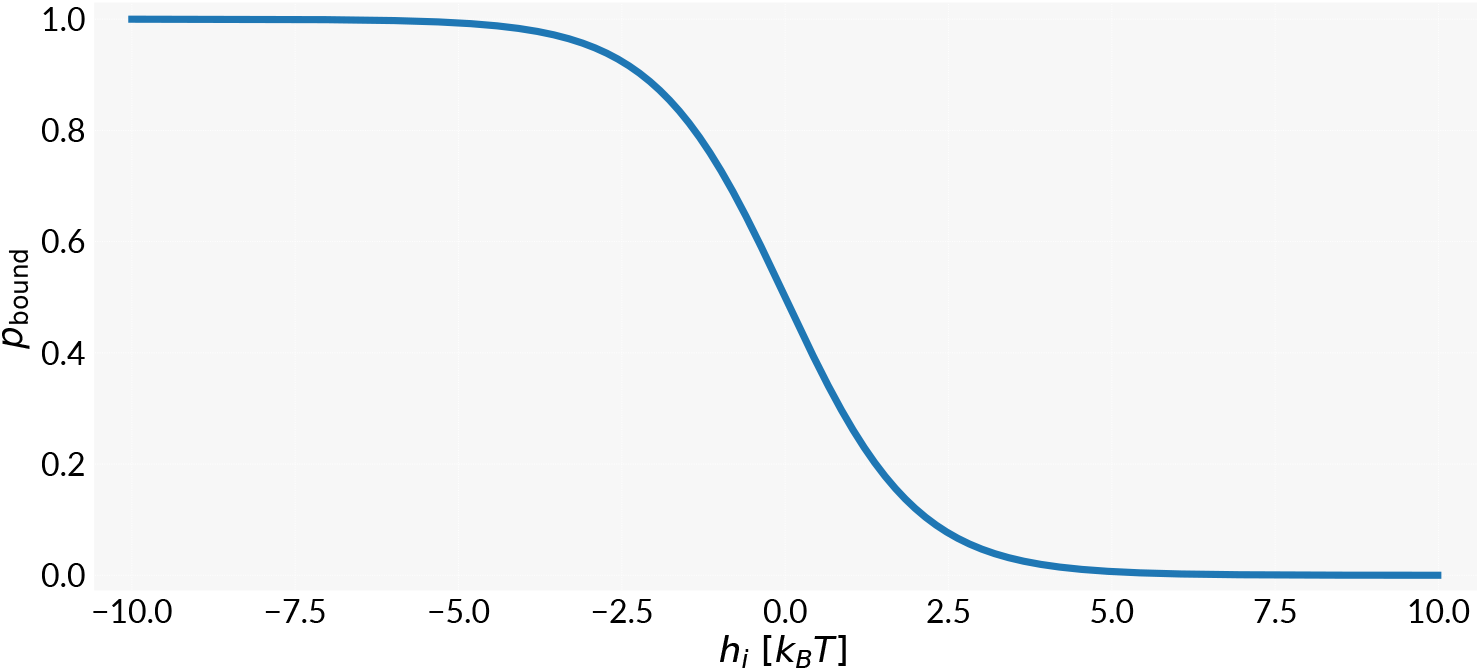
Binding probability in the toy example. Plot of equation (S2) for changes in binding affinity.

To compute the marginal entropy, we compute the Shannon entropy using the two state entropy,

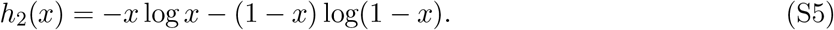

Here we used the fact that if there are only two states, there is only one free variable. If the probability of state 1 is given by *x*, then the probability of state 2 is 1 − *x*, therefore, we can write the two state entropy as a function of *x* only. Using the probabilities from equation (S3-S4), we can compute the marginal entropy *H*(*ω*) as two state entropy,

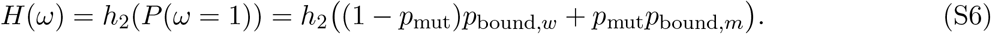

The marginal entropy is thus the entropy of a mixture of the two probabilities *p*_bound,*w*_ and *p*_bound,*m*_, weighted by the probability of getting a mutation or wild type base.

To calculate mutual information, we also need to calculate the conditional entropy *H*(*ω*|*σ*), which measures how much uncertainty about the occupancy state *ω* is remaining after knowing the sequence state,

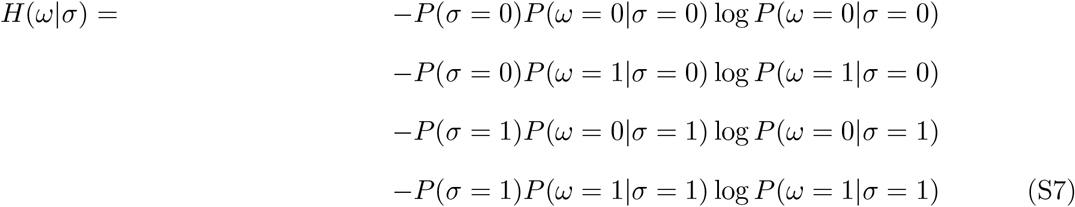

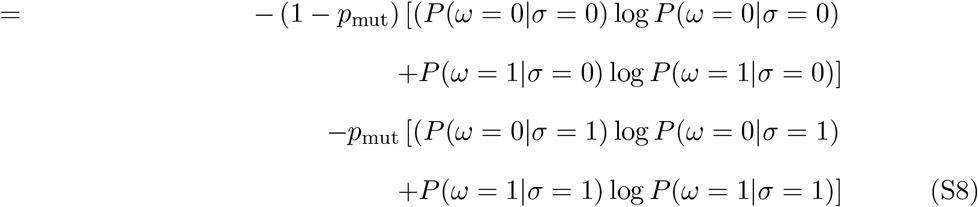

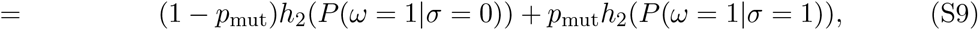

where we used that *P*(*ω* = 0|*σ* = 0) = 1 − *P*(*ω* = 1|*σ* = 0) and *P*(*ω* = 0|*σ* = 1) = 1 − *P*(*ω* = 1|*σ* = 1), and then used the two state entropy equation (S5). Then, the conditional entropy reduces to a mixture of the entropy of the wild type and mutated site each,

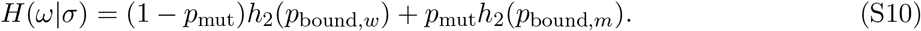

Now we are equipped to compute mutual information from equation (S1),

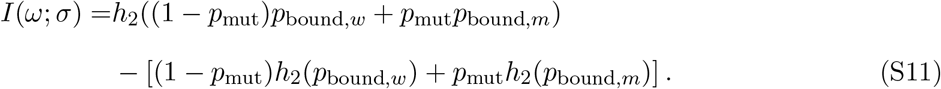

Here, mutual information reduces to the difference between entropy of the mixture and mixture of entropy.

Figure S4 shows the marginal entropy *H*(*ω*) (equation (S6)), the conditional entropy (equation (S10)) and mutual information (equation (S11)) for multiple sets of parameters for binding energies and mutation rates. It is often useful to express the binding energy of the regulator to the mutant sequence as a difference in binding energy to the wild type Δ*h* = *h*_*m*_ − *h*_*w*_, as in practice, it is often easier to infer differences in binding energy. In the case of strong binding to the wild type site (*h*_*w*_ = −10), the difference in binding energies has to be large for there to be a measurable difference in binding probability, as shown in Figure S4AB. However, if the difference of binding energies is high and the gene is always off in the mutated state, then mutual information is maximized, as shown in Figure S4. For weaker binding to the wild type sequence, smaller changes in binding energy lead to measurable differences in expression and therefore, mutual information, as shown in Figure S4CD. The upper limit of mutual information also depends on the mutation rate. We find the maximum mutual information generally decreases with lower mutation rates, as shown in Figure S4BD. An interesting observation is that the conditional entropy has a local maximum at Δ*h* = − *h*_*w*_. In that case the binding probability for the mutant becomes *p*_bound,*m*_ = 1/2, which maximizes its two state entropy, and thus, conditional entropy.

**Figure S4.**
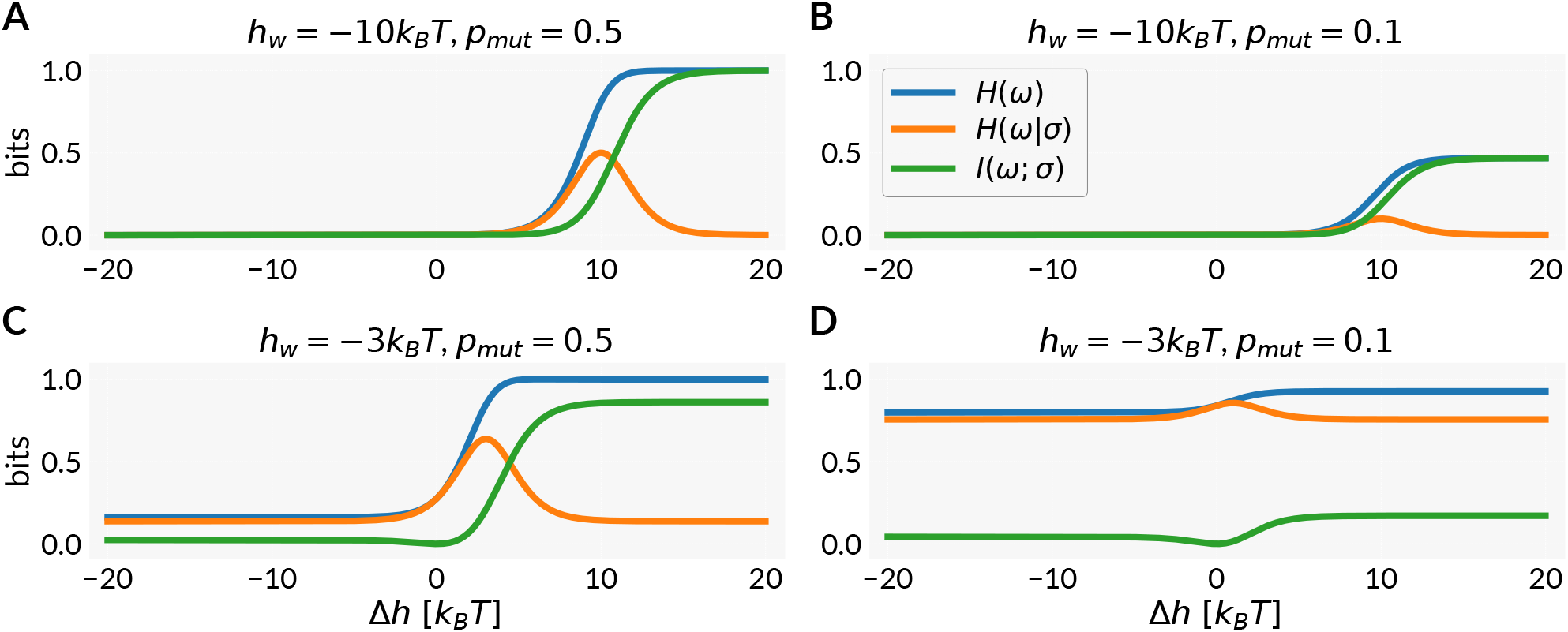
Entropy and mutual information in the one site toy example. Mutual information between binding state *ω* and sequence state *σ* (green) compared to the marginal entropy *H*(*ω*) (blue) and conditional entropy *H*(*ω*|*σ*) (orange), shown as functions of the difference in binding energies Δ*h* for fixed parameters *h*_*w*_ and *p*_mut_.

This example illustrates the constraints on designing experiments to measure differences in binding probability across mutant binding sites. While the example above was computed for a single site, we extrapolate to larger binding sites. If we are dealing with a strong binding site, single possibly small effect mutations are not going to lead to any measurable difference in binding probabilities, hence, one needs to accumulate multiple mutations to perturb binding of the regulator enough to get a measurable difference. Here, we considered a single site. This leads us to our next toy example, where we explore longer binding sites and multiple mutations.

##### S3.1.2 Coin flip toy example

In the next toy example, we consider a series of coin flips. Say, we are performing *L* total flips, with probability *p* of getting heads. This is conceptually equivalent to the mutagenesis scheme in Reg-Seq experiments, where at each base we perform a coin flip if the base gets mutated or stays wild type. Again we are going to compute mutual information, where in this case the random variables are the result of the first coin flip, *X*, and the second variable is the total number of heads *M* in the sequence of *L* flips. We start with equation (S1), which means we compute the entropy for the distribution of coin flip outcomes. We know that the probability of getting *M* heads out of *L* flips is given by a binomial distribution,

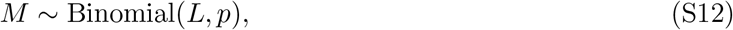

thus the entropy *H*(*M*) is given by the entropy of a binomial distribution,

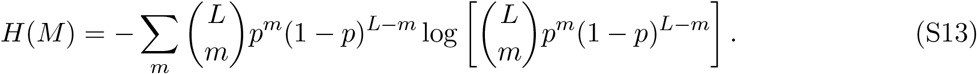

There is no simple way to perform the sum, but we can try approximating the binomial distribution by a Gaussian distribution, which would allow us to compute the entropy analytically. The Gaussian approximation of a Binomial has mean *μ* = *Lp* and variance *Lp*(1 − *p*). Then we can write the Gaussian approximation *P*_*G*_ as

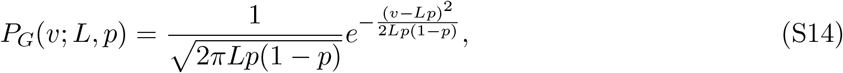

where we replaced the number of heads *m* with a continuous approximation *v*. To compute the differential entropy (differential because *v* is continuous) of this distribution, we have to compute the integral over the continuous variable *v*,

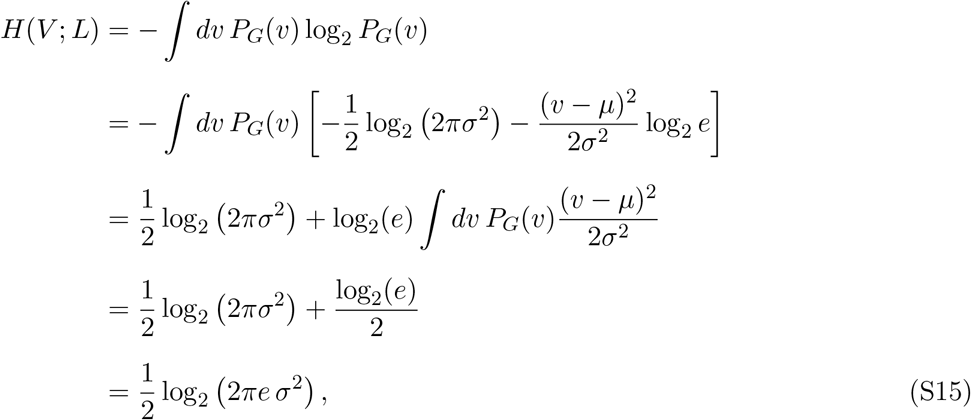

where *μ* = *Lp, σ*^2^ = *Lp*(1 − *p*), and we used *∫ dv P*_*G*_(*v*)(*v* − *μ*)^2^/*σ*^2^ = 1. Substituting back,

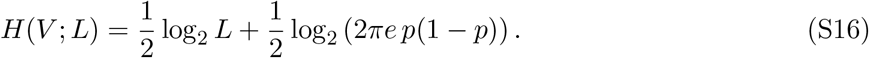

We can see the entropy scales with the log of the length of the site. Next we consider the conditional entropy, *H*(*M*|*X*). Here, we know the outcome of the first flip, and then ask what is the entropy for the set of possible outcomes for the remaining flips. This is again another binomial distribution, but with *L* − 1 flips, as we have already performed one flip. Then, the conditional entropy can be expressed as the entropy of a binomial distribution with *L* − 1 flips, similarly to the marginal entropy in equation (S13). Computing mutual information from equation (S1) is thus equivalent to computing the loss in entropy by going from a binomial of *L* flips to a binomial with *L* − 1 flips. This entropy loss scales with the number of flips, as well as the probability of getting heads. As shown in Figure S5, the mutual information *I*(*M*; *X*) decays with the total number of flips, and the rate of decay scales with the bias of the coin. For a small number of coin flips, the mutual information is higher for a fair coin *p* = 0.5 than for very biased coin *p* = 0.1. Intuitively, that makes sense, as for the biased coin, getting mostly or only tails (if *p* is the probability of getting heads) is very likely, and measuring one coin flip is not giving us much information about the remaining outcomes. But as we perform more coin flips, at some point mutual information becomes larger for the biased coin. When considering the biased coin with *p* = 0.1 and *L* = 20, we get *I*(*M*; *X*) ≈ 0.041 bits. If we used the Gaussian approximation, we would compute mutual information as the difference of entropy of two Gaussian distributions,

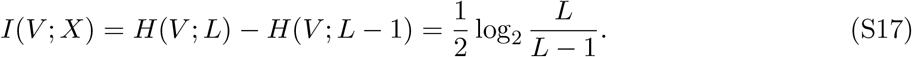

**Figure S5.**
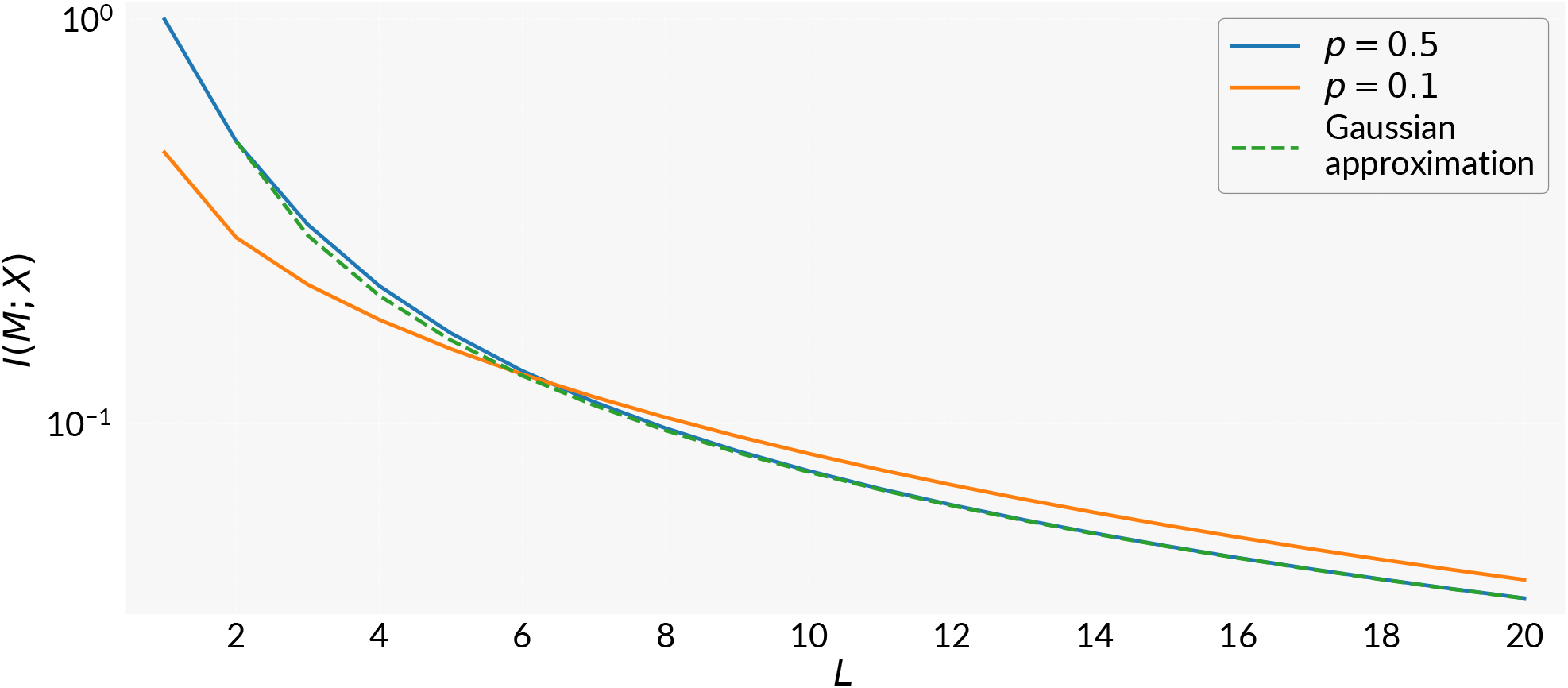
Mutual information in the coin flip toy example. Mutual information between the result of an individual coin flip *X* and the total number of heads *M* for *L* flips with probability *p* of getting heads.

Note that this result is independent of the mutation rate, and most closely resembles the case of *p* = 0.5, as shown in Figure S5. For *L* = 20 we get *I*(*V*; *X*) ≈ 0.037 bits, which is very close to our answer using binomial distributions.

This toy example provides a theoretical upper bound for mutual information in this scenario. As will become clear in the following section, this upper bound applies to Reg-Seq experiments as well.

##### S3.1.3 Mutual information in Reg-Seq experiments

The coin-flip example is directly relevant to the way mutations are introduced in Reg-Seq. Mutations are randomly spaced throughout the sequence, just like coin flips. In our first toy example, we have discussed how a mutation in a sequence can affect the binding energy of a regulator and how it influences its binding probability. Suppose that binding of the regulator leads to a measurable level of gene expression *Y*,

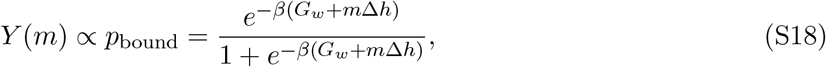

where *m* is the number of mutations in the binding site, each associated with a binding energy cost Δ*h*, and *G*_*w*_ is the free energy of binding. We switch to free energy at this time to account for the fact that there could be multiple non-specific binding sites *N*_*NS*_ in the system that can compete with binding of *R* molecules, *G*_*w*_ = *h*_*w*_ − 1/*β* log(*R*/*N*_*NS*_), where *h*_*w*_ is now the difference in binding between the non-specific background sites and our target binding site. In this case, equation (S18) is the expression for gene expression from a constitutive promoter, where the binding “regulator” in this context is RNA polymerase.

We now ask how much can be learned about the observed gene expression by knowing whether a specific base in the binding site is mutated. Then, there are three variables, the identity of one specific base *X*, the total number of mutations in the sequence *M*, and the observed gene expression *Y*. These three variables can be written in a Markov chain

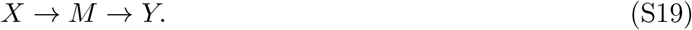

An important concept from information theory is the data processing inequality, which notes that mutual information cannot increase through the Markov chain described in equation (S19),

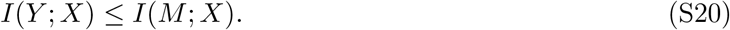

Here we can see how the mutual information between gene expression *Y* and the identity of one specific base is bound from above by the second toy example, just by the random nature of mutations in the sequence. And this is true for any functional form of equation (S18). In the following, we will introduce various steps to the pipeline and therefore the Markov chain, but whatever we do, *I*(*M*; *X*) sets the upper bound for mutual information in our model.

The next step we include is a measurement of gene expression, which leads to an observed gene expression *μ*. Here, we assume there is a lower bound for the observable amount of gene expression we can measure *Y*_min_. If the true expression is below this limit, we will still observe it as *Y*_min_,

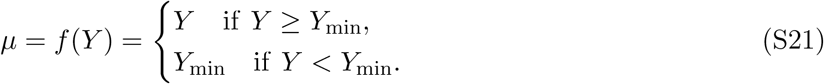

Note that at this point in time, our measurement of gene expression is noiseless, and we only introduced a detection limit. As the observed expression *μ* only depends on the true expression *Y*, we can include *μ* into the Markov chain,

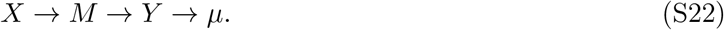

Before adding more steps to the pipeline, we note what is actually being computed: the mutual information between the identity of a specific base and the observed level of gene expression. This is informative because, a priori, the location of the binding site is not known. So far we have assumed we know where the binding site that controls gene expression is, and which mutations lead to changes in binding energy. But assume we don’t know that a priori, and try to find where it is. As we have noted above, a mutation outside a binding site leads to no change in gene expression, and therefore, mutual information is zero. Hence, if we compute the mutual information *I*(*X*_*i*_; *μ*) for every base *i* in our sequence, we would find an island of high mutual information where the binding site is, and a sea of zero mutual information outside, which would tell us exactly where the binding site is located.

If we wanted to compute mutual information at this point, we could use equation (S1), rewritten to use the current variables,

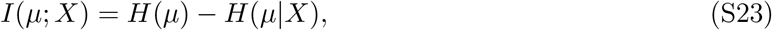

which we will proceed to compute analytically now. Readers interested only in the result may skip to equation (S60). To compute the marginal entropy *H*(*μ*), we need to know the marginal probability *p*(*μ*). We can always express a marginal probability as sum over conditional probabilities. As *μ* only depends on the underlying true expression *Y*, we can write

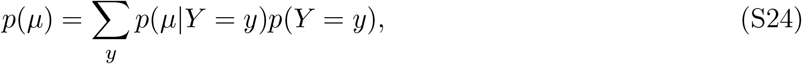

where we went back one step in the Markov chain. We can repeat this process, as *p*(*Y*) only depends on the number of mutations *M* in the sequence through equation (S18),

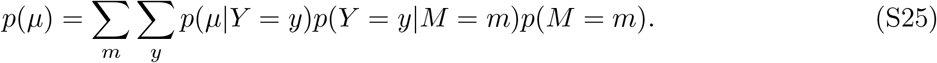

Repeating the process another time, going back to the first variable in our Markov chain, we write

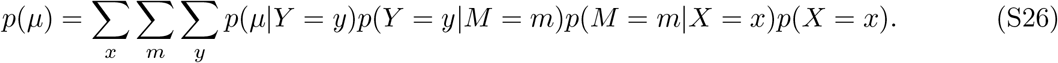

This decomposition allows us to write closed-form expressions for each conditional probability, and to evaluate the sum analytically. For example, the probability of observing *μ* can be expressed as

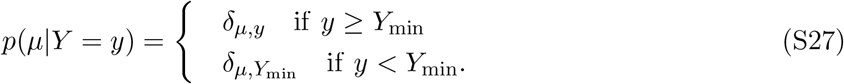

Since the true expression *Y* is a deterministic function of the number of mutations, as expressed in equation (S18), *Y* also takes on discrete values and the conditional probability *p*(*Y* = *y*|*M* = *m*) collapses to a delta,

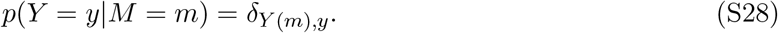

Finally, the probability *p*(*M* = *m*|*X* = *x*) of finding *M* mutations in the sequence, given the state of a specific base *X*, is defined by a binomial distribution, as we have defined above in our toy example in equation (S12). With all these conditional probabilities in hand, we can compute the marginal probability of observed expression,

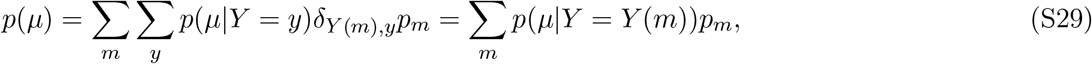

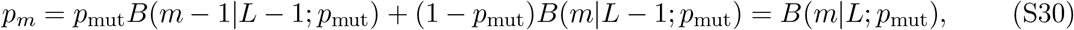

where we wrote *B*(*m*|*L*; *p*_mut_) for the probability of getting *m* mutations in a sequence of length *L* with mutation rate *p*_mut_ per position, and used the Kronecker deltas to collapse the sums. The terms in *p*_*m*_ describe the two possible ways to go from either *m* or *m* − 1 mutations in *L* − 1 positions to *m* mutations in *L* positions, and therefore, combine to the binomial distribution again.

Next, we define a maximal number of mutations for which we can measure expression above the threshold,

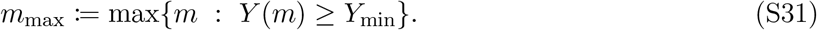

Here we implicitly assumed that Δ*h >* 0, and therefore, *Y* (*m*) is a monotonically decreasing function of *m*, the more mutations there are, the less expression there is. We use this to split the sum in equation (S29) into

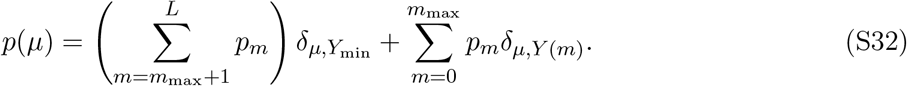

Here we have one component for the probability mass that accumulates at *μ* = *Y*_min_, as the detection limit does not allow us to resolve any expression below the limit. Using this expression, we compute the marginal entropy,

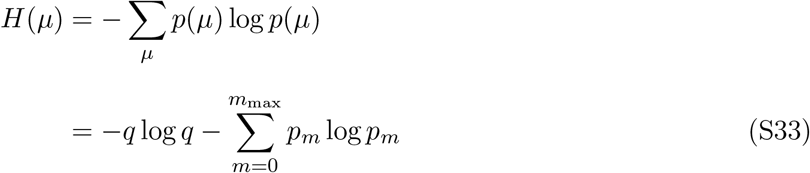

where

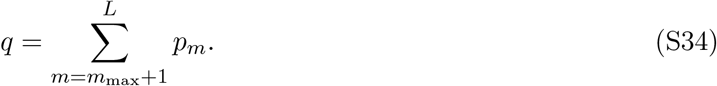

Here, we were able to separate the contributions to entropy between the probability mass below the detection limit from the probabilities of finding a measurement above due to the Kronecker deltas in the summation of the marginal entropy. This allows us to nicely show the change of entropy in equation (S33) due to detection limit by expanding the sum,

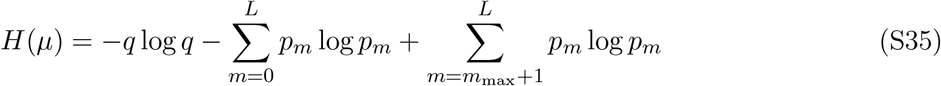

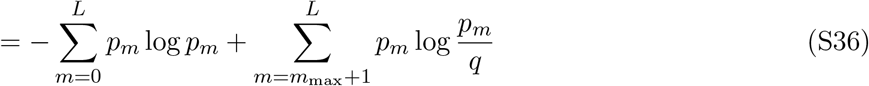

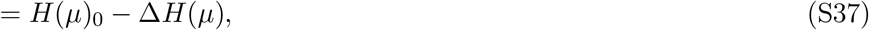

where we defined the entropy in the absence of a detection limit as

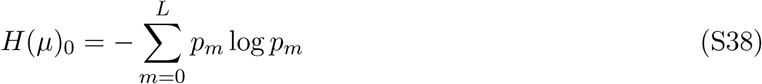

and the entropy cost of the detection limit

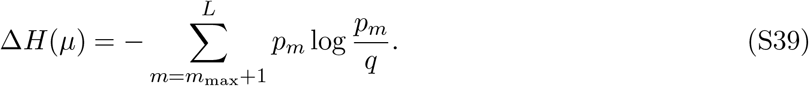

A quick check shows that if *m*_max_ = *L*, thus there is no detection limit, the sum term in the cost vanishes. Note that *p*_*m*_ was used as placeholder of the binomial distribution in equation (S30), and as such, the entropy in the absence of a detection limit is the entropy of the binomial distribution, as we found in our coin flip toy example. We now compute the conditional entropy. The conditional entropy is computed as

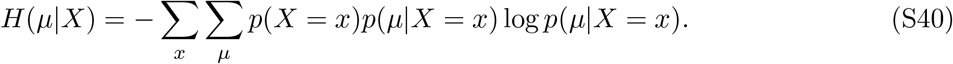

Here we need to compute the conditional probability first. We can write the conditional probability in a similar way to the marginal probability in equation (S26),

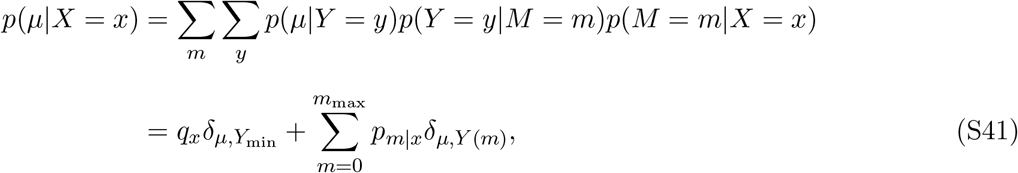

with

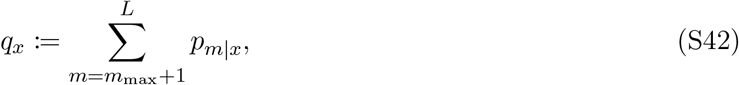

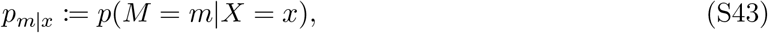

being introduced for easier notation. Here we are plugging in again the Binomial distribution,

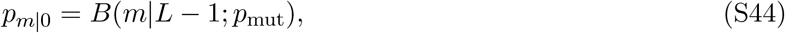

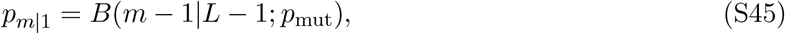

which gives us

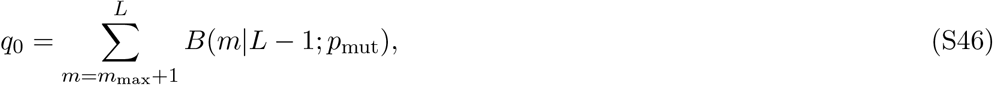

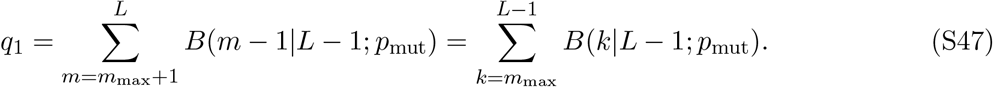

We can plug this directly into equation (S40) to get

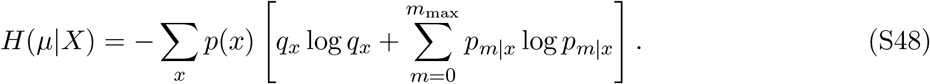

Following the same approach used for the marginal entropy, we expand the sum to write the conditional entropy in terms of entropy without thresholding and a cost term, we expand the sum to write the conditional entropy in terms of entropy without thresholding and a cost term,

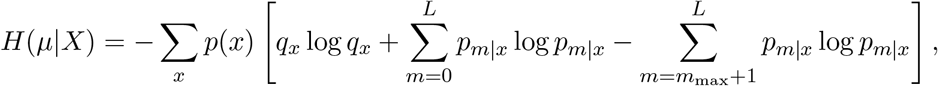

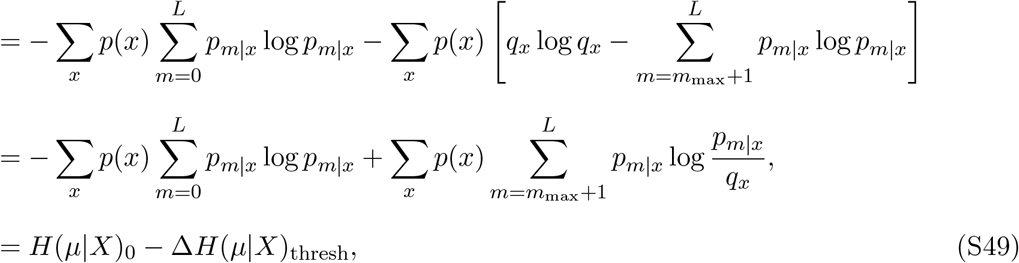

with

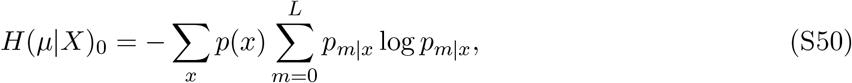

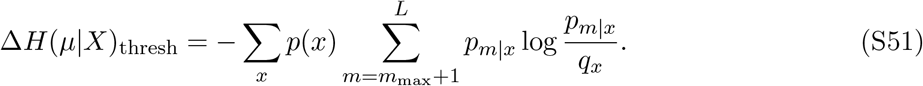

At this point, we are fully equipped to compute mutual information when we include the detection limit,

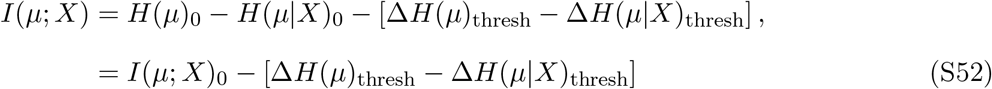

Here we defined the mutual information in the absence of a detection limit as *I*(*μ*; *X*)_0_. Now we plug in the derivations for Δ*H*(*μ*)_thresh_ and Δ*H*(*μ*|*X*)_thresh_ from equation (S39) and equation (S51) to get

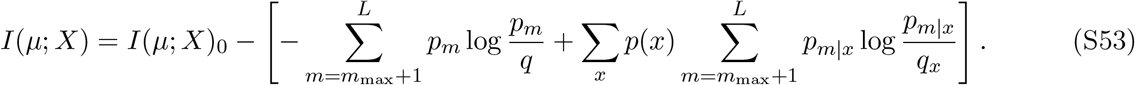

Now by reorganizing some terms, we can simplify this expression. First, let’s define truncated distributions,

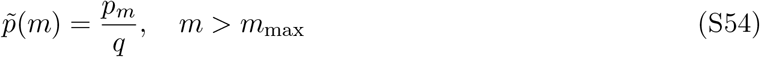

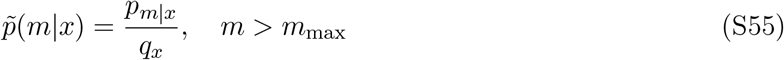

which are properly normalized in their regime, as denominator is a sum of the numerator, 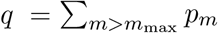 and 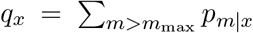. Next, we define a truncated version 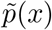, which we compute using Bayes theorem,

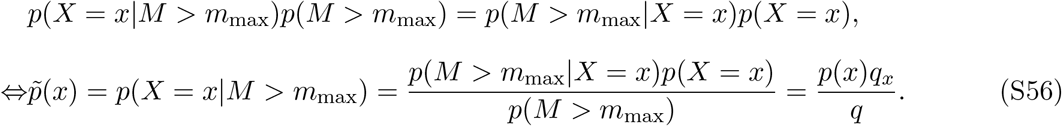

Plugging all these definitions into equation (S53), we get

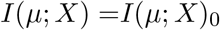

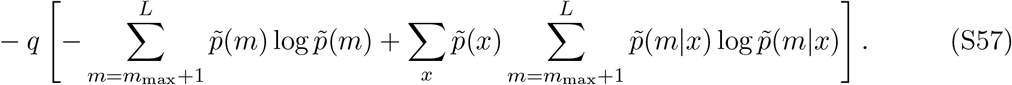

The expression in parenthesis is recognizable as a difference between marginal and conditional entropies,

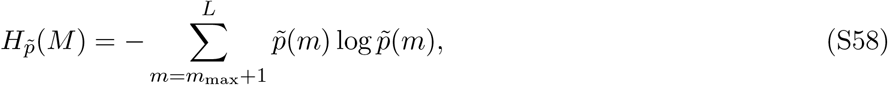

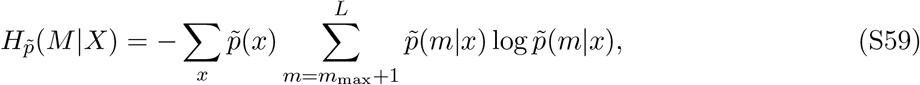

where we used the index 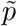 to indicate that the entropies are only calculated over the number of mutations over the detection limit. The parenthetical expression in equation (S57) is mutual information evaluated in the truncated regime,

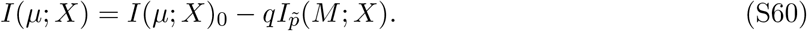

This result quantifies how a detection limit for the measure of gene expression reduces the mutual information we can get between the observed level of gene expression *μ* and the identity *X* of a specific base. Mutual information is reduced by the mutual information between the base identity *X* and the number of mutations *M* in the sequence for the outcomes where *M > m*_max_, and the difference is weighted by the total marginal probability mass that is taken by these outcomes *q*. We verify two limiting cases, what happens when *m*_max_ = 0 and *m*_max_ = *L*. In the case of no detection limit,*m*_max_ = *L*, we get that *q* = 0, since the sum has no terms, which is also true for the calculations of 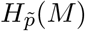 and 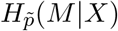 in equations (S58) and (S59) respectively. Hence, we simply get back the entropy we started with

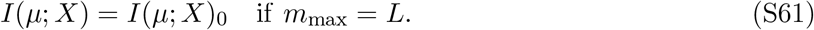

Let’s consider the case of *m*_max_ < 0, which simply means that every possible measurement of gene expression is under the detection limit. In this scenario, we cannot measure the number of mutations in the sequence at all. First we observe that *q* = 1, as it becomes a sum over a normalized probability. Then, we find that the definition of entropy in equation (S58) becomes equal to the entropy without a detection limit in equation (S38). Furthermore, the conditional entropy in equation (S59) becomes equal to the conditional entropy without a detection limit in equation (S50). Therefore, we get

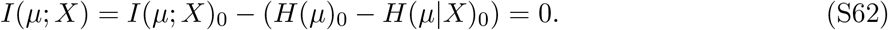

In the case where we cannot measure the number of mutations in the sequence, the two random variables essentially become independent, and mutual information vanishes.

Here we were able to show how mutual information is reduced if a step in the Markov chain introduces an uncertainty in the observed expression, and calculate the difference analytically.

So far we have assumed that each measurement is perfect, and that there is no noise on the biological nor on the measurement level. We have assumed that each mutation comes with the same binding site defect. Now we will relax these assumptions one by one, and explain how this affects the Markov chain and calculation of mutual information.

##### S3.1.4 Addition of noise

We begin by adding biological noise. Even though cells can have the same genomic sequences and grow in the same environment, there are fluctuations on the cellular level. There can be fluctuations in the number of molecules across cells, such as RNA polymerase and regulators, but also fluctuations within cells at different points in time. Here, we are modeling this noise as one process, where we say the realized level of gene expression *Y* ^′^ is drawn from a Gaussian distribution where the true expression *Y* (*m*) is the mean,

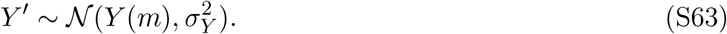

Since *Y* ^′^ is only a function of *Y*, we can include this step into our Markov chain,

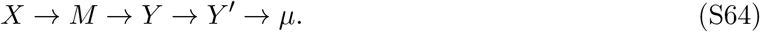

Now the measured expression *μ* only depends on the realized expression *Y* ^′^, so we redefine this step and write,

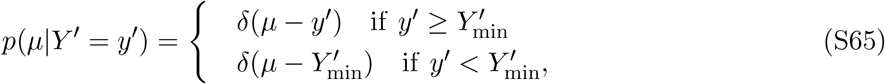

where the detection limit is now set on the realized expression *Y* ^′^, and we write Dirac delta functions instead of Kronecker Deltas, since the realized gene expression *Y* ^′^ is a continuous quantity, and as such, the measured gene expression *μ* is also a continuous quantity. To compute mutual information, we would have to compute again the marginal entropy *H*(*μ*) and conditional entropy *H*(*μ*|*X*). Since we are dealing with continuous quantities now, this turns sums into integrals, and makes it rather difficult to find closed expressions for the marginal and conditional entropy. The marginal probability of measured expression becomes,

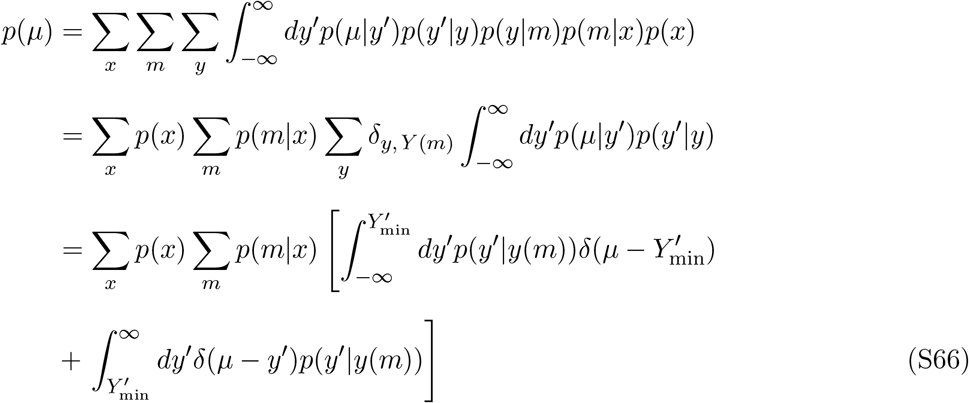

In our final expression we can see two integrals containing delta functions, but they affect the integrals differently. In the first integral, we are dealing with the case of expression below the detection limit. There, we have to perform a bounded integral of a Gaussian, *p*(*y*^′^|*y*(*m*)), which yields the cumulative distribution function of a Gaussian (an error function), and therefore does not admit a simple closedform expression for the entropy.

Before we compute mutual information, we introduce another step to our Markov chain, and that is binning. Instead of dealing with continuous measurements of gene expression *μ*, we discretize the outcomes by choosing a set of *r* bins *B* ={*b*_1_, *b*_2_, …, *b*_*r*_}. This has another practical motivation, as in experiments we get a limited number of samples, and binning allows us to approximate probability distributions of outcomes. We explore the practical aspects later in more detail, and will focus for now on the Markov chain. Since the bins only depend on the measured expression *μ* by definition, we can write the Markov chain as

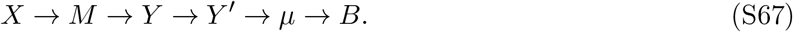

Mathematically, we define the conditional entropy of observing a bin *B*_*i*_,

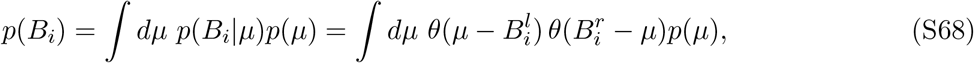

where *θ* denotes the Heaviside-theta function, and 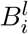 and 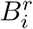 are the lower and upper bounds of bin *i* respectively. Here, we are using arbitrary bins for now, but a common choice in practice is to either choose equal bin widths or equally distributed bins, where each bin has the same probability of being observed, and bin edges are chosen accordingly. When we are computing mutual information, we are then computing

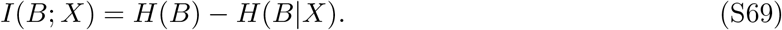

As shown in Figure S6, mutual information decrease with expression noise, but increases with the number of bins. This intuitively makes sense. If measuring the expression is noisy, we lose information. By increasing the number of bins, our approximation of the continuous distribution of measured expression becomes more accurate, hence, we increase mutual information. So why would one ever choose a smaller number of bins? This leads is to a crucial difference between our theoretical treatment of mutual information and the way it is calculated in practice from data.

**Figure S6.**
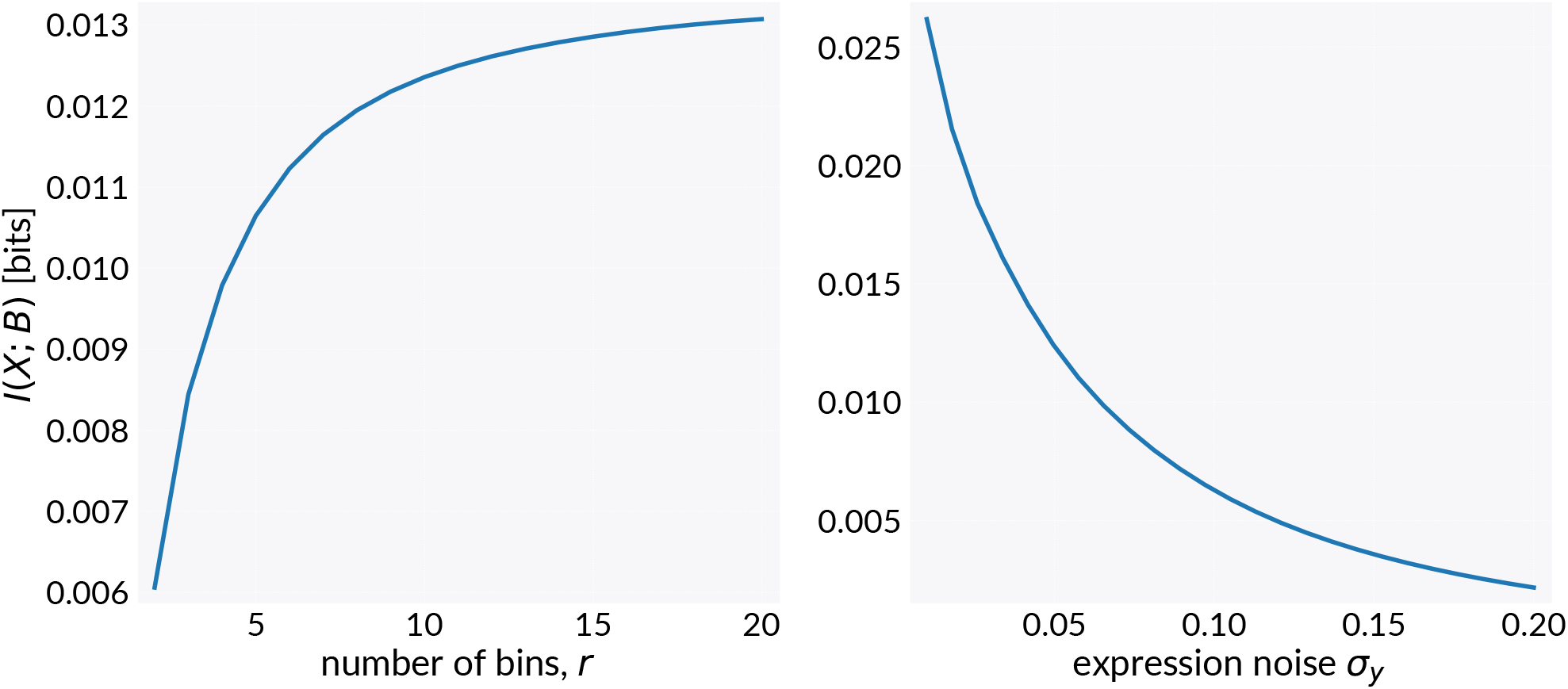
Effects of binning and expression noise on mutual information. Mutual information *I*(*X*; *B*) computed using the marginal probabilities from equation (S66) and equation (S68), and the corresponding conditional probabilities. Left: For fixed expression noise (*σ*_*Y*_ = 0.05) mutual information is shown as function of number of bins *r*. Bins are chosen such that *p*(*B*) is uniform. Right: For fixed number of uniform bins (*r* = 10) mutual information is shown as function of expression noise *σ*_*Y*_. Other parameters: *L* = 20, *p*_mut_ = 0.1, *Y*_min_ = 0.3, *h*_*w*_ = −6*k*_*B*_*T*, Δ*h* = 1*k*_*B*_*T*.

##### S3.1.5 Computing mutual information from data

Up to this point, we have been strictly talking about true mutual information using analytical expressions. This assumes we have a precise and accurate description of the probability distribution *p*(*X, B*). In practice, we are estimating this distribution with a subset of all possible sequences. Using a mutation rate of 0.1, the average number of mutations in the sequence of length 160 is 16. The total number of possible mutant sequences is then

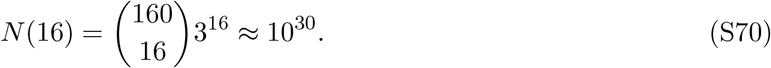

If we approximate the weight of one nucleotide to be 300 Daltons, then just having a one single stranded copy of each of these sequences would lead to a library with a total weight of

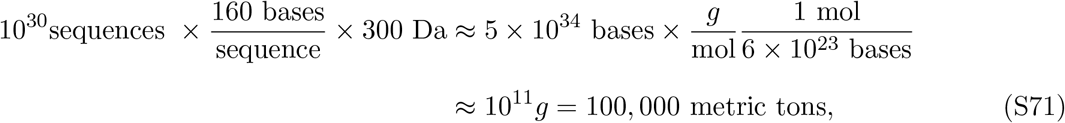

which is as much as a large U.S. Navy aircraft carrier. Obviously we cannot run experiments with such library of DNA. Thus, we sample from the total space of possible sequences and create a smaller library. Experiments then give us an estimate of the probability distribution *p*(*B, X*) which we will call 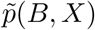. Since we estimate the underlying distribution, we also get an estimate for the mutual information, which we will call *Ĩ*(*B*; *X*). Finite sampling does not change the true mutual information; it changes our estimate of it.

Analytically, it was simple to write down mutual information in terms of entropy and conditional entropy, and taking this viewpoint was very insightful. However, in practice, it is much easier to compute mutual information in terms of joint distributions and marginal distributions, which we can directly compute from the data itself,

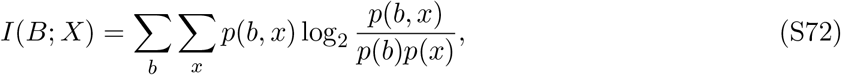

where we can plug in our estimated probabilities directly

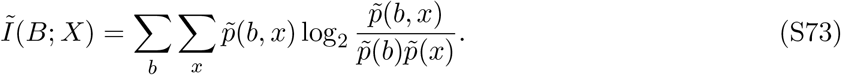

The estimator *Ĩ* itself is a random variable, as it depends on the outcome of the experiment. There exist multiple expressions for the mean of the estimator, and one established one is the Miller-Madow formula [112],

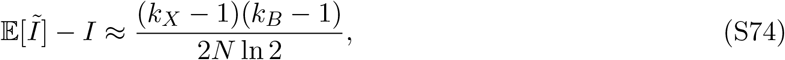

where *k*_*X*_ is the number of outcomes of *X*, hence *k*_*X*_ = 2, *k*_*B*_ is the number of bins, and *N* is the number of sequences. In experiments based on sequencing data, expression is measured as barcode of counts in RNA sequencing relative to DNA sequencing. An alternative way to binning expression as ratio of counts, is to associate bins with RNA and DNA reads separately. We introduce a random variable *Z* ∈ {DNA, RNA}, which says if a sequencing read is coming from DNA sequencing or RNA sequencing. For sequences with high expression, finding a RNA read is more likely than for a sequence with low expression. This binning strategy removes the need of choosing a binning strategy for continuous expression measurements and reduces the noise of barcodes with low counts by putting more weight on barcodes with high counts. However, this can be problematic if there are a few barcodes that take up a majority of the counts. Let *D*_*b*_ be the measured DNA counts for barcode *b*, and *R*_*b*_ the amount of RNA counts. The total number of counts is then given by *T*_*b*_ = *D*_*b*_ + *R*_*b*_. We can compute barcode level conditional probabilities, meaning how likely it is for a count to be either RNA or DNA,

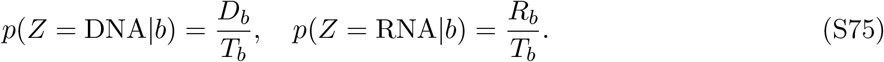

If we now look at a certain position in the sequence, we can compute the joint probability of having a mutation and if the read is RNA or DNA,

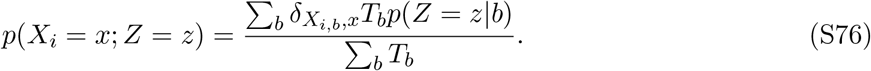

Here we simply sum up DNA or RNA counts for all barcodes that either have a mutation or the wild type base, and normalize by the total number of counts. If a few barcodes with high count numbers dominate the distribution, one can choose different ways to compute the weights *T*_*b*_, such as the square root of the sum of counts 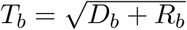.

From equation (S74) we can see by using sampling from the library of possible sequences, we overestimate mutual information, and by using more bins we increase the amount we overestimate by. We can improve the estimate by increasing the size of the library. Importantly, this positive bias exists even when the true mutual information is zero, and we see this in practice, where there is mutual information measured across the entire sequence. In the experiments, there are promoter sequences for which we did not detect any signal, meaning no binding sites. How this was concluded is explained below, but for now we can use these promoters to investigate the scaling of mutual information with the number of sequences identified for each promoter. Figure S7 shows how the maximum mutual information scales with the number of sequenced barcodes for a few noisy promoters. We can see a clear scaling with 1/*N*, showing how our estimate of mutual information scales as described by equation (S74). This also gives us an estimate for the noise for footprints that contain noise. We can check how many barcodes we find for a promoter in a given condition, and where it falls on the curve in Figure S7, which we can use as baseline for any signal we are trying to identify. Figure S8 shows in the results for the *araB* promoter, which was discussed in detail in the introduction. Here, cells were grown in the presence of arabinose, and expression is induced from the promoter. This can be seem from the peak at the −10 region, which is indicative of binding by *σ*_70_. Then, there are multiple peaks indicating binding of AraC. The peaks at −35 and −50 belong to one binding site, while the peaks at −60 and −75 belong to the other. Once the background is subtracted, the binding sites are easy to identify. We apply this threshold to any measurement of gene expression for a promoter to identify binding sites. Now we have identified regions where mutations significantly change expression. However, the procedure is agnostic to the sign of change, meaning if a mutation increases or decreases expression. This is very important information, since we want to know if a repressor or activator binds to our binding site of interest.

**Figure S7.**
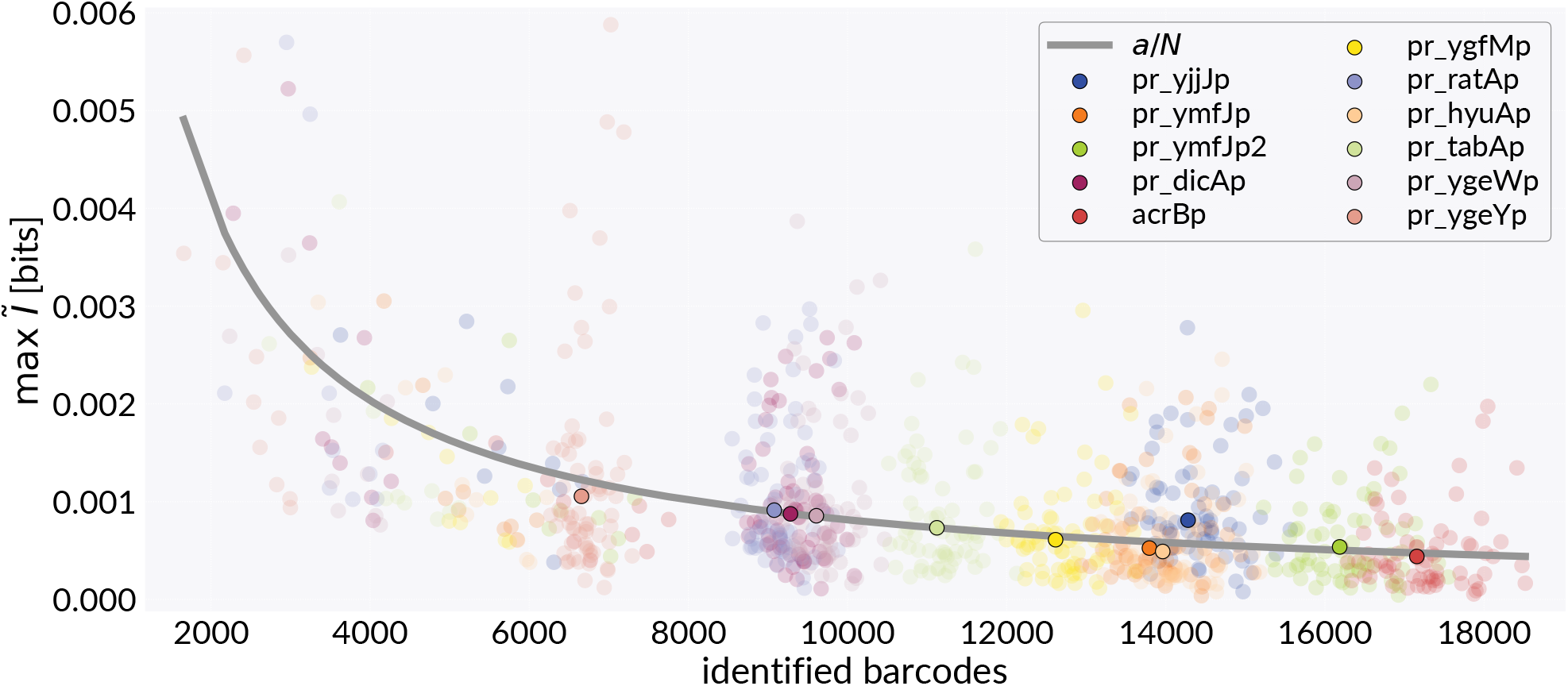
Scaling of maximum mutual information with number of barcodes. Maximum mutual information of noisy promoters across all growth conditions. The suffix “pr_” indicates the promoter location was predicted using computational models, which are explained below.

#### S3.2 Expression shift matrices

Now we are going back to looking at promoters in individual conditions and ask the question how much expression is changed for every possible point mutation in the sequence. The important feature is that this quantity is signed, meaning for mutations that increase expression, expression shift is positive and vice versa. Also, we can compute expression shift for every individual base at every position, and we can resolve imperfections in binding sites. For example, on average mutations in activator sites are going to reduce expression, since binding of the activator is weaker. However, in reality, most binding sites for transcription factors are imperfect, which means that there are specific mutations that can actually increase binding of the transcription factor. While mutual information is agnostic to this, expression matrices can resolve such features. We compute the expression shift Δ*s*_*x,l*_ for base *x* at position *l* as

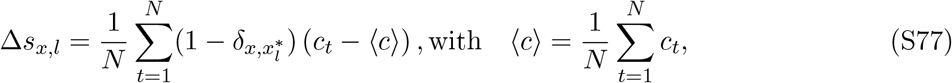

where *N* is the total number of barcodes, 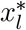 is the wild type base at position *l*, and *c*_*t*_ is the measured expression for barcode *t*. We compute expression shift as difference to the average expression of the promoter across all its variants. If we want to compare expression shifts across different conditions, we need to take into account that expression measured as number of RNA counts to DNA counts is sensitive to the total number of counts, or in other words, read depth. If I accumulate twice as many total RNA reads, relative expression for all barcodes will also go up by a factor of two. Hence, we have to normalize with respect to read depth, and we do that by finding the median of relative expression across all barcodes from all promoters in the library within each experiment. Let *λ* be the median, then we compute the scaled expression shift as

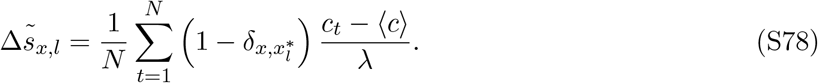

**Figure S8.**
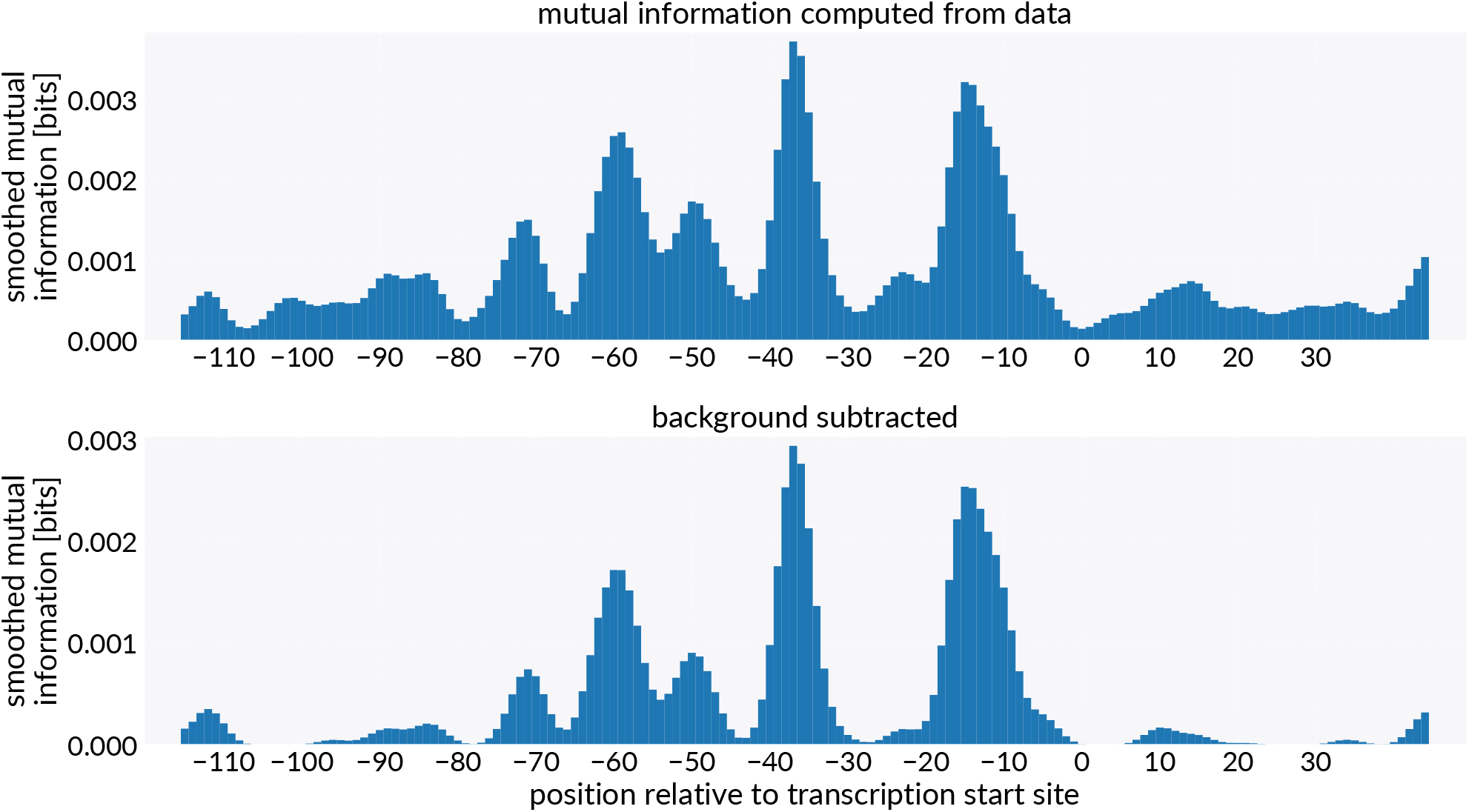
Background subtraction for information footprints. Top: Information footprint for the *araB* promoter for cells grown in the presence of arabinose. Bottom: Information footprint after background subtraction, where the level of background was determined from the fit in Figure S7. Both footprints were smoothed using a Gaussian Kernel of width *σ* = 2.

As shown in Figure S9, we can combine the expression shift with mutual information footprints for a more complete picture of binding sites. In the example the footprint for the *araB* promoter is shown, where cells were grown in the presence of arabinose. We can see the −10 region of the sigma factor, as well as the binding sites for AraC. By coloring bases in the mutual information footprint by the average expression shift across all bases, we can see that there are two bases in the −10 region where mutations increase expression by bringing the sequence closer to the optimal −10 sequence. The activator binding sites mostly contain bases where mutations decrease expression as expected, but there are also a few mutations where expression is strongly increased. Throughout this work, footprints are colored by expression shift.

**Figure S9.**
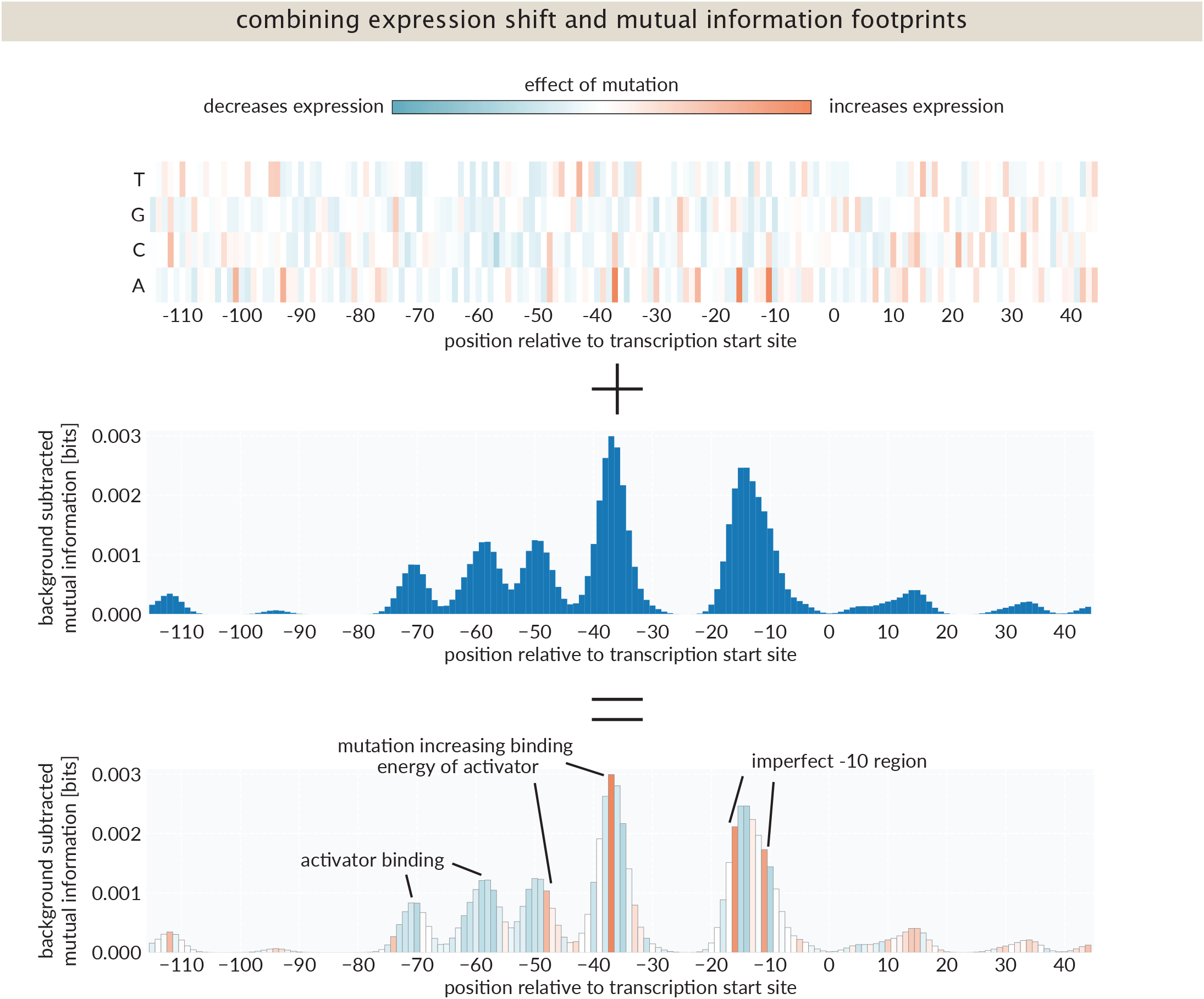
Combination of expression shift and mutual information. Expression shift matrix and mutual information footprint for the *araB* promoter for cells grown with arabinose.

### S4 Per-promoter accounting of annotated binding sites

A per-promoter accounting of every RegulonDB binding-site annotation within our 160 bp mutagenesis windows is provided as **Supplementary Note 1**. The accounting covers all 43 promoters in our pool with previously annotated regulation and addresses each of the 143 in-window binding-site entries individually. For each entry, we record the transcription factor, position, and RegulonDB confidence level, and classify the outcome of the Reg-Seq experiment as one of: clear footprint, weak footprint, no footprint, or untestable in our assay. For each non-recovered annotation, we provide a rationale drawn from the categories described in the main text: degenerate binding by nucleoid-associated factors (Fis, HU, CspA), cooperative architectures whose partners fall outside the reporter window (e.g. AraC looping), low promoter activity in our reporter, conditions outside our experimental panel, and W-confidence annotations supported only by sequence similarity to a TF consensus. Where relevant, we also note in vivo evidence from the primary literature (genetic, biochemical, or chip-based) that bears on whether a non-recovered annotation reflects a methodological limitation of Reg-Seq or a genuine absence of regulatory function under the conditions tested.

### S5 Supplementary Results

#### S5.1 Position weight matrices from expression shifts

In two parts of this work, we construct position weight matrices (PWMs) from a small number of binding-site sequences. PWMs are usually calculated from sequence alignments and quantify how preferred each base is across positions in a binding site. Any DNA sequence can then be evaluated against a PWM to score how well it matches. Computing a PWM from a small number of sequences is inherently noisy. However, our data provides expression shift matrices (Section S3.2), which contain information about how expression changes for each possible mutation in the sequence. We use these to weight the PWM by the effect of mutations on expression, giving a more direct map from sequence to expression. Below we describe the construction in two settings: (i) the activator binding sites at the *yjbJ* and *ybaY* promoters under osmotic stress, and (ii) the *σ*^*S*^ − 10 element across stationary-phase promoters.

##### S5.1.1 Activator binding sites under osmotic stress

For each promoter *p* and each position *l* within the identified binding site, we have the scaled expression shift 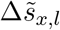 for each base *x* ∈ {*A, C, G, T*}, as defined in Section S3.2. The wild-type base has 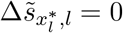 by definition. We average the expression shifts across the two strong sites (the activator binding sites in the *yjbJ* and *ybaY* promoters) at each aligned position, weighted by the total signal strength at each promoter,

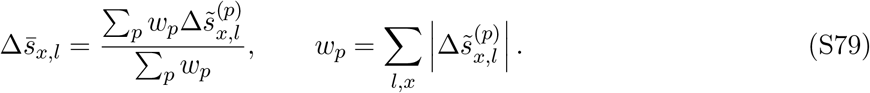

Since the expression shift is not isolated to the activator binding site, but also depends on other parameters such as binding of sigma factors and interaction energies of activators to RNA polymerase, the weights normalize the expression shift for each promoter. Choosing the sum of absolute expression shifts as weight reduces the impact of the stronger expressing promoter on the resulting PWM.

We convert the averaged expression shift matrix to base frequencies using a Boltzmann transformation,

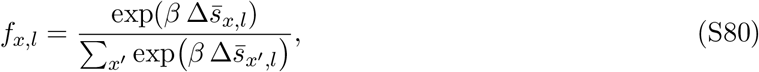

where *β* is an inverse temperature parameter matching the scale of the expression shifts. Mutations that disrupt binding produce large negative 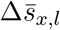 and therefore receive low frequency, while the wild-type base 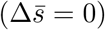 receives high frequency. The log-odds PWM is

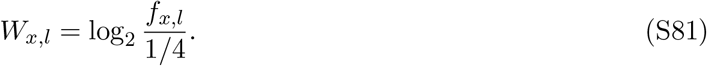

The information content at each position is IC_*l*_ = 2 + ∑_*x*_ *f*_*x,l*_ log_2_ *f*_*x,l*_. The absolute information content per site depends on the choice of weights in equation (S79); high information content indicates that the specific base at that position is important for expression and, by association, binding.

##### S5.1.2 Comparison of the de novo motif against known transcription factor PWMs

We compared the de novo motif against PWMs for all 145 transcription factors with at least 3 annotated binding sites in RegulonDB, using two complementary similarity measures.

The first is the Pearson correlation *r*_TF_ between the de novo PWM 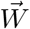 and each known transcription factor PWM, maximized over all alignment offsets and both orientations:

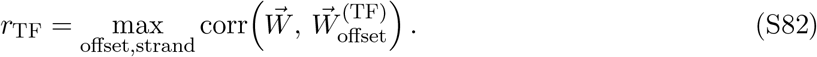

Since we do not have expression shift data for other transcription factors (especially not in the same condition), the PWMs for known TFs are computed from sequence alignments alone. These motifs are therefore far less informative about expression, and we use a second measure to complement the comparison.

For the second measure, we evaluate every known binding site for a transcription factor against the de novo motif. For each TF, we score its *N* known binding site sequences *a*^(*j*)^ with the de novo PWM and normalize by the self-score, the mean score of those same sites under their own PWM:

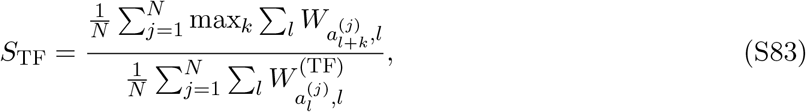

where the maximization over *k* accounts for the unknown alignment of each site relative to the de novo motif if the lengths differ. A value of *S*_TF_ = 1 would indicate that the known binding sites for the TF score as well against the de novo PWM as they do against their own.

To establish a baseline for non-specific similarity, we scored 1000 random genomic sequences with the de novo PWM and computed the same normalized score for each TF. The 99th percentile of this null distribution represents the upper bound expected in the absence of specific binding.

##### S5.1.3 Genome-wide motif scan

To identify additional candidate targets of the unknown activator, we scanned every *E. coli* K-12 MG1655 promoter region with the de novo PWM. For each of 4726 annotated genes, we extracted the 400 bp immediately upstream of the coding sequence start, oriented in the 5^′^ → 3^′^ direction relative to the gene. We slid the 20 bp PWM across all positions on both strands, retaining the maximum score:

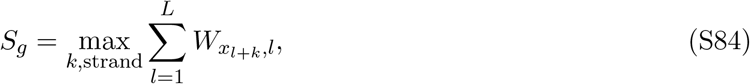

where *x*_*l*+*k*_ is the base at position *l*+*k* in the promoter (or its reverse complement). To compare scores across genes, we computed a robust *z*-score,

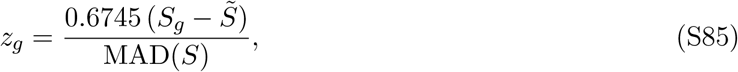

where 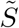 is the median score across all genes and 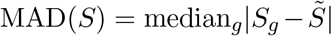. The factor 0.6745 makes the robust *z*-score comparable to a standard *z*-score for normally distributed data.

##### S5.1.4 Motif from stationary-phase promoters

We applied the same approach to construct an expression-weighted motif for the *σ*^*S*^-dependent −10 element, using 13 stationary-phase promoters with single-peak footprint architecture (Figure 6). The promoters and the wild-type sequences from −13 to −7 relative to the aligned −10 peak are listed in Table S3. Information footprints for each promoter were aligned at the dominant peak position, identified by Gaussian smoothing (*σ* = 2 bp) followed by local maximum detection, with all promoters aligned relative to the −10 peak of the template promoter (the one with the highest total information signal).

**Table S3.** Stationary-phase promoters used to construct the *σ*^*S*^ −10 PWM. Wild-type sequences are shown from position −13 to −7 relative to the aligned −10 peak, spanning the C(−13) discriminator and the canonical −10 hexamer.

| Promoter | −13 to −7 |
| --- | --- |
| *ybaY* | CCACACT |
| *ecnBp* | CTATTAT |
| *yqjE* | CTATGAT |
| *fldAp* | CTATGAT |
| *ftsKp1* | TTACACT |
| *kbpp4* | CTACACT |
| *yjbJ* | CTACACT |
| *icdC* | CTATAGT |
| *elaBp* | CTATAGT |
| *yadE* | CTACAAT |
| *ompRp3* | GTATACT |
| *sohAp* | TTATAAT |
| *furpa* | CTATAAT |

Each promoter’s expression shift matrix *E*^(*p*)^ is normalized by its maximum absolute value, 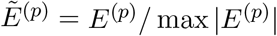, so that all promoters contribute equally in shape regardless of absolute expression level. Each promoter receives a weight 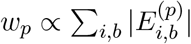, reflecting confidence that genuine regulatory contacts are present. Weights are capped at 15% to prevent any single promoter from dominating and renormalized to sum to unity. The promoter-level matrices are combined,

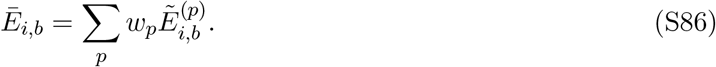

The averaged matrix is converted to base frequencies using equation (S80) with *β* = 10.

Because we now have 13 sequences rather than 2, we can additionally use sequence conservation across the aligned binding sites. We supplement the expression-derived frequencies with the observed wild-type base frequencies through a sequence blend,

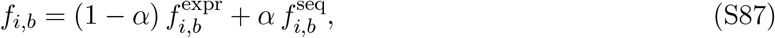

where 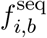 is the frequency of base *b* at position *i* across the aligned wild-type sequences (with a pseudocount of 0.5), and *α* = 0.5. This blend rescues positions where the wild-type base is essential but the expression assay cannot distinguish among the uniformly deleterious mutations: any of the three substitutions abolishes binding, so all three appear equivalently bad in expression shift, but the wild-type base is highly conserved across the aligned set. The final PWM is calculated from equation (S81) and information content from IC_*i*_ = 2 + ∑_*b*_ *f*_*i,b*_ log_2_ *f*_*i,b*_.

To avoid circularity when scoring promoters against the motif we report leave-one-out (LOO) scores: for each promoter *p*, the PWM is rebuilt from all other promoters and *p* is scored against this held-out motif. A positive LOO score indicates that the promoter’s sequence is consistent with the consensus derived from independent data.

#### S5.2 CRP PWM construction and ChIP-exo analysis at *yadI*

To assess whether the activator peak in the *yadI* promoter is consistent with a CRP binding site, we constructed a position weight matrix from 269 CRP binding sites with strong or confirmed evidence in RegulonDB. The PWM uses standard log-odds scoring with a uniform background and pseudocounts as in [113]. The best-scoring 20-mer within the activator peak region (GTTTTGACGGCTATCACCCT, positions −45 to −26 relative to the predicted transcription start site, dyad axis at −34.5) received a log-odds score of 8.69, placing it at the 36th percentile of known CRP sites. The full distribution of CRP site scores has median 10.38 and range −17.6 to 19.1, so the *yadI* site falls comfortably within the distribution of confirmed CRP targets, though on the lower-affinity side.

As independent corroboration, we examined a CRP ChIP-exo dataset [61], in which CRP binding was measured at single-nucleotide resolution in *E. coli* grown on glycerol minimal medium. The *yadI* locus is included among the CRP ChIP-exo peak regions in that study. The signal shows the TSS-centered protection pattern described as characteristic of CRP-activated promoters [61], as opposed to the motif-centered profiles seen at repressor sites such as ArcA and Fur. This is consistent with active CRP-dependent transcription at the *yadI* locus.

Mass spectrometry did not identify CRP at *yadIp* or at control promoters with confirmed CRP binding sites, indicating that CRP is not reliably recovered in our DNA chromatography setup. We hypothesize this reflects either (i) the relatively low affinity of CRP for any individual binding site combined with the large number of CRP binding sites genome-wide, which dilutes the pool of CRP available to bind the bait sequence, or (ii) loss of cAMP during lysate preparation despite supplementation. Because CRP is not detected at any of the eight other promoters in our pool with strong or confirmed CRP annotations, the absence of CRP at *yadIp* in mass spectrometry does not argue against the assignment.

#### S5.3 Novel transcription start sites

Of the studied promoters in this work, 29 had predicted transcription start sites. Here, we report putative start sites for 15 of these promoters. For the remaining 14 we either did not find any active sites, or the evidence was inconclusive to suggest a start site. For each of the 15 promoters, we identified the center of the −10 element as average across all conditions it was identified in. These locations are reported in Table S4.

#### S5.4 *De novo* promoters

A common feature in some promoters in our experiments was that single mutations could lead to the creation of a new putative transcription start site, different from the annotated one.

##### S5.4.1 Neighboring Transcription Start Sites

Earlier work on the 5’ ends of mRNAs found multiple species, where initially two start sites were identified [114], *tolCp1* and *tolCp2*, and later, two additional sites were found [115], *tolCp3* and *tolCp4*. One *tolC* site, (*tolCp1* and *tolCp2*) is active in most conditions and is known to be activated by PhoP in magnesium-limiting conditions. Indeed, in one of our replicates for magnesium starvation, we find an activator binding site at the annotated position at −46 relative to the annotated transcription start site for *tolCp2*. The footprints for *tolCp1* and *tolCp2* are very similar, with an offset that is equal to the annotated distance between the two start sites (Figure 7(B)), indicating that there is actually a single transcription start site with mRNAs of different lengths originating from this single start site. The other two annotated promoters, *tolCp3* and *tolCp4*, are 50 bases downstream, and have been found to be activated by MarA, SoxS, and Rob through a mar-box [115, 116]. Here, we only observe activation by Rob under its activating conditions, namely induction with 2,2-dipyridyl. The other annotated activators, i.e. MarA and SoxS, are not found to activate *tolC* from this site even under inducing conditions. In previous work, activation of these two transcription factors relied on overexpressing the proteins [115, 116], which may have led to conditions where protein copy numbers heavily exceed those in physiological conditions. We do observe activation by these transcription factors in these conditions on other promoters, such as the *acrZ* promoter. Interestingly, we find the binding site for Rob on the studied region for *tolCp2*, which contains the activator binding site and the −10 region of the *tolCp3* and *tolCp4* promoter on its 3’ end due to the proximity of the promoters, but not in the promoter regions for *tolCp3* and *tolCp4*. The sequence for *tolCp2* does not include the annotated binding site for the small RNA SdsR, which binds the 5’ UTR of the mRNA and reduces mRNA abundance [117], which explains why we only find noisy footprints for *tolCp3* and *tolCp4*. To summarize, for *tolC* we find two distinct transcription start sites rather than the four annotated sites. We find more examples for promoters with multiple annotated transcription start sites in our dataset, where only a subset of the annotated start sites are active, such as the promoter for *galE* and *crp* (Figure S12).

**Table S4.** Confirmed and discovered transcription start sites. Genomic positions of transcription start sites confirmed by Reg-Seq information footprints, based on the position of the *σ* factor contact points in conditions where both biological replicates agree. The four promoters in the top section were reported in Chapters 4 and 5. The remaining 14 promoters had only computationally predicted transcription start sites or database entries without experimental validation prior to this work. Genomic coordinates refer to *E. coli* K-12 MG1655. ^*†*^The *yahC* promoter shows a second, distinct *σ* factor pattern in stationary phase approximately 100 bp downstream of the primary pattern, suggesting a possible dual promoter architecture.

| Promoter | Strand | TSS (genomic) | Active conditions |
| --- | --- | --- | --- |
| *Previously reported, weak evidence* |  |  |  |
| *recNp* | + | 2,751,760 | H<sub>2</sub>O<sub>2</sub>, PMS |
| *cpxRp2* | − | 4,105,685 | Gentamicin, copper sulfate |
| *marRp* | + | 1,619,094 | Salicylate induction |
| *Newly confirmed transcription start sites* |  |  |  |
| *yacH* | − | 131,388 | Stationary phase, salicylate, starvations |
| *yadE* | + | 145,056 | Stationary phase, Mg starvation |
| *yadI* | + | 144,550 | Stationary phase, galactose, acetate |
| *yagB/insX/yagA* | − | 282,204 | Stationary phase, Mg starvation |
| *yahC*<sup>†</sup> | − | 335,425 | Broadly active |
| *yahL* | + | 344,159 | Broadly active |
| *yahM* | + | 344,477 | Broadly active |
| *ybaY* | + | 475,324 | Stationary phase, osmotic, starvations |
| *ybiY/ybiW* | − | 863,534 | anaerobic |
| *icdC* | + | 1,211,149 | Stationary phase |
| *tmaR* | − | 2,079,462 | Broadly active |
| *ygiW* | − | 3,169,717 | Stationary phase |
| *yqjE/yqjK/yqjD* | + | 3,248,893 | Stationary phase, osmotic, starvations |
| *zapB* | + | 4,118,431 | Constitutive |
| *yjbJ* | + | 4,259,180 | Osmotic stress, stationary phase |

**Figure S10.**
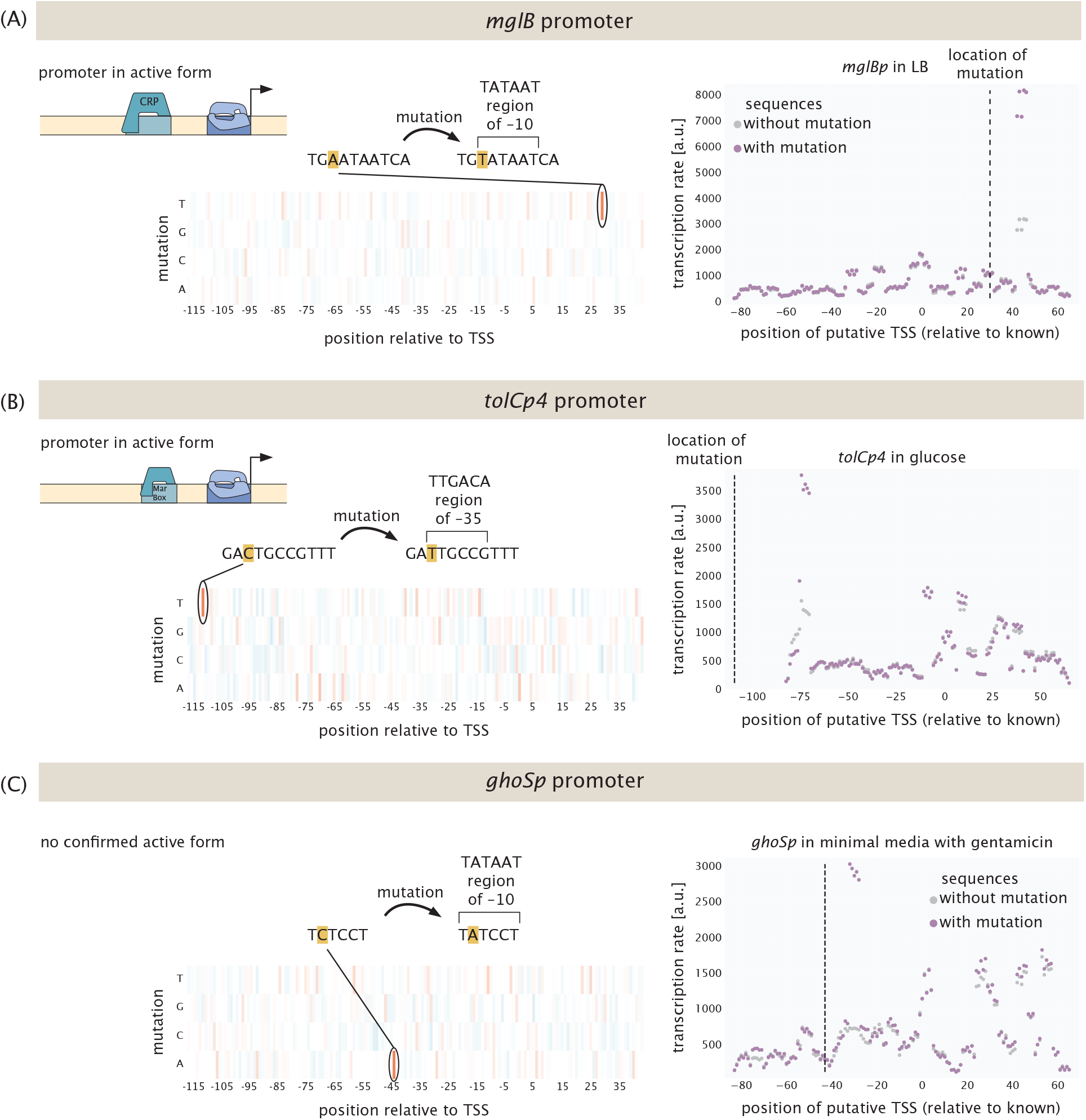
De novo emergence of new transcription start sites. Single mutations create new transcription start sites in the (A) *mglBp*, (B) *tolCp4* and (C) *ghoSp* promoters. Mutations either effect the −10 or −35 regions. Each of these promoters is activated under certain conditions. On the right, predicted transcription rates are shown for sequences with the suggested mutation and without the suggested mutation for each possible transcription start site.

**Figure S11.**
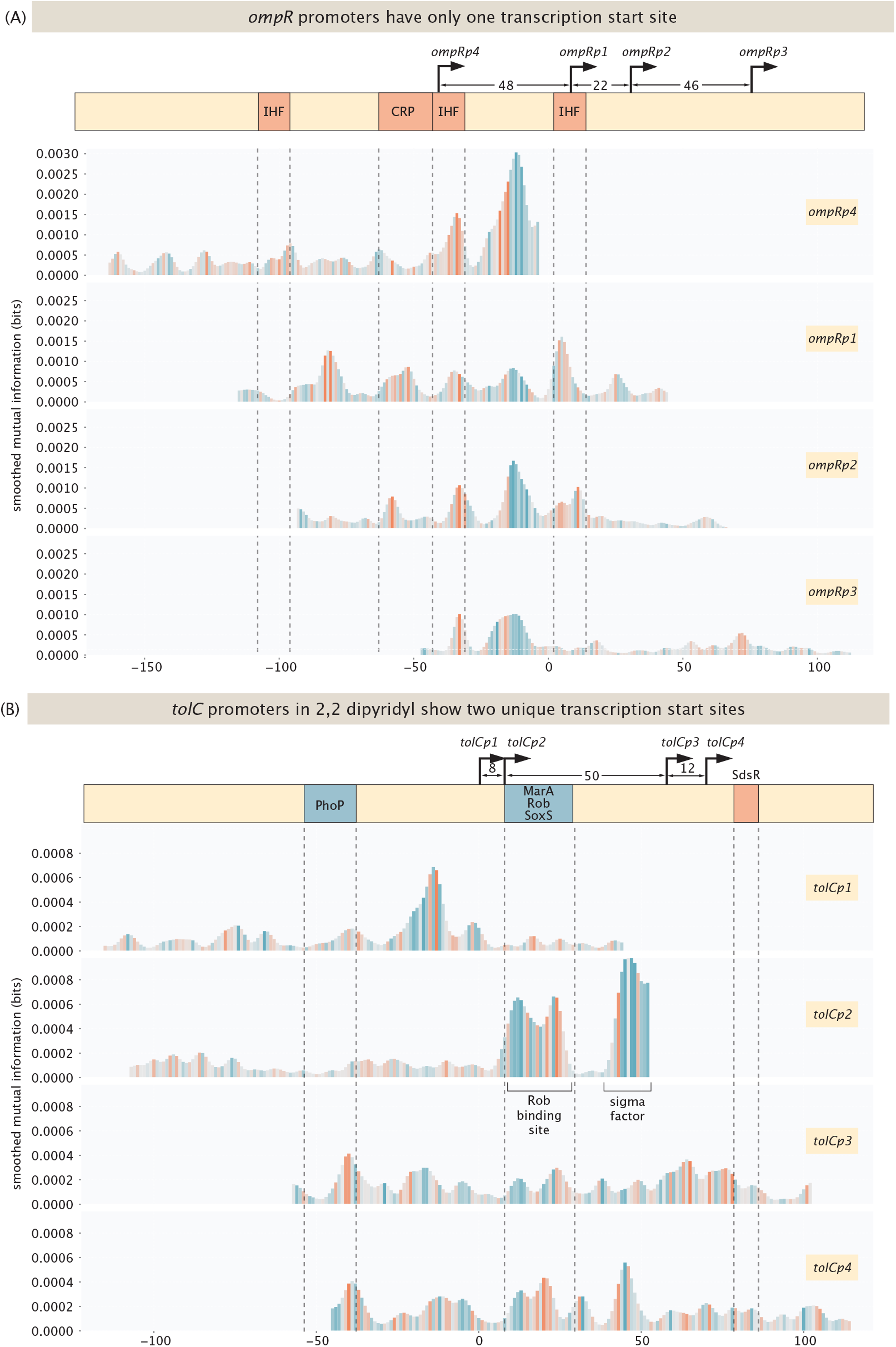
Transcription start sites for the *ompR* and *tolC* promoters. (A) Footprints for the four reported *ompR* promoters are shown for medium cold shock (19C for 1h), where the genomic locations of the footprints are aligned. Annotated binding sites are shown for IHF and CRP. (B) Footprints for the four reported *tolC* promoters are shown for induction with 2,2-dipyridyl, where the genomic locations of the footprints are aligned. Annotated binding sites are shown for PhoP, MarA, Rob, SoxS and the small RNA SdsR.

**Figure S12.**
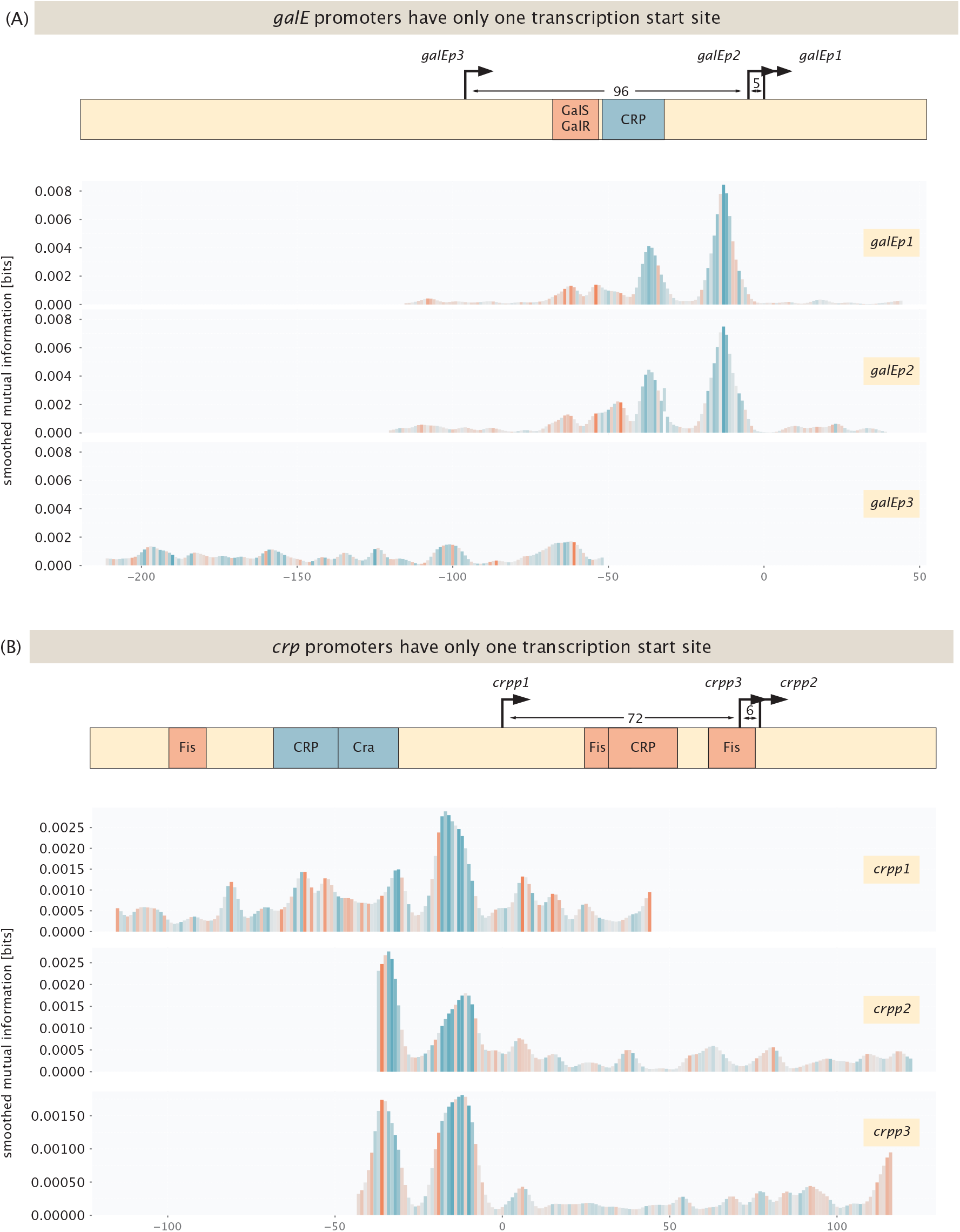
Transcription start sites for the *crp* and *galE* promoters. **(A)** The *galE* gene has three annotated transcription start sites (galEp1, galEp2, galEp3) within a window of ~ 100 bases, with annotated binding sites for GalS/GalR and CRP. Information footprints for the three reported promoters in 1-day stationary phase, with genomic locations aligned, show a single coherent footprint architecture (CRP activator site and the −10 element) consistent with one active transcription start site associated with galEp1/galEp2 rather than three independent initiation events. **(B)** The *crp* gene has three annotated transcription start sites (crpp1, crpp2, crpp3) with annotated binding sites for Fis, CRP, and Cra. Footprints for the three reported promoters, with genomic locations aligned, show a single active site associated with crpp1; crpp2 and crpp3 show footprints that are offset replicas of the crpp1 signal rather than independent initiation events.

## Supplementary Figures

**Figure S13.**
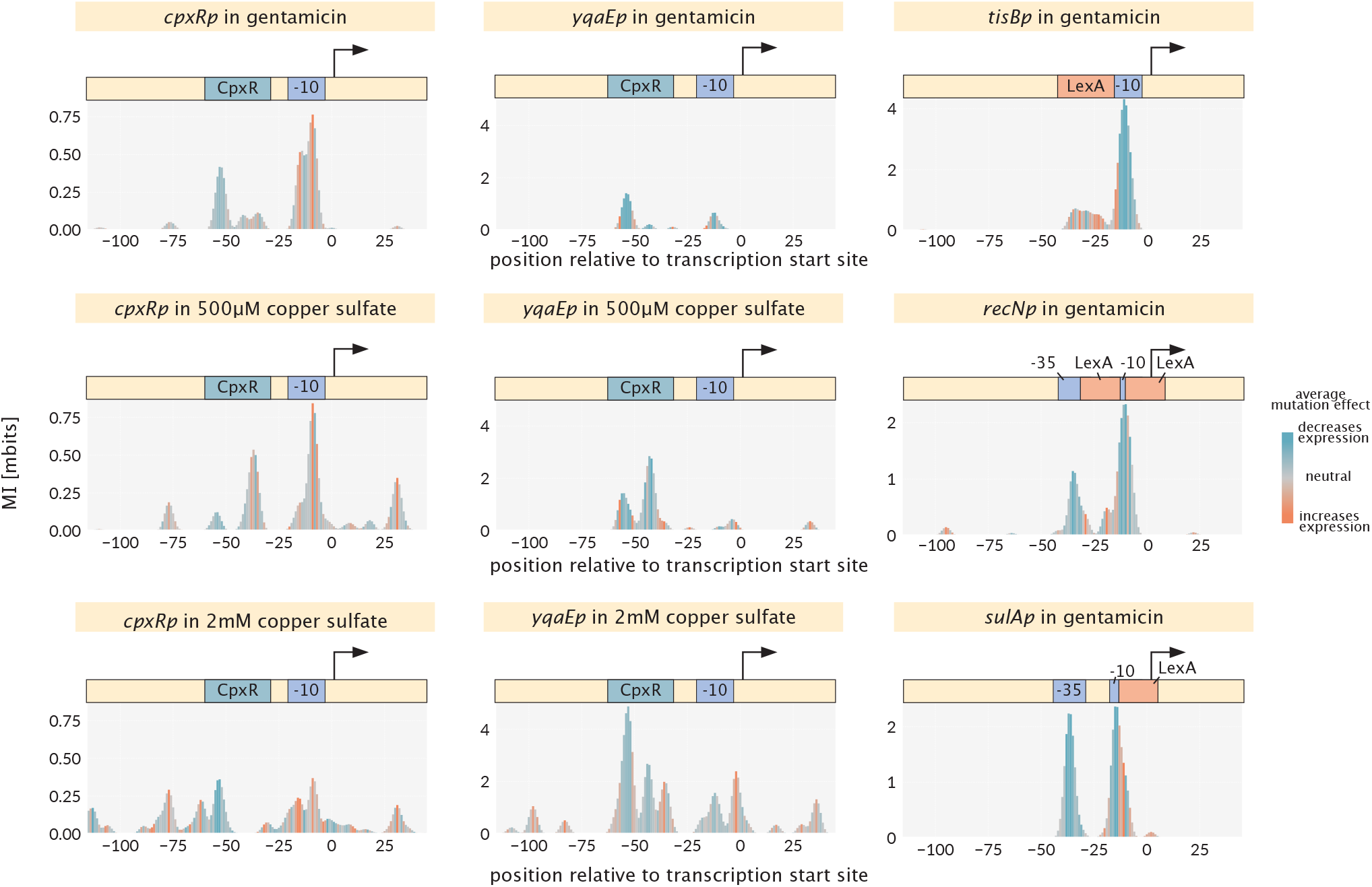
Information footprints supporting condition-dependent regulation by CpxR and LexA. Footprints for *cpxRp* and *yqaEp* under gentamicin, 500 μM copper sulfate, and 2 mM copper sulfate (left and middle columns), confirming the CpxR activator site shared by both promoters. Footprints for *tisBp, recNp*, and *sulAp* under gentamicin (right column), showing the LexA repressor signal at promoters in our pool. Annotated binding sites (CpxR, LexA) and core promoter elements (−10, −35) are indicated above each panel. Color encodes the average effect of mutations at each position (blue: mutations decrease expression; orange: mutations increase expression). Absolute mutual information values are not directly comparable across promoters because barcode counts differ between promoters and conditions; see Section S3.2.

**Figure S14.**
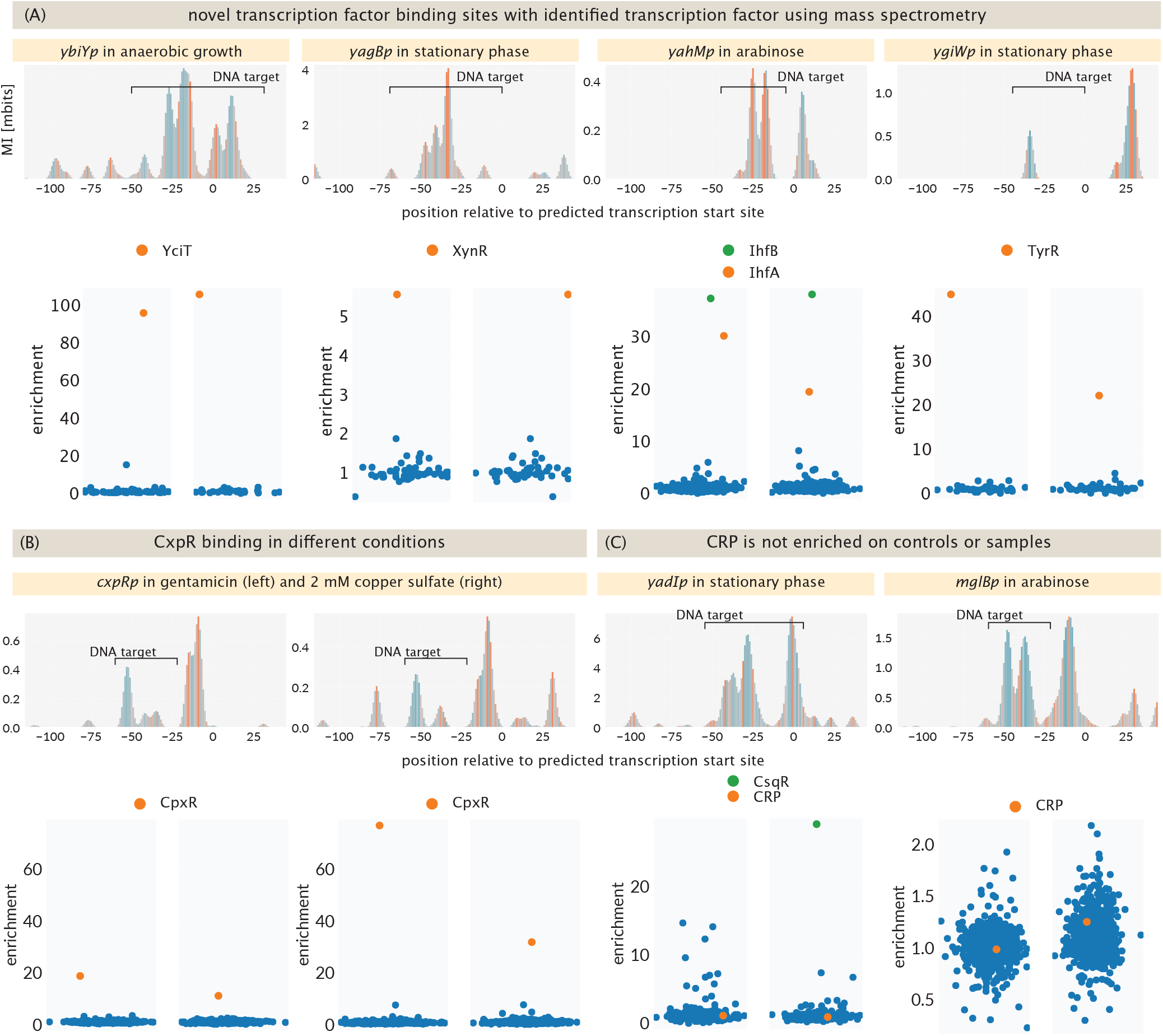
Information footprints paired with mass spectrometry enrichment for transcription factor identifications. Each promoter is shown as a footprint (top) with the bait sequence used for DNA chromatography indicated by the bracket labeled “DNA target,” and the corresponding mass spectrometry enrichment plot (bottom). The two columns within each enrichment plot are the two biological replicates. Enrichment is defined as in Section S2.7.3: a value of 1 (no specific binding) indicates that a protein occupies the same fraction of the recovered proteome on the target as on the control. Each dot represents one protein detected by mass spectrometry; the protein highlighted in orange is the primary candidate transcription factor identified for each bait, and a second highlighted hit (where present) is shown in green. (A) Novel binding sites with transcription factor identified by mass spectrometry: *ybiYp* in anaerobic growth (YciT), *yagBp* in stationary phase (XynR), *yahMp* in arabinose (IhfA, IhfB), and *ygiWp* in stationary phase (TyrR). (B) Confirmation of CpxR binding at *cpxRp* under gentamicin (left) and 2 mM copper sulfate (right). (C) Mass spectrometry does not recover CRP at the candidate *yadIp* bait (stationary phase) or at the *mglBp* positive control with a confirmed CRP binding site (arabinose). CRP is detected in both samples but is not enriched over the control bait. For *mglBp*, the left replicate was supplemented with cAMP and the right was not; the lack of enrichment in either case indicates that cAMP supplementation is not the limiting factor for CRP recovery in our setup. The full discussion of the CRP negative result and its implications for the *yadIp* CRP assignment is given in Section S5.2.

**Figure S15.**
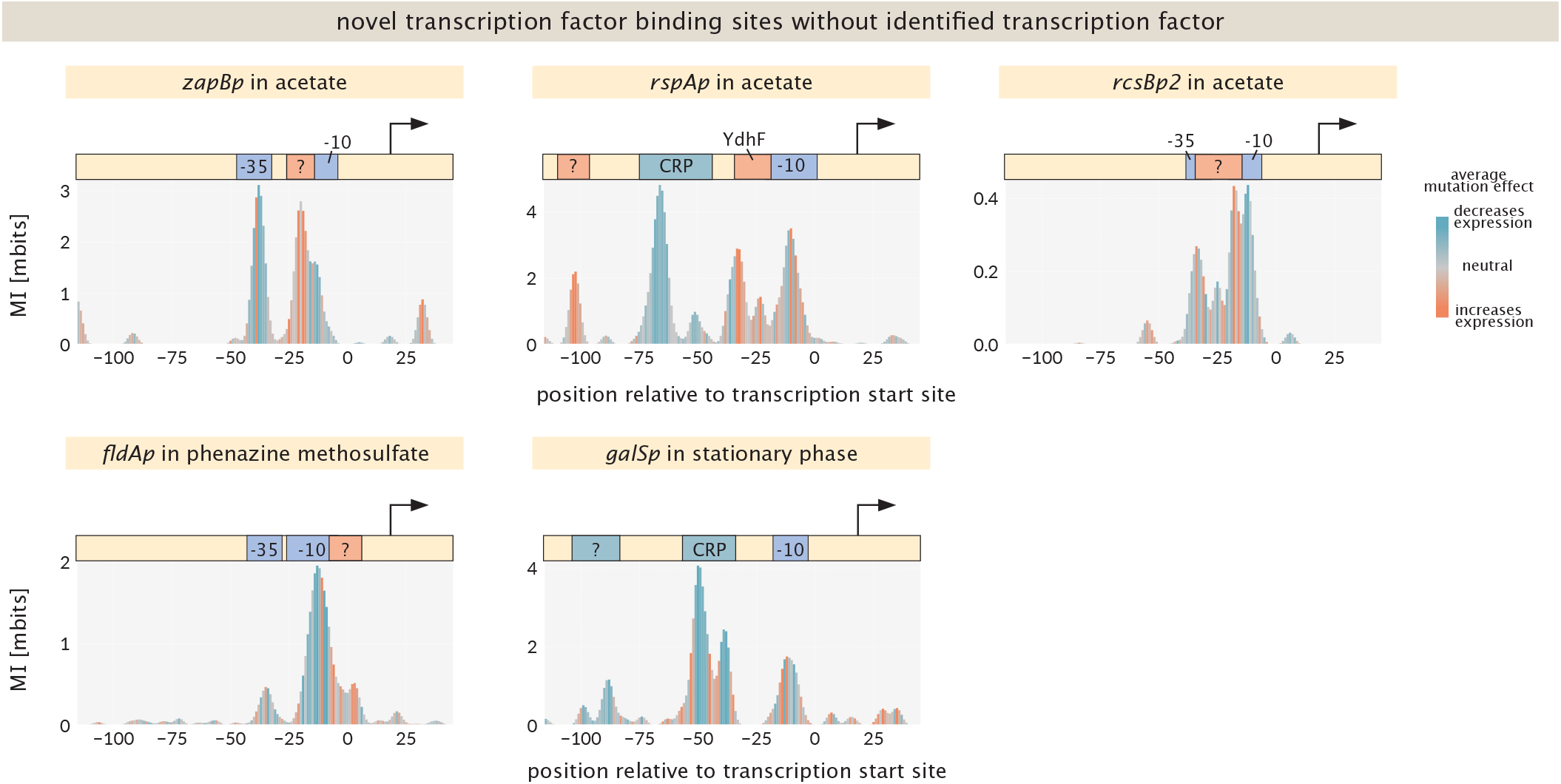
Information footprints for novel binding sites at which the responsible transcription factor could not be identified. Each panel shows the footprint for one promoter under the condition in which the novel signal is most pronounced. Boxes above the footprint indicate the architecture: known binding sites for annotated transcription factors (named), core promoter elements (−10, −35), and the position of the novel unidentified site (marked “?”). Color encodes the average effect of mutations as in Figure S13. The novel site at *rspAp* is the unidentified repressor at approximately −100; the known RspR and CRP sites are also visible at this promoter and are shown for context. The novel site at *galSp* is the unidentified activator at approximately −95; the known CRP site is also visible. For *zapBp, rcsBp2*, and *fldAp*, no annotated binding sites overlap the novel signals.

## Supplementary Tables

**Table S5.**
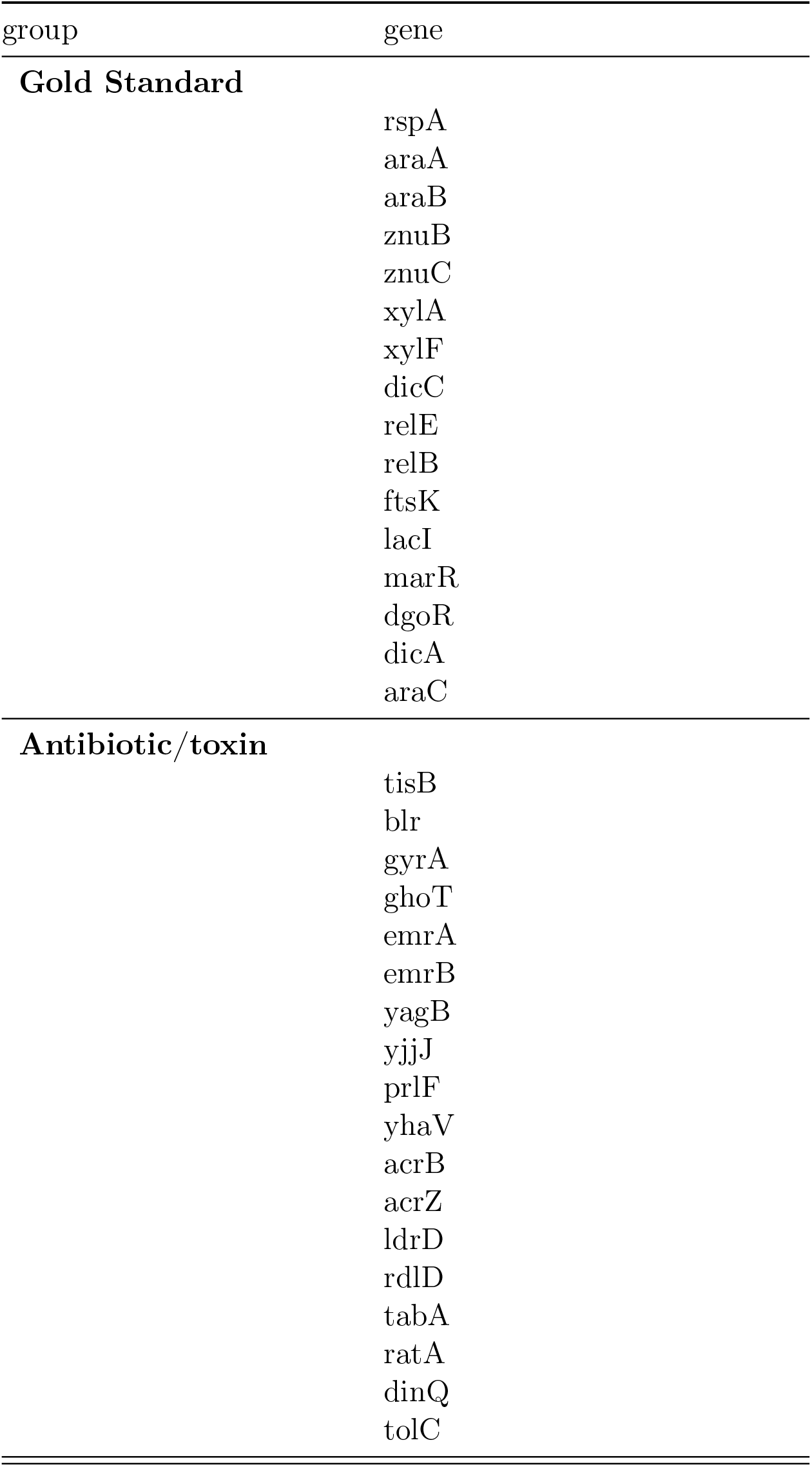
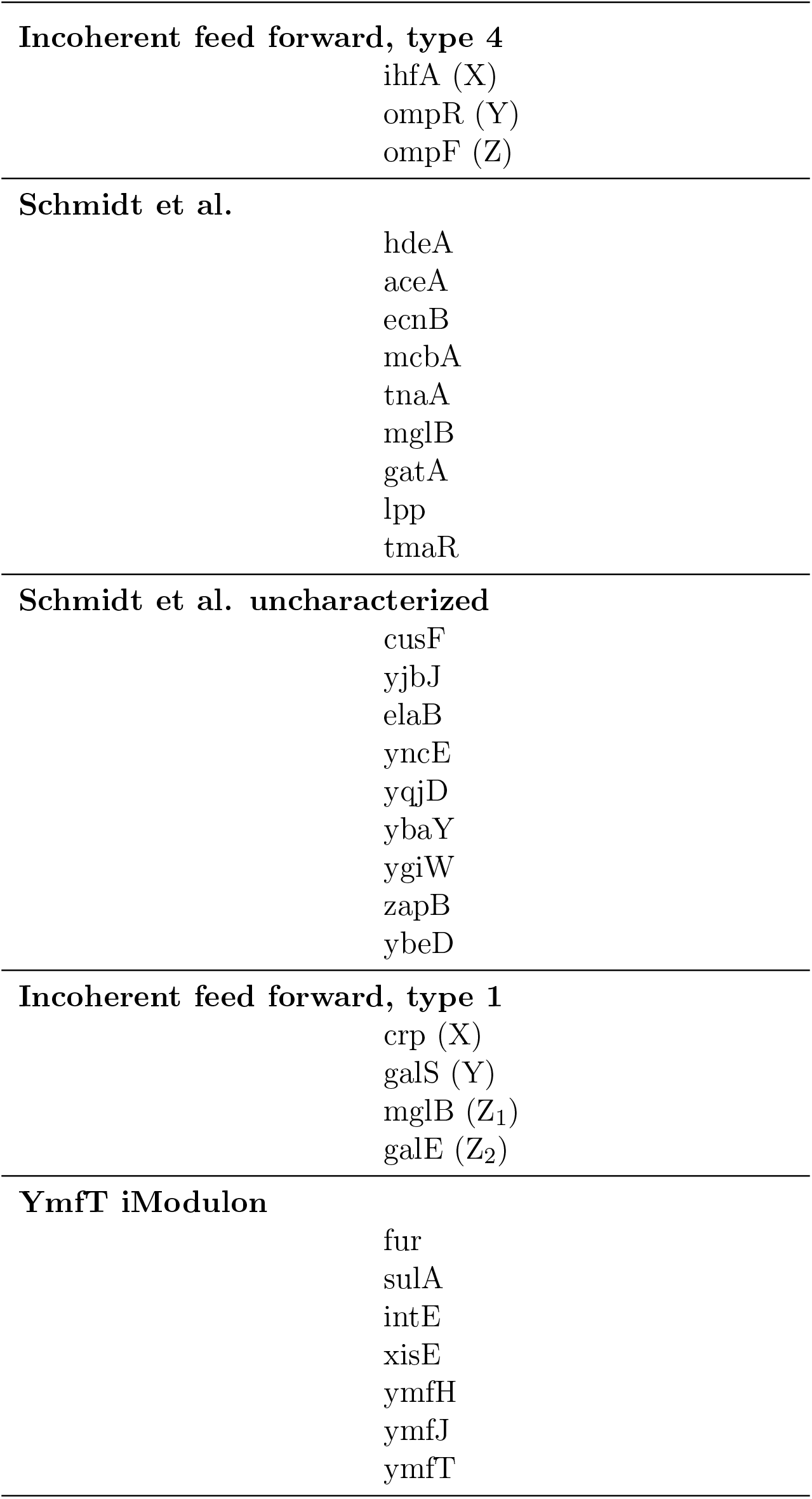
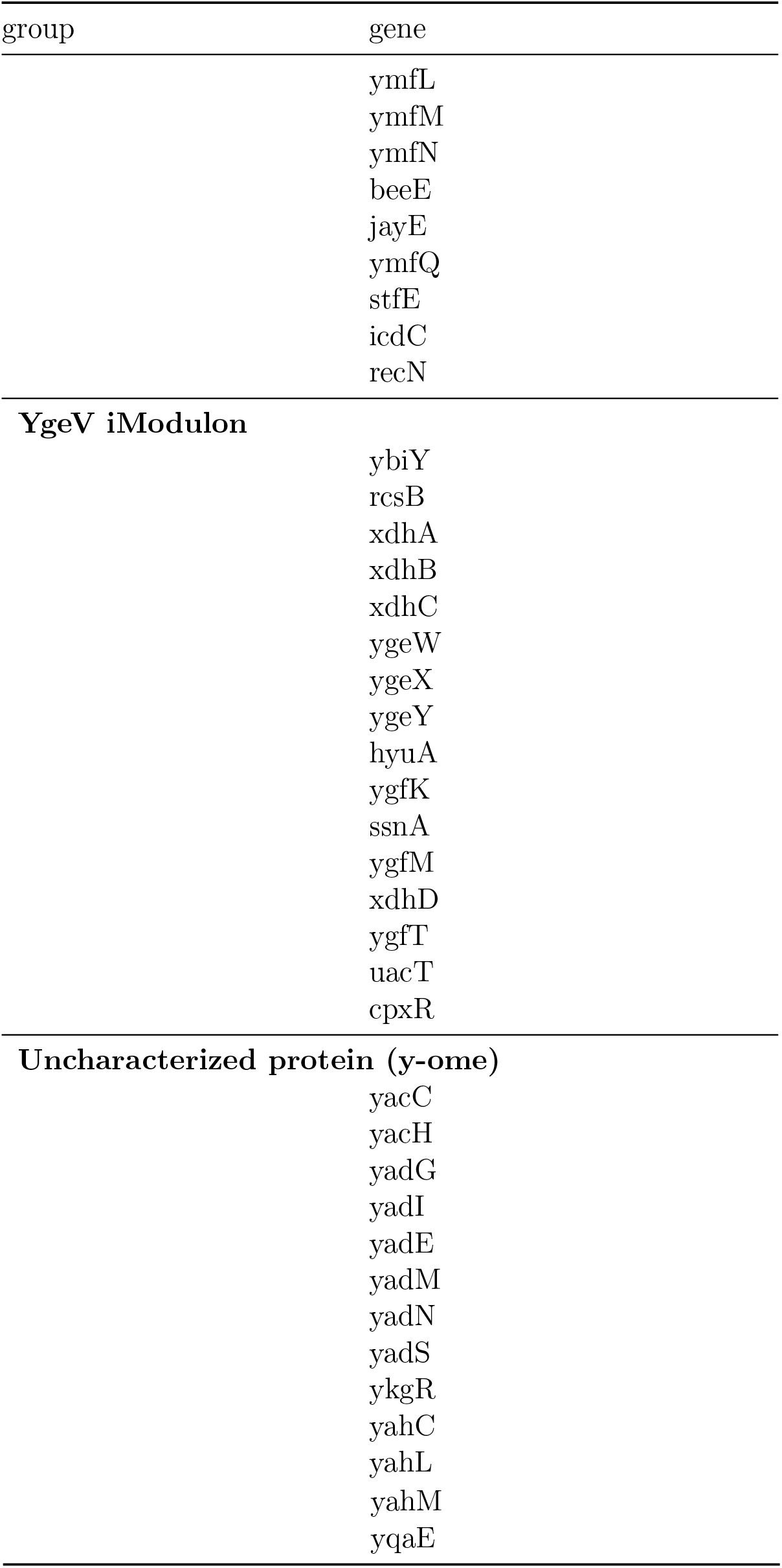
All genes chosen for this study.

**Table S6.**
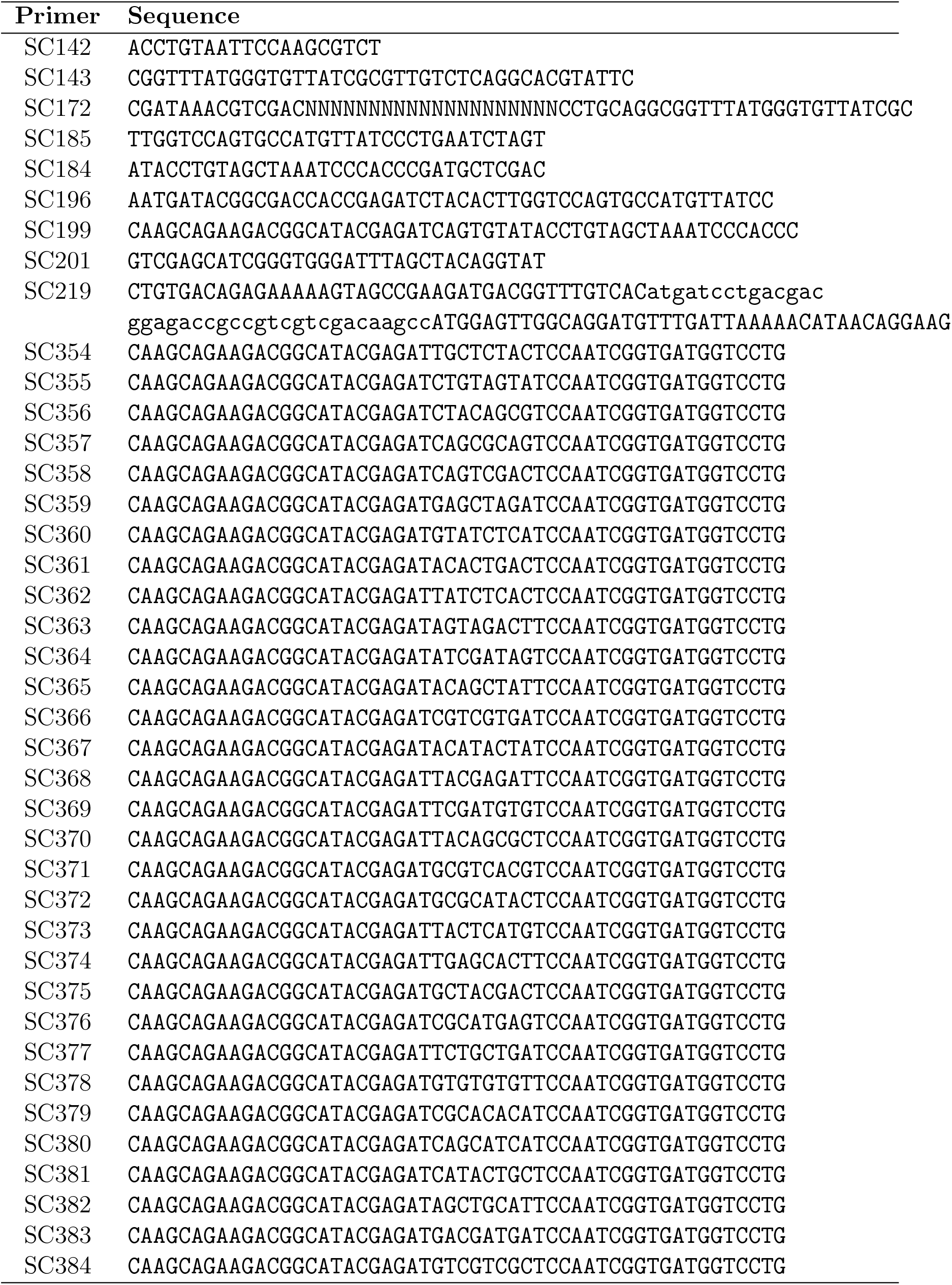
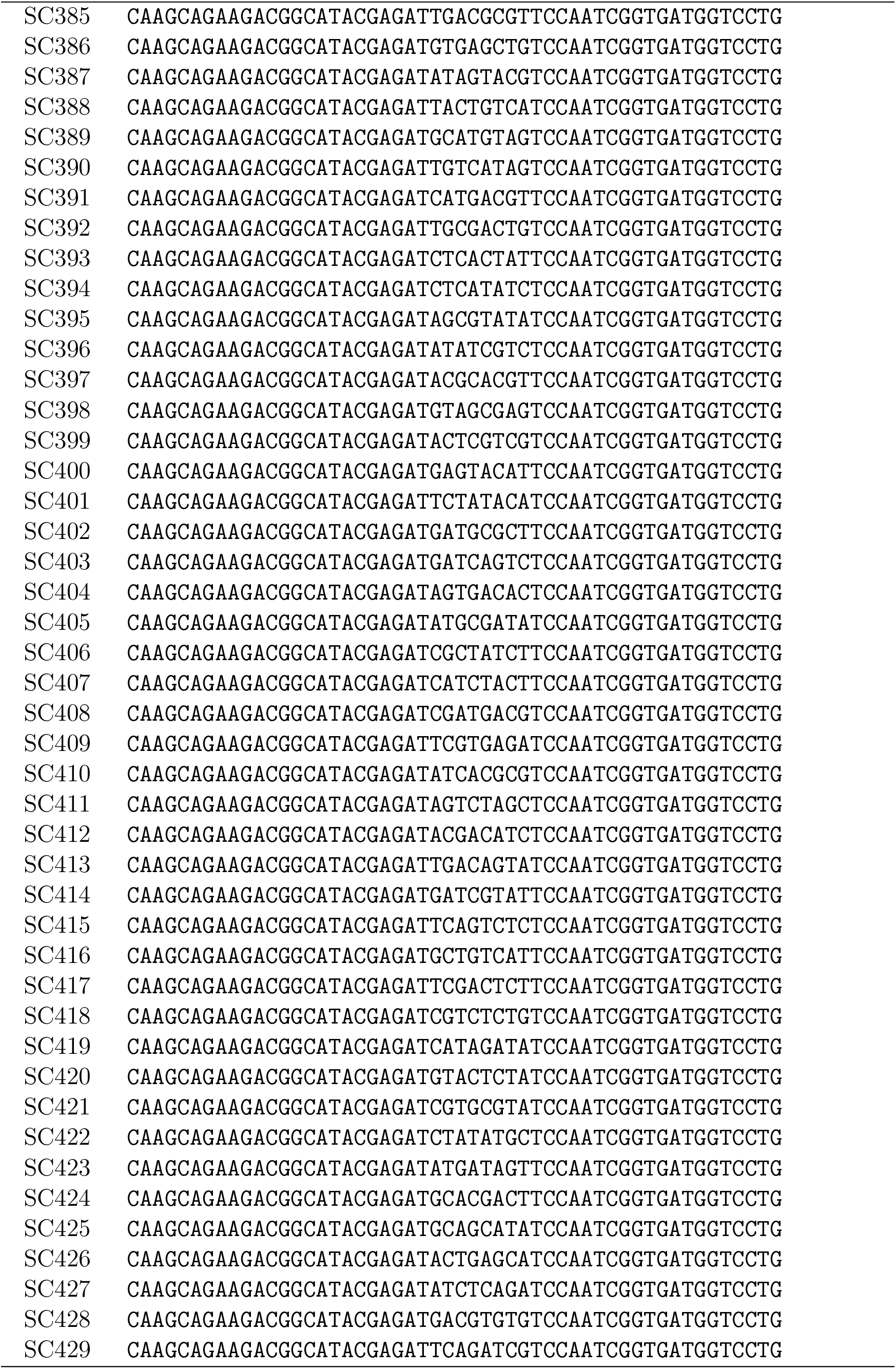
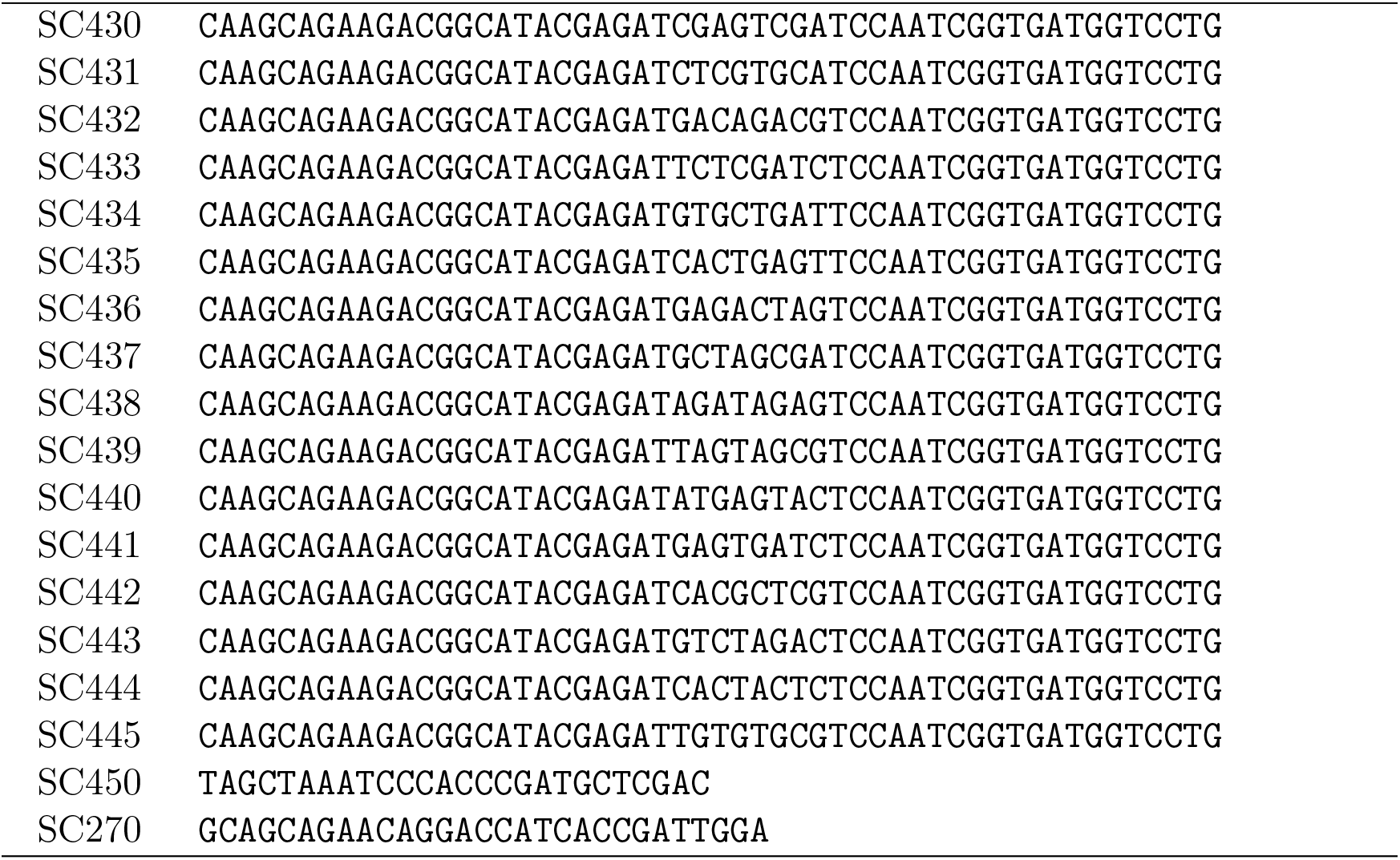
All primers and identifies used in this work.

**Table S7.** Growth Conditions for lysates used for mass spectrometry and associated supplements added. Notes: A supplement of 1 mM cAMP, a known co-factor for CRP binding, was used for our initial mass spectrometry runs of M9-glucose, M9-arabinose, stationary phase, and high osmolarity growth conditions, where we expected there to be annotated (e.g. *mglB* and *araB*) or predicted (e.g. *yadI*) binding sites of CRP. Since no significant enrichment for CRP was found for any of these runs, cAMP was not added to lysates for the other growth conditions. For high osmolarity, we based the concentration of supplemented salts on Figure 3 of Shabala et.al [118], which shows measurements of intracellular salt concentrations for growth in external media of varying NaCl concentrations.

| Growth Condition of Lysate | Final Concentration of Supplement(s) Added |
| --- | --- |
| M9-Glucose | 0.5% Glucose, 1 mM cAMP |
| M9-Arabinose | 0.5% Arabinose, 1 mM cAMP |
| M9-Xylose | 0.5% Xylose |
| LB | None Added |
| Stationary Phase (1d) | 1 mM cAMP |
| Leucine | 10 mM Leucine |
| Phenazine Methosulfate | 100 μM Phenazine Methosulfate |
| 2,2 Dipyridyl | 5 mM 2,2 Dipyridyl |
| Gentamicin | 5 mg/l Gentamicin |
| Copper Sulfate | 2 mM Copper Sulfate |
| Heat Shock | None added |
| H<sub>2</sub>O<sub>2</sub> | 2.5 mM H<sub>2</sub>O<sub>2</sub> |
| High Osmolarity (LB + 0.75 M NaCl) | 0.2 M NaCl, 0.15 M KCl, 1 mM cAMP |

**Table S8.** Liquid chromatography gradient parameters for mass spectrometry. Mobile Phase A contains 0.2% formic acid, 2% acetonitrile, and 97.8% water. Mobile Phase B contains 0.2% formic acid, 80% acetonitrile, and 19.8% water.

| Time | Duration | Flow (nl/min) | %B |
| --- | --- | --- | --- |
| 0:00 | 0:00 | 300 | 2 |
| 7:30 | 7:30 | 300 | 6 |
| 90:00 | 72:00 | 300 | 25 |
| 120:00 | 30:00 | 300 | 40 |
| 121:00 | 1:00 | 300 | 98 |
| 130:00 | 9:00 | 300 | 98 |

**Table S9.** Mass spectrometry scan settings for TMT samples.

| **Global Settings** |  |
| --- | --- |
| Ion source type | NSI |
| Spray voltage | 1500 V |
| Ion transfer tube temperature | 275 C |
| Polarity | Positive |
| **MS1 Scan Settings** |  |
| Resolution | 120000 |
| Normalized AGC target | 250 |
| Maximum IT | 50 msec |
| Scan range | 375-1600 m/z |
| **MS2 Scan Settings** |  |
| Resolution | 50000 |
| Normalized AGC target | Standard |
| Maximum IT | Dynamic |
| Loop time | 3 sec |
| Isolation window | 0.7 m/z |
| NCE | 35 |
| Spectrum data type | Centroid |
| Fixed first mass | 110 Z |

**Table S10.**
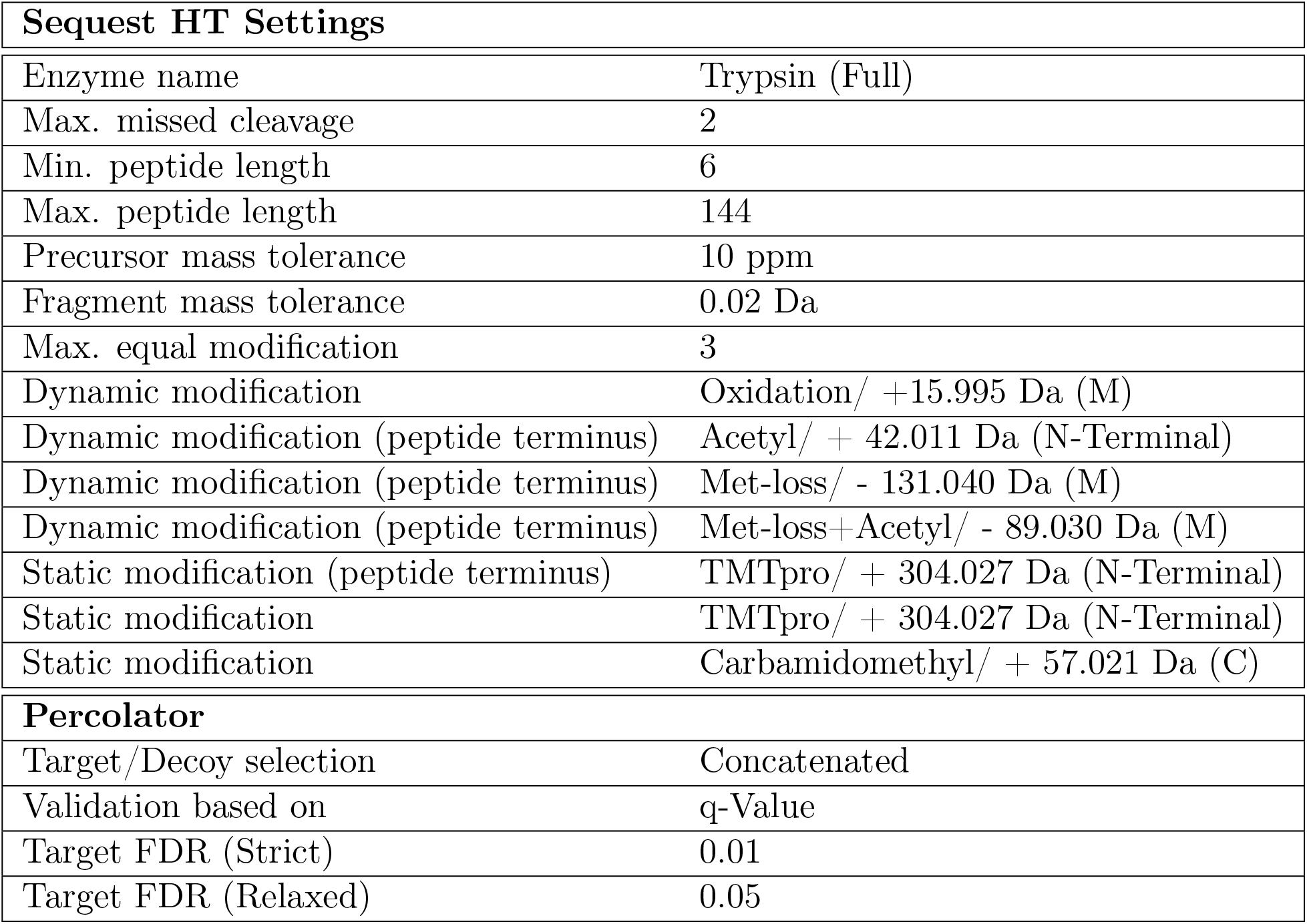
Search parameters for Protein Discoverer 2.5.

| **Sequest HT Settings** |  |
| --- | --- |
| Enzyme name | Trypsin (Full) |
| Max. missed cleavage | 2 |
| Min. peptide length | 6 |
| Max. peptide length | 144 |
| Precursor mass tolerance | 10 ppm |
| Fragment mass tolerance | 0.02 Da |
| Max. equal modification | 3 |
| Dynamic modification | Oxidation/ +15.995 Da (M) |
| Dynamic modification (peptide terminus) | Acetyl/ + 42.011 Da (N-Terminal) |
| Dynamic modification (peptide terminus) | Met-loss/ - 131.040 Da (M) |
| Dynamic modification (peptide terminus) | Met-loss+Acetyl/ - 89.030 Da (M) |
| Static modification (peptide terminus) | TMTpro/ + 304.027 Da (N-Terminal) |
| Static modification | TMTpro/ + 304.027 Da (N-Terminal) |
| Static modification | Carbamidomethyl/ + 57.021 Da (C) |
| **Percolator** |  |
| Target/Decoy selection | Concatenated |
| Validation based on | q-Value |
| Target FDR (Strict) | 0.01 |
| Target FDR (Relaxed) | 0.05 |

