## Supplementary File 1 for "Illuminating the uncharacterized regulatory genome of *E. coli* with massively parallel reporters"

### Contents

**Supplementary Information: Notes on Binding Site Annotations** 1
aceBp (aceBAK operon) 2
acrAp (acrAB operon) 3
acrZp (acrZ, small membrane protein associated with AcrB) 5
araBp (araBAD operon) 6
araCp (araC autoregulated promoter, divergent from araBp) 7
cpxRp and cpxRp2 (cpxR autoregulation, envelope stress response) 8
crpp1, crpp2, crpp3 (crp autoregulated promoter) 9
cusCp (cusCFBA copper/silver efflux operon) 10
dgoRp (dgoRKADT D-galactonate operon) 11
dicCp (dicB operon, DicA repression) 11
dinQp (dinQ, SOS-induced gene) 12
ecnBp (entericidin B lipoprotein) 12
elaBp (elaB C-tail anchored inner membrane protein) 13
fldAp (fldA, flavodoxin A) 14
ftsKp1 (ftsK, SOS-induced cell division gene) 14
furpa (fur, iron uptake regulator autoregulatory promoter) 15
furpb (fur, alternative TSS) 15
galEp1 (gal operon, upstream promoter) 16
galEp2 (gal operon, alternative TSS) 17
galSp (galS autoregulatory promoter) 17
gatYp (gat operon, galactitol utilization) 17
gyrAp (DNA gyrase A subunit) 18
hdeAp (hdeA, acid-inducible periplasmic chaperone) 18
ihfAp4 (ihfA, IHF alpha subunit, autoregulated) 20
marRp (marRAB operon, multiple antibiotic resistance) 20
mglBp (mglBAC high-affinity galactose transport operon) 22
mhpRp1 (mhpR, 3-(3-hydroxyphenyl)propionate degradation regulator) 22
mprAp (mprA/emrR, autorepressor of the emrRAB efflux operon) 22
ompFp (ompF, outer membrane porin F) 23
ompRp1 (ompR-envZ operon) 23
recNp (recN, SOS regulon, double-strand break repair) 25
relBp (relBE toxin-antitoxin operon) 26
rspAp (rspAB operon, putative D-mannose dehydrogenase) 27
sulAp (sulA, SOS-induced cell division inhibitor) 27
tisBp (tisB, SOS-induced toxin, type I toxin-antitoxin tisB/istR) 28
tnaCp (tnaLAB operon, tryptophanase, tryptophan degradation) 28
tolCp1, tolCp2, tolCp3, tolCp4 (tolC, outer membrane channel) 29
uofp (uof-fur operon, iron uptake regulator under oxidative stress) 31
xylAp (xylAB operon, D-xylose isomerase and kinase) 32
xylFp (xylFGHR operon, xylose ABC transporter and regulator) 33
ykgRp (ykgR, small protein of unknown function) 33
yncEp (yncE, iron-regulated periplasmic β-propeller protein) 34
yqaEp (yqaE, inner membrane protein of unknown function) 34
znuCp (znuCB operon, zinc ABC transporter) 35

### Supplementary Information: Notes on Binding Site Annotations

*This document was prepared with Claude (Anthropic) and the cited references were verified against PubMed.*

A working document tracking observations and caveats about individual RegulonDB annotations for binding sites at Reg-Seq promoters.

#### aceBp (aceBAK operon)

**RegulonDB annotations in our window:**

| Position | TF | Function | Confidence |
|---|---|---|---|
| −104 | IclR | repressor | S |
| −72 | IHF | activator | S |
| −53 | IclR | repressor | S |
| −38 | IclR | repressor | S |
| −30.5 | CRP | repressor | W |
| −26 | IclR | repressor | S |

**On the multiple IclR annotations.** RegulonDB lists four IclR positions, but these correspond to only two physical binding regions identified by DNase I footprinting. Yamamoto and Ishihama (2003) define:

- IclR box II between −52 and −19 (contains the −53, −38, and −26 RegulonDB entries)
- IclR box I between −125 and −99 (contains the −104 entry)

IclR binds as a tetramer to two dimeric 14–15 bp half-sites. The consensus palindrome is 5′-TGGAAATNATTTCCA-3′ (Pan et al. 1996; Molina-Henares et al. 2006). The three annotations within box II most likely represent the individual half-site recognition events of the tetramer rather than independent binding sites. For the purpose of counting sites, box I and box II are two physical binding events.

**What we observe.** We do not detect IclR binding at aceBp in any condition, including acetate where IclR repression is expected to be relieved and aceBAK induction should be strongest. In stationary phase we see a small repressor-like peak near −60, but it is upstream of IclR box II (−52 to −19), and the mutations that increase expression at that position could also create an improved −10 hexamer, so the signal is not specifically assignable to IclR. In addition, the aceBp reporter shows no substantial expression change across conditions relative to our non-functional control, so there is little dynamic range in which repressor footprints could be resolved.

**In vivo evidence that IclR represses aceBp.** Several lines of in vivo evidence support IclR repression of aceBAK: *iclR* loss-of-function mutants show constitutive aceBAK expression and are complemented in trans by wild-type *iclR* (Sunnarborg et al. 1990); point mutations within IclR box II reduce repression of an aceB::lacZ fusion in cells (Pan et al. 1996); and IHF activates aceB::lacZ under inducing conditions by opposing IclR repression (Resnik et al. 1996). The genetic evidence is sufficient to expect a Reg-Seq signal if the promoter were expressed and IclR were active.

**Possible reasons we do not see IclR.** Several non-exclusive explanations are consistent with our data: the native aceBp activity may not be recapitulated in our chromosomally integrated reporter, so mutagenesis cannot relieve a repressor that is not limiting in the construct; IclR box I (−125 to −99) extends beyond our upstream boundary at −115, so the box I contribution is partially lost — though Yamamoto and Ishihama (2003, Figure 6) show that box II alone still competes with RNA polymerase, so this cannot explain the absence of a box II footprint; and the physiological IclR ligand is not firmly established (Lorca et al. 2007 identified glyoxylate and pyruvate as antagonistic in vitro effectors, but the inducer responsible for aceBAK derepression on acetate in vivo remains unidentified).

**Other annotations in the window.** Besides the IclR boxes, RegulonDB lists two further TF entries at aceBp within our window:

- IHF at −72 (activator, S confidence). This entry is the IHFb footprint of Resnik et al. (1996), at −96 to −61 (an upstream IHFa site at −178 to −145 is outside our window). IHFb is immediately downstream of IclR box I. IHF at aceBp acts indirectly by relieving IclR repression rather than by direct contact with RNA polymerase (Resnik et al. 1996), so even if the promoter were active we would not expect a direct IHF-activator footprint here.

- CRP at −30.5 (repressor, W confidence). Sequence-similarity annotation only; no experimental evidence for CRP at aceBp in the literature we examined. Not a credible site.

**Accounting for the missed-sites tally.** RegulonDB lists six TF entries at aceBp in our window, but these do not correspond to six independent missed sites:

- The four IclR entries (−104, −53, −38, −26) are two physical binding events (boxes I and II); the three entries within box II reflect the half-site structure of tetrameric binding, not separate sites.
- IclR box I (−125 to −99) is only partially within our 160 bp window.
- IHF at −72 is only interpretable here as an activator via relief of IclR repression. In the absence of an IclR signal, an IHF contribution cannot be resolved, so the IHF entry is not an independent miss.
- CRP at −30.5 is based on sequence similarity alone (W confidence) and is not treated as a credible site (see above).

The honest accounting is that aceBp contributes **one missed experimentally supported binding site fully within our window: IclR box II.** Box I is a partial miss outside the window.

**References.**


- Yamamoto, K. and Ishihama, A. (2003). Two different modes of transcription repression of the *Escherichia coli* acetate operon by IclR. *Mol Microbiol* 47:183–194. PMID 12492863.
- Sunnarborg, A., Klumpp, D., Chung, T., and LaPorte, D.C. (1990). Regulation of the glyoxylate bypass operon: cloning and characterization of iclR. *J Bacteriol* 172:2642–2649. PMID 2185227.
- Pan, B., Unnikrishnan, I., and LaPorte, D.C. (1996). The binding site of the IclR repressor protein overlaps the promoter of aceBAK. *J Bacteriol* 178:3982–3984. PMID 8682810.
- Resnik, E., Pan, B., Ramani, N., Freundlich, M., and LaPorte, D.C. (1996). Integration host factor amplifies the induction of the aceBAK operon of *Escherichia coli* by relieving IclR repression. *J Bacteriol* 178:2715–2717. PMID 8626344.
- Lorca, G.L., Ezersky, A., Lunin, V.V., Walker, J.R., Altamentova, S., Evdokimova, E., Vedadi, M., Bochkarev, A., and Savchenko, A. (2007). Glyoxylate and pyruvate are antagonistic effectors of the *Escherichia coli* IclR transcriptional regulator. *J Biol Chem* 282:16476–16491. PMID 17426033.
- Molina-Henares, A.J. et al. (2006). Members of the IclR family of bacterial transcriptional regulators function as activators and/or repressors. *FEMS Microbiol Rev* 30:157–186. PMID 16472303.


#### acrAp (acrAB operon)

**RegulonDB annotations in our window:**

| Position | TF | Function | Confidence |
|---|---|---|---|
| −72.5 | MarA | activator | S |
| −72.5 | SoxS | activator | S |
| −22.5 | AcrR | repressor | C |
| −22.5 | EnvR/AcrS | repressor | S |
| −16 | MprA | repressor | W |
| +32 | PhoP | repressor | W |

**What we observe.** We detect a clear repressor footprint centered near −22.5. We cannot distinguish whether this footprint corresponds to AcrR or EnvR/AcrS because both proteins bind the same operator region (see below). We do not detect separate footprints at −16 (MprA) or +32 (PhoP). The marbox at −72.5 (MarA/SoxS/Rob) is not resolved in our data.

**AcrR and EnvR bind the same operator.** AcrR and EnvR (also called AcrS) are paralogous TetR-family repressors that bind the same operator at acrAB. The operator is a 24-bp inverted repeat 5′-TACATACATTTGTGAATGTATGTA-3′ overlapping the acrAB promoter, predicted by Rodionov et al. 2001 and confirmed for AcrR by fluorescence polarization with purified protein ($K_D \approx 20$ nM, two AcrR dimers cooperatively bound; Su et al. 2007). Hirakawa et al. 2008 demonstrated by DNase I footprinting that EnvR also binds the same 24-bp palindrome (protected region −118 to −89 relative to the acrA start codon), and showed by EMSA and overexpression assays that EnvR is a more potent repressor than AcrR (310-fold vs 2.4-fold acrA transcript reduction; ~3-fold higher binding affinity). The earlier identification of AcrR-dependent regulation by Ma et al. 1996 used gel shift with cell lysates and localized binding to the 180 bp acrR–acrAB intergenic region without base-pair resolution. Because the −22.5 entries for AcrR and EnvR correspond to a single physical operator, we count this as one site, not two: a single recovered footprint matches both annotations but cannot distinguish between them.

**Why Reg-Seq cannot distinguish AcrR from EnvR here.** Unlike the MarA/SoxS/Rob trio at acrZp, where each paralog has a distinct cognate inducer, AcrR and EnvR cannot be distinguished by condition in our panel. The known AcrR inducers (ethidium bromide, rhodamine 6G, proflavine, and elevated polyamines; Su et al. 2007; Harmon and Ruiz 2022) are not present in our 41 growth conditions, so AcrR is expected to remain bound throughout. EnvR's physiological inducer is unknown, and its native expression is at or below the detection threshold under standard laboratory conditions (Hirakawa et al. 2008). Based on expression levels alone, AcrR is the likely primary contributor to the observed −22.5 footprint, but our data do not formally exclude an EnvR contribution.

**MprA annotation is not supported by experimental evidence.** MprA (also called EmrR) is a MarR-family repressor of the *emrRAB* multidrug efflux operon, not of acrAB (Lomovskaya et al. 1995; Xiong et al. 2000). The direct experimental characterisation of MprA/EmrR binding is confined to the *emrR* promoter. We are not aware of any primary experimental demonstration that MprA binds the acrA promoter. The −16 annotation is W-confidence (sequence similarity only) and is consistent with an incidental match of the MarR-family recognition motif to a subsequence near the acrAp TSS; we do not treat this as a credible site.

**PhoP annotation is computational.** The PhoP entry at +32 is W-confidence and derives from the genome-wide position-weight-matrix scan of Zwir et al. (2005), which identified putative PhoP targets by combining expression clustering with a PhoP-box consensus model. PhoP/PhoQ has been reported to indirectly enhance acrAB expression, but we are not aware of a primary experimental demonstration that PhoP binds directly to the acrA promoter. We do not treat this as a credible site.

**MarA/SoxS/Rob marbox at −72.5.** The acrAB marbox is a well-characterised target of the mar/sox/rob regulon and a principal route by which salicylate, paraquat, and dipyridyl confer efflux-mediated antibiotic and solvent tolerance (Martin and Rosner 2002; White et al. 1997). The marbox binds MarA tightly in vitro but requires high MarA concentrations for half-saturation in vivo (Martin et al. 2008), so the effect size under physiological induction may be modest. We tested salicylate (MarA), PMS (SoxS), and dipyridyl (Rob) but did not resolve a marbox footprint in any of these conditions; we count each as an honest miss.

**Accounting for the missed-sites tally.** RegulonDB lists six TF entries at acrAp in our window, but these do not correspond to six independent sites:

- AcrR and EnvR/AcrS at −22.5 share a single 24-bp palindromic operator; one physical site, one recovered footprint, ambiguous assignment between two paralogs.
- MarA, SoxS, and Rob at −72.5 share a single marbox; one physical site, three regulatory interactions.
- MprA at −16 is W-confidence and not supported by primary experimental evidence at acrAp.
- PhoP at +32 is W-confidence and derives from a computational PWM scan, not experimental demonstration.

The honest accounting is that acrAp contains **two experimentally supported physical binding sites in-window: the AcrR/EnvR operator at −22.5 (recovered) and the MarA/SoxS/Rob**

**marbox at −72.5 (not recovered).** Counted as physical sites: **1/2 recovered.** Counted as regulatory interactions (three paralog activators at the shared marbox): **1/4 recovered** (AcrR/EnvR operator plus three marbox misses).

**References.**


- Ma, D., Alberti, M., Lynch, C., Nikaido, H., and Hearst, J.E. (1996). The local repressor AcrR plays a modulating role in the regulation of acrAB genes of *Escherichia coli* by global stress signals. *Mol Microbiol* 19:101–112. PMID 8821940.
- Rodionov, D.A., Gelfand, M.S., Mironov, A.A., and Rakhmaninova, A.B. (2001). Comparative approach to analysis of regulation in complete genomes: multidrug resistance systems in gamma-proteobacteria. *J Mol Microbiol Biotechnol* 3:319–324. PMID 11321589.
- Su, C.C., Rutherford, D.J., and Yu, E.W. (2007). Characterization of the multidrug efflux regulator AcrR from *Escherichia coli*. *Biochem Biophys Res Commun* 361:85–90. PMID 17644067.
- Hirakawa, H., Takumi-Kobayashi, A., Theisen, U., Hirata, T., Nishino, K., and Yamaguchi, A. (2008). AcrS/EnvR represses expression of the acrAB multidrug efflux genes in *Escherichia coli*. *J Bacteriol* 190:6276–6279. PMID 18567659.
- Lomovskaya, O., Lewis, K., and Matin, A. (1995). EmrR is a negative regulator of the *Escherichia coli* multidrug resistance pump EmrAB. *J Bacteriol* 177:2328–2334. PMID 7730261.
- Xiong, A., Gottman, A., Park, C., Baetens, M., Pandza, S., and Matin, A. (2000). The EmrR protein represses the *Escherichia coli* emrRAB multidrug resistance operon by directly binding to its promoter region. *Antimicrob Agents Chemother* 44:2905–2907. PMID 10991887.
- Zwir, I. et al. (2005). Dissecting the PhoP regulatory network of *Escherichia coli* and *Salmonella enterica*. *Proc Natl Acad Sci USA* 102:2862–2867. PMID 15703297.
- Martin, R.G. and Rosner, J.L. (2002). Genomics of the marA/soxS/rob regulon of *Escherichia coli*: identification of directly activated promoters by application of molecular genetics and informatics to microarray data. *Mol Microbiol* 44:1611–1624. PMID 12067348.
- Martin, R.G., Bartlett, E.S., Rosner, J.L., and Wall, M.E. (2008). Activation of the *Escherichia coli* marA/soxS/rob regulon in response to transcriptional activator concentration. *J Mol Biol* 380:278–284. PMID 18514222.
- White, D.G., Goldman, J.D., Demple, B., and Levy, S.B. (1997). Role of the acrAB locus in organic solvent tolerance mediated by expression of marA, soxS, or robA in *Escherichia coli*. *J Bacteriol* 179:6122–6126. PMID 9324261.
- Harmon, D.E. and Ruiz, C. (2022). The multidrug efflux regulator AcrR of *Escherichia coli* responds to exogenous and endogenous ligands to regulate efflux and detoxification. *mSphere* 7:e00474-22. PMID 36416552.


#### acrZp (acrZ, small membrane protein associated with AcrB)

**RegulonDB annotations in our window:**

| Position | TF | Function | Confidence |
|---|---|---|---|
| −40.5 | MarA | activator | W |
| −40.5 | Rob | activator | W |
| −40.5 | SoxS | activator | W |

**What we observe.** We detect an activator footprint at −40.5 in three distinct conditions, each corresponding to the cognate inducer of one of the three paralogs: 2,2′-dipyridyl (Rob), sodium salicylate (MarA), and phenazine methosulphate (SoxS). The footprint at −40.5 is thus recovered in all three conditions, with condition-specific assignment of the responsible paralog.

**One marbox, three paralogs, three regulatory events.** MarA, SoxS, and Rob are paralogous AraC/XylS-family activators that share a common ~20-bp degenerate recognition sequence, the marbox (Martin et al. 1999; Martin and Rosner 2002). At acrZp, RegulonDB lists three entries at −40.5, one for each paralog. Structurally these correspond to a single physical marbox. The three paralogs are known to respond to distinct signals: MarA is derepressed by sodium salicylate (via

MarR inactivation), SoxS is induced by superoxide generators such as phenazine methosulphate (via SoxR), and Rob is activated by 2,2′-dipyridyl (Rosner et al. 2002). In our data, the −40.5 activator footprint is specific to the cognate inducing condition of each paralog, indicating that in vivo each paralog does engage the marbox under its own inducing condition rather than all three binding interchangeably. We therefore count acrZp as contributing three recovered regulatory interactions at one physical site.

**Class II marbox.** The −40.5 position overlaps the −35 hexamer, consistent with a class II marbox (Martin et al. 1999). In vitro binding affinity correlates only weakly with in vivo activation efficiency for marboxes (Martin et al. 2008), so class designation alone does not predict footprint strength.

**Evidence basis for acrZ as a Mar/Sox/Rob target.** The assignment of acrZ to the Mar/Sox/Rob regulon is based on expression responses to regulator overexpression rather than on direct biochemistry at acrZp itself: Hobbs et al. (2012) established that acrZ (formerly ybhT) is co-induced with acrAB under MarA, SoxS, and Rob activation and identified a class II marbox upstream of acrZ matching the established consensus (Martin et al. 1999; Martin and Rosner 2002). We are not aware of a direct EMSA or footprinting demonstration of MarA/SoxS/Rob binding at acrZp specifically. Our data provide the in-vivo, condition-specific evidence that each paralog engages the acrZp marbox under its cognate inducing condition.

**Accounting for the missed-sites tally.** RegulonDB lists three TF entries at acrZp in our window, all at the same position (−40.5). These correspond to one physical marbox recognised by three paralogous activators. Under the framing of this accounting, each TF–site interaction is a distinct regulatory event: the marbox is recovered in each of the three cognate inducing conditions (dipyridyl, salicylate, PMS), providing in vivo evidence for each paralog at the same operator. We count acrZp as contributing **three recovered regulatory interactions at one physical binding site.**

**References.**


- Hobbs, E.C., Yin, X., Paul, B.J., Astarita, J.L., and Storz, G. (2012). Conserved small protein associates with the multidrug efflux pump AcrB and differentially affects antibiotic resistance. *Proc Natl Acad Sci USA* 109:16696–16701. PMID 23010927.
- Martin, R.G., Gillette, W.K., Rhee, S., and Rosner, J.L. (1999). Structural requirements for marbox function in transcriptional activation of mar/sox/rob regulon promoters in *Escherichia coli*: sequence, orientation and spatial relationship to the core promoter. *Mol Microbiol* 34:431–441. PMID 10564485.
- Martin, R.G. and Rosner, J.L. (2002). Genomics of the marA/soxS/rob regulon of *Escherichia coli*: identification of directly activated promoters by application of molecular genetics and informatics to microarray data. *Mol Microbiol* 44:1611–1624. PMID 12067348.
- Martin, R.G., Bartlett, E.S., Rosner, J.L., and Wall, M.E. (2008). Activation of the *Escherichia coli* marA/soxS/rob regulon in response to transcriptional activator concentration. *J Mol Biol* 380:278–284. PMID 18514222.
- Rosner, J.L., Dangi, B., Gronenborn, A.M., and Martin, R.G. (2002). Posttranscriptional activation of the transcriptional activator Rob by dipyridyl in *Escherichia coli*. *J Bacteriol* 184:1407–1416. PMID 11844771.


---

> **Note (verification status):** The aceBp, acrAp, and acrZp sections above have been audited at full claim-verification level against the cited primary sources. Sections from araBp onward (below) are drafts pending the same audit; references and specific claims have not yet been confirmed against the underlying papers.

---

#### araBp (araBAD operon)

The Schleif light-switch mechanism (Schleif 2010) requires AraC to bridge araI1 (−64) and araO2 (~−270, RegulonDB position −275) into a ~210 bp repressive loop in the absence of arabinose, and to occupy araI1+araI2 (−43) as an activator in its presence (Lobell and Schleif 1990). Because araO2 lies ~150 bp outside our 160 bp window, the loop cannot form in our construct and the loop-mediated repression mode is not assayable; only the arabinose-induced activation mode is in window.

| Position | TF | Function | Observed? | Reason / notes |
|---|---|---|---|---|
| −93.5 | CRP | activator | weak | Loop-relief is the dominant CRP role at native araBp (Hahn et al. 1984); loop is absent here, leaving only the smaller direct-activation component (Zhang and Schleif 1998). |
| −64 | AraC | repressor | no | Loop partner araO2 outside window; uninduced baseline near background, no dynamic range for a repressor footprint. |
| −64 | AraC | activator | yes (arab.) | araI1 in the AraC dimer activator complex. |
| −43 | AraC | activator | yes (arab.) | araI2 in the AraC dimer activator complex. |

RegulonDB also lists AraC repressor sites at −117 (window edge), −138, and −275 (araO2); all outside or at the boundary, excluded under loop-outside-window.

**Accounting.** 3/4 in-window functions recovered (CRP weakly). The repressor function at −64 is not recovered but is not counted as a separate missed site: the same physical position is recovered as the activator, and the silent uninduced baseline precludes repressor detection here.

**References.**


- Schleif, R. (2010). *FEMS Microbiol Rev* 34:779–796. PMID 20491933.
- Lobell, R.B. and Schleif, R.F. (1990). *Science* 250:528–532. PMID 2237403.
- Hahn, S., Dunn, T., and Schleif, R. (1984). *J Mol Biol* 180:61–72. PMID 6392569.
- Zhang, X. and Schleif, R. (1998). *J Bacteriol* 180:195–200. PMID 9440505.


#### araCp (araC autoregulated promoter, divergent from araBp)

araCp shares its regulatory DNA with araBp; the same physical CRP site (araBp −93.5 / araCp −54.5) and AraC binding region serve both, in opposite directions. Hamilton and Lee (1988) showed that araC autoregulation requires three AraC sites — araI1, araO1, and araO2 — only one of which (the araO1 region, overlapping the araCp −35/−10) is fully contained in our window; araI1 and araO2 lie outside, so the cooperative autorepression complex cannot form. The −19.5/+2.5 XylR sites are the EMSA-confirmed XylR binding region in the araC/araB intergenic DNA (Koirala et al. 2016).

| Position | TF | Function | Observed? | Reason / notes |
|---|---|---|---|---|
| −105 | AraC | activator | no | Upstream half-site of an AraC binding pair; cooperative loop partner araI1 outside window. |
| −84 | AraC | rep+act | no | Same pair as −105; same reason. |
| −54.5 | CRP | activator | yes | Same physical CRP site as araBp −93.5; cleaner here because araCp lacks the strong AraC-driven activation that compresses dynamic range at araBp. |
| −31 | AraC | repressor | no | Half-site of araO1 overlapping the araCp −35; full autorepression complex requires araI1 and araO2, both outside window. |
| −19.5 | XylR | repressor | no | Honest miss; xylose-dependent EMSA binding (Koirala et al. 2016) does not produce a Reg-Seq footprint here. |
| −10 | AraC | repressor | no | Half-site of araO1 overlapping the araCp −10; same reason as −31. |
| +2.5 | XylR | repressor | no | Honest miss; same EMSA evidence as −19.5. |

**Accounting.** 1/7 in-window annotations recovered (CRP at −54.5). Four AraC sites (−105, −84, −31, −10) are excluded under loop-outside-window — all four belong to autoregulation complexes that require partner sites outside our window. The two XylR sites (−19.5, +2.5) are honest misses. **1/3 testable recovered.**

**References.**


- Lee, N.L., Gielow, W.O., and Wallace, R.G. (1981). *Proc Natl Acad Sci USA* 78:752–756. PMID 6262769.
- Hamilton, E.P. and Lee, N. (1988). *Proc Natl Acad Sci USA* 85:1749–1753. PMID 3279415.
- Koirala, S., Wang, X., and Rao, C.V. (2016). *J Bacteriol* 198:386–393. PMID 26527647.


#### cpxRp and cpxRp2 (cpxR autoregulation, envelope stress response)

**RegulonDB lists CpxR sites at −47 (S) and −67 (W).** Both are annotated separately under cpxRp and cpxRp2, but both promoters share the same genomic coordinate in our reporter, so the physical sites are just two (−47 and −67); we do not double-count.

| Position | TF | Function | Observed? | Reason / notes |
|---|---|---|---|---|
| −47 | CpxR | activator | yes | High-confidence site, recovered. |
| −67 | CpxR | activator | no | W-confidence ChIP-exo annotation only; ChIP-exo cannot resolve two CpxR dimer footprints 20 bp apart, so −67 is most parsimoniously a peak-caller artifact on the same signal supporting −47, not an independent binding event. |

**Accounting.** 1/1 credible physical sites recovered (−67 not counted as a separate site).

#### crpp1, crpp2, crpp3 (crp autoregulated promoter)

RegulonDB lists the same physical sites multiple times under three alternative TSS (crpp1/2/3); only crpp1 is active in our data, so we use crpp1 numbering and do not double-count. The Fis site at +69 is downstream of our boundary and excluded, leaving 6 in-window sites. CRP is recovered well at many other promoters in our dataset, so the failure here is promoter-specific.

| Position | TF | Function | Observed? | Reason / notes |
|---|---|---|---|---|
| −92 | Fis | repressor | no | Fis-degenerate exclusion (no Fis site is recovered anywhere in our panel; Hübner and Arber 1989). |
| −59.5 | CRP | activator | no | Honest miss; this is CRP II (González-Gil et al. 1998), a modest activator at this promoter; small contribution likely compressed by the FIS-dominated regulatory background here. |
| −45.5 | Cra | activator | no | Honest miss; tested in acetate (condition 39) where Cra is active. Possible interference from nearby CRP II footprint (14 bp away) and downstream FIS/CRP sites. |
| +31 | Fis | repressor | no | Fis-degenerate exclusion. |

| Position | TF | Function | Observed? | Reason / notes |
|---|---|---|---|---|
| +41.5 | CRP | repressor | no | This is CRP I; González-Gil et al. (1998) place it in a complex repression mechanism involving cooperative FIS binding (notably at +68); the assay does not capture the full nucleoprotein context. |
| +49 | Fis | repressor | no | Fis-degenerate exclusion. |

**Accounting.** 0/6 recovered. Three Fis sites excluded under Fis-degenerate. CRP at +41.5 is excluded as a complex nucleoprotein-mechanism site outside the assay's reach. CRP at −59.5 and Cra at −45.5 are honest misses. **0/3 testable recovered.**

**References.**


- González-Gil, G., Kahmann, R., and Muskhelishvili, G. (1998). *EMBO J* 17:2877–2885. PMID 9582281.
- Hübner, P. and Arber, W. (1989). *EMBO J* 8:577–585. PMID 2656257.


#### cusCp (cusCFBA copper/silver efflux operon)

RegulonDB lists three activator annotations at the same physical position (~−53.5/−54) for CusR, HprR, and PhoB — three OmpR-family response regulators sharing a recognition consensus. The three annotations correspond to one physical site recognised by three paralogs, each tested under its cognate condition. The promoter is essentially silent in HOCl and phosphate starvation, so absence of expression in those conditions is itself evidence that the respective paralog is not functionally engaging the site here.

| Position | TF | Function | Observed? | Reason / notes |
|---|---|---|---|---|
| −53.5 | CusR | activator | yes (Cu) | Recovered in copper sulfate; matches Munson et al. 2000 cusRS copper-responsive TCS. |
| −54 | PhoB | activator | no | Promoter silent in phosphate starvation; PhoB annotation is most likely a sequence-match assignment by paralog consensus, not a functional regulatory interaction in our conditions. |

| Position | TF | Function | Observed? | Reason / notes |
|---|---|---|---|---|
| −53.5 | HprR | activator | no | Promoter silent in HOCl (4 mM); HprR annotation likely sequence-match (Urano et al. 2015 demonstrated CusR/YedW share recognition at *hiuH*, which is the basis for the paralog cross-assignment). |

**Accounting.** One physical site, three annotated interactions. **1/3 recovered** (CusR in copper). HprR and PhoB annotations are not supported as functional in our conditions.

**References.**


- Munson, G.P., Lam, D.L., Outten, F.W., and O'Halloran, T.V. (2000). *J Bacteriol* 182:5864–5871. PMID 11004187.
- Urano, H., Umezawa, Y., Yamamoto, K., Ishihama, A., and Ogasawara, H. (2015). *Microbiology (Reading)* 161:729–738. PMID 25568260.


#### dgoRp (dgoRKADT D-galactonate operon)

The RegulonDB-annotated TSS for dgoRp is not active in our data; an upstream alternative TSS is active instead, consistent with the multiple overlapping RNAP binding sites identified by Belliveau et al. (2018). Under the active-TSS coordinates, two of the three annotated sites fall outside the window.

| Position | TF | Function | Observed? | Reason / notes |
|---|---|---|---|---|
| −117 | CRP | activator | no | Outside window under re-mapped active TSS coordinates. |
| −110 | DgoR | repressor | no | Outside window under re-mapped active TSS coordinates. |
| −88.5 | DgoR | repressor | yes | In-window; matches DgoR binding identified at this promoter by mass spec (Belliveau et al. 2018). |

**Accounting.** **1/1 in-window site recovered.** Two annotations excluded as out-of-window under the re-mapped active TSS.

**References.**


- Belliveau, N.M. et al. (2018). *PNAS* 115:E4796–E4805. PMID 29728462.


#### dicCp (dicB operon, DicA repression)

| Position | TF | Function | Observed? | Reason / notes |
|---|---|---|---|---|
| +2.5 | DicA | repressor | yes (indirect) | No information footprint, but elevated mutation rate inside the operator during cloning indicates negative selection: DicA expressed from plasmid library titrates from its chromosomal site, imposing a fitness defect on operator-intact variants. By assay time, surviving variants carry already-disrupted operators, so no expression shift is resolvable. Discussed in detail elsewhere in the SI. |

**Accounting. 1/1 recovered** (operator function confirmed by the cloning-stage selection signature).

#### dinQp (dinQ, SOS-induced gene)

| Position | TF | Function | Observed? | Reason / notes |
|---|---|---|---|---|
| −29.5 | LexA | repressor | no | Position overlaps the RNAP −35 region; under SOS-inducing conditions LexA signal here would be inseparable from sigma binding. Also W-confidence — may simply have weaker intrinsic affinity than the −7.5 site. |
| −7.5 | LexA | repressor | yes | Recovered. |

**Accounting. 1/2 recovered.**

#### ecnBp (entericidin B lipoprotein)

| Position | TF | Function | Observed? | Reason / notes |
|---|---|---|---|---|
| +18.5 | OmpR | repressor | yes (weak) | Weak but reproducible footprint at +18.5 in some conditions; ecnB is σS-dependent and derepressed in *ompR* mutants (Bishop et al. 1998), but Bishop et al. only inferred the +18.5 site by sequence similarity to the OmpR consensus (no direct binding assay). Our footprint provides the first direct functional evidence that OmpR engages this specific sequence. |

**Accounting. 1/1 weakly recovered.** Condition-dependent signal consistent with a modest repressive input operating against a strong σS-driven activation.

**References.**


- Bishop, R.E., Leskiw, B.K., Hodges, R.S., Kay, C.M., and Weiner, J.H. (1998). *J Mol Biol* 280:583–596. PMID 9677290.


#### elaBp (elaB C-tail anchored inner membrane protein)

Both annotations sit at the same physical position (one binding site, two regulatory functions). Guo et al. (2019) demonstrated direct OxyR binding here by EMSA and showed OxyR acts as activator in exponential phase (RpoS-dependent) and as repressor when overproduced in stationary phase.

| Position | TF | Function | Observed? | Reason / notes |
|---|---|---|---|---|
| −31 | OxyR | repressor | yes | Robust footprint across galactose, acetate, xylose, ethanol — all conditions of high respiratory flux relative to glucose, which elevates endogenous H2O2 (Seaver and Imlay 2001) sufficient to engage OxyR's repressor mode without exogenous peroxide. |

| Position | TF | Function | Observed? | Reason / notes |
|---|---|---|---|---|
| −31 | OxyR | activator | no | Honest miss; tested under exogenous H2O2 (2.5 mM/10 min and 0.1 mM/30 min, exponential phase); regimes may saturate too briefly or be efficiently scavenged by basal OxyR regulon before reporter library imprint. |

**Accounting.** One physical site, two regulatory functions. **1/2 recovered** (repressor robust, activator missed under exogenous H2O2).

**References.**


- Guo, Y., Li, Y., Zhan, W., Wood, T.K., and Wang, X. (2019). *Microb Biotechnol* 12:392–404. PMID 30656833.
- Seaver, L.C. and Imlay, J.A. (2001). *J Bacteriol* 183:7173–7181. PMID 11717276.


#### fldAp (fldA, flavodoxin A)

| Position | TF | Function | Observed? | Reason / notes |
|---|---|---|---|---|
| −61.5 | SoxS | activator | no | Honest miss under PMS (cognate SoxS inducer); annotation is from low-affinity Mar-box overexpression target set (Martin et al. 2008), so SoxS may not productively engage at endogenous induction. (We separately detect an unidentified repressor signal at −102 under PMS, discussed in the novel-sites section.) |

**Accounting. 0/1 recovered.**

**References.**


- Martin, R.G., Bartlett, E.S., Rosner, J.L., and Wall, M.E. (2008). *J Mol Biol* 380:278–284. PMID 18514222.


#### ftsKp1 (ftsK, SOS-induced cell division gene)

| Position | TF | Function | Observed? | Reason / notes |
|---|---|---|---|---|
| −0.5 | LexA | repressor | yes (weak) | Operator overlaps the TSS (standard LexA repression arrangement at SOS-inducible promoters); footprint weaker than at sulAp/tisBp, likely reflecting lower-affinity operator. |

**Accounting. 1/1 recovered.**

#### furpa (fur, iron uptake regulator autoregulatory promoter)

The two Fur annotations at −7 and −1 are 6 bp apart and almost certainly correspond to a single Fur dimer operator (tandem half-sites of the ~19 bp Fur box); we count them as one physical site. Two physical sites in total.

| Position | TF | Function | Observed? | Reason / notes |
|---|---|---|---|---|
| −77.5 | CRP | activator | no | Honest miss; canonical class-I CRP activator position, no footprint detected in any CRP-favoring condition. |
| −7, −1 | Fur | repressor | no | Honest miss; Fur should be bound (no iron-limitation condition in our panel) and a uniform repression should still produce a mutational footprint. The positions sit in the −10/TSS spacer where mutations have very large RNAP-engagement effects, likely dominating any Fur-disruption signal. |

**Accounting. 0/2 recovered.** Both honest misses.

#### furpb (fur, alternative TSS)

furpb is an alternative TSS of the same *fur* promoter as furpa, shifted by 7 bp. The annotated binding sites are at identical genomic coordinates to those of furpa — redundant annotations of the same physical sites. See furpa for accounting.

#### galEp1 (gal operon, upstream promoter)

Two additional annotations fall outside our window (GalR at +46.5 and GalS at +52.5, the downstream OI operator of the gal bipartite operator system); excluded from in-window accounting.

| Position | TF | Function | Observed? | Reason / notes |
|---|---|---|---|---|
| −60.5 | GalR | repressor | yes | Operator engaged in acetate (no galactose), released in galactose. Same physical site as GalS. |
| −60.5 | GalS | repressor | yes | Same physical site as GalR; both paralog interactions supported. |
| −47 | H-NS | repressor | no | One of four H-NS annotations spanning −47 to −2; H-NS binds AT-rich regions as cooperative nucleoprotein filaments rather than at discrete operators (Bouffartigues et al. 2007; Dame 2005), so the four annotations likely correspond to contact points within one extended region. |
| −41.5 | CRP | activator | yes (weak) | Recovered in galactose; canonical class-I CRP activator position. |
| −36 | H-NS | repressor | no | Part of extended H-NS region (see −47). |
| −18 | H-NS | repressor | no | Part of extended H-NS region (see −47). |
| −2 | H-NS | repressor | no | Part of extended H-NS region (see −47). |
| +6.5 | HU | repressor | no | HU-degenerate exclusion (HU binds essentially sequence-independently; no HU site is recovered anywhere in our panel). |

**Accounting.** Eight in-window annotations covering **seven physical sites** after collapsing GalR/GalS at −60.5. Excluding HU under the dataset-wide exclusion leaves six testable physical sites; **3 recovered (GalR, GalS, CRP) + 4 H-NS not recovered** (treated as one extended region under cooperative-binding exclusion → 1 testable physical H-NS region not recovered, so **3/4 testable physical sites recovered**).

**References.**


- Bouffartigues, E., Buckle, M., Badaut, C., Travers, A., and Rimsky, S. (2007). *Nat Struct Mol Biol* 14:441–448. PMID 17435766.

- Dame, R.T. (2005). *Mol Microbiol* 56:858–870. PMID 15853876.

#### galEp2 (gal operon, alternative TSS)

galEp2 is an alternative TSS of the gal operon, shifted 5 bp relative to galEp1. All annotated binding sites are at identical genomic coordinates to those of galEp1 and correspond to the same physical sites. Some TFs are annotated with opposite regulatory functions at galEp1 vs galEp2 (CRP activator at galEp1, repressor at galEp2; GalR/GalS repressors at galEp1, activators at galEp2), reflecting the two-promoter switch at the gal operon. Our reporter measures combined output from both TSSs and cannot resolve the two directions separately. See galEp1 for accounting.

#### galSp (galS autoregulatory promoter)

Two additional annotations fall outside our window (GalR at +91.5 and GalS at +92.5, the downstream operator); excluded.

| Position | TF | Function | Observed? | Reason / notes |
|---|---|---|---|---|
| −61.5 | GalR | repressor | no | Both annotations W-confidence; galS autoregulatory operator may have lower intrinsic affinity than the main galE operator. A novel putative activator signal we detect further upstream (∼−95) may also compete for visibility in this region. |
| −61.5 | GalS | repressor | no | Same physical site as GalR (collapsed). |
| −42.5 | CRP | activator | yes | Canonical class-I CRP activator position. |

**Accounting.** Three in-window annotations, two physical sites after collapsing GalR/GalS. **1/2 physical sites recovered** (CRP recovered; GalR/GalS operator not).

#### gatYp (gat operon, galactitol utilization)

Only the CRP annotation has a defined position; ArcA and GatR annotations appear in RegulonDB without coordinates (gene-level regulatory evidence not localized to a binding site). GatR is a pseudogene in *E. coli* K-12 MG1655 and derivatives, interrupted by an IS3 insertion (Nobelmann and Lengeler 1996; Soupene et al. 2003) — non-functional in our strain background regardless.

| Position | TF | Function | Observed? | Reason / notes |
|---|---|---|---|---|
| −40.5 | CRP | activator | yes | Recovered. |
| (no pos) | ArcA | repressor | n/a | Annotation has no coordinate; not site-level testable. |

| Position | TF | Function | Observed? | Reason / notes |
|---|---|---|---|---|
| (no pos) | GatR | repressor | n/a | Pseudogene in MG1655; non-functional in our strain. |

**Accounting.** **1/1 recovered** (considering only annotations with defined positions).

**References.**


- Nobelmann, B. and Lengeler, J.W. (1996). *J Bacteriol* 178:6790–6795. PMID 8955298.
- Soupene, E. et al. (2003). *J Bacteriol* 185:5611–5626. PMID 12949114.


#### gyrAp (DNA gyrase A subunit)

| Position | TF | Function | Observed? | Reason / notes |
|---|---|---|---|---|
| −108 | CspA | activator | no | CspA-degenerate exclusion: CspA binds RNA/ssDNA cooperatively without specific sequence requirements (Jiang et al. 1997) and functions as a nucleic acid chaperone rather than a sequence-specific TF; same exclusion rationale as Fis and HU. |
| −88 | CspA | activator | no | CspA-degenerate exclusion (see −108). |
| −60 | CspA | activator | no | CspA-degenerate exclusion (see −108). |
| −42.5 | Fis | repressor | no | Fis-degenerate exclusion. |

**Accounting.** Four annotations, all under dataset-wide exclusions for nonspecific TFs. **No testable in-window sites remain**; gyrAp does not contribute to the site-level recovery tally.

**References.**


- Jiang, W., Hou, Y., and Inouye, M. (1997). *J Biol Chem* 272:196–202. PMID 8995247.
- Bae, W., Phadtare, S., Severinov, K., and Inouye, M. (1999). *Mol Microbiol* 31:1429–1441. PMID 10200963.


#### hdeAp (hdeA, acid-inducible periplasmic chaperone)

hdeAp and hdeAp2 are alternative TSS annotations of the same promoter; all annotated binding sites are at identical genomic coordinates and we treat hdeAp2 as fully redundant. GadW and GadX share DNA sites at gad regulatory regions (Tramonti et al. 2006), so the four annotations at −74.5 and the four at −53.5 collapse to two physical sites. Additional annotations outside our window (Lrp at −116, −159; H-NS at −118; GadE at −117.5; GadE/GadW/GadX at −126.5) are excluded.

After collapsing GadW/GadX paralogs, **12 in-window physical sites**. The promoter expresses in our data, but no TF-specific footprint is recovered at any annotated position. The mutational landscape is dominated by two strong variance hotspots in the −35 and −10 (improvements toward consensus σ70 elements) rather than TF-specific signals.

| Position | TF | Function | Observed? | Reason / notes |
|---|---|---|---|---|
| −97 | Lrp | repressor | no | W-confidence; no footprint in minimal-media conditions where Lrp should be active. Honest miss. |
| −74.5 | GadW/X | act+rep | no | Paralog pair, one physical site; acid conditions tested (pH 2.0/2.5, 1 h, acute); no footprint. Honest miss. |
| −64 | Lrp | repressor | no | W-confidence; same reason as −97. |
| −55 | Lrp | repressor | no | W-confidence; same reason as −97. |
| −53.5 | GadW/X | act+rep | no | Same as −74.5. |
| −51.5 | TorR | activator | n/a | Excluded: TorR activated by anaerobic TMAO respiration, not in our condition panel. |
| −39 | MarA | repressor | no | Salicylate tested (recovers MarA footprints elsewhere in our dataset); no footprint here. Honest miss. |
| −32 | PhoP | activator | no | Mg2+ limitation tested; no footprint. Honest miss. |
| −27.5 | TorR | activator | n/a | Excluded (see −51.5). |
| −18 | Lrp | repressor | no | W-confidence; same reason as −97. |
| +8 | FliZ | repressor | n/a | Excluded: no flagellar/motility-inducing condition in our panel. |
| +9.5 | GadE | activator | no | W-confidence; acid conditions tested; no footprint. Honest miss. |

The hdeA acid-fitness island requires the GadEWX cascade with stationary phase + acid pH simultaneously (Ma et al. 2002; Tramonti et al. 2002; Seo et al. 2015). Our acute acid-shock protocol (pH 2.0/2.5, 1 h on exponential cells) may not fully fire the RpoS-dependent Gad cascade, which is one possible reason for the systematic miss.

**Accounting.** 14 in-window annotations → 12 physical sites (GadW/GadX collapsed). Excluding 3 condition-not-tested (TorR ×2, FliZ) → **9 testable physical sites. 0 recovered.**

**References.**

- Ma, Z., Richard, H., Tucker, D.L., Conway, T., and Foster, J.W. (2002). *J Bacteriol* 184:7001–7012. PMID 12446650.
- Tramonti, A., De Canio, M., Delany, I., Scarlato, V., and De Biase, D. (2006). *J Bacteriol* 188:8118–8127. PMID 16980449.
- Tramonti, A., Visca, P., De Canio, M., Falconi, M., and De Biase, D. (2002). *J Bacteriol* 184:2603–2613. PMID 11976288.
- Seo, S.W., Kim, D., O'Brien, E.J., Szubin, R., and Palsson, B.O. (2015). *Nat Commun* 6:7970. PMID 26258987.


#### ihfAp4 (ihfA, IHF alpha subunit, autoregulated)

A third IHF annotation at +54 is outside our window and excluded. IHF autoregulates ihfA. Unlike HU, IHF is sequence-specific (consensus WATCAANNNNTTR) and its footprints are legitimately resolvable in Reg-Seq.

| Position | TF | Function | Observed? | Reason / notes |
|---|---|---|---|---|
| −47 | IHF | repressor | no | Honest miss; possible reasons include differential intrinsic affinity between the two sites or lower per-condition occupancy. |
| −21 | IHF | repressor | yes (weak) | Strongest in slow-growing conditions (galactose, acetate, stationary phase) — consistent with growth-rate-dependent IHF levels rising on entry to stationary phase (Ali Azam et al. 1999), and with autoregulation mediated by low-affinity IHF sites that engage only at high IHF levels (Aviv et al. 1994). |

**Accounting. 1/2 in-window physical sites recovered.**

**References.**


- Ali Azam, T., Iwata, A., Nishimura, A., Ueda, S., and Ishihama, A. (1999). *J Bacteriol* 181:6361–6370. PMID 10515926.
- Aviv, M., Giladi, H., Schreiber, G., Oppenheim, A.B., and Glaser, G. (1994). *Mol Microbiol* 14:1021–1031. PMID 7715442.


#### marRp (marRAB operon, multiple antibiotic resistance)

The marbox at −61.5 is a single physical site bound by three paralogs (MarA, Rob, SoxS), same as at acrZp. MarR binds two operators (site I at −18, site II at +17) — the canonical tandem-operator architecture; two physical sites. With marbox collapsed: **8 physical sites**.

| Position | TF | Function | Observed? | Reason / notes |
|---|---|---|---|---|
| −81 | Fis | activator | no | Fis-degenerate exclusion. |
| −72.5 | Cra | repressor | no | Honest miss; not recovered in acetate (Cra-active condition). |
| −61.5 | MarA | activator | yes (sal) | Recovered under salicylate — the one panel condition that both activates MarA and inactivates MarR (Martin and Rosner 1995; Alekshun and Levy 1999), so the promoter is derepressed. |
| −61.5 | Rob | activator | no | Honest miss under dipyridyl. Hypothesis: MarR continues to repress marRp (dipyridyl doesn't inactivate MarR), so even when Rob is cytoplasmically activated, the activator signal can't propagate through a repressed promoter — explains why the same marbox is recovered at acrZp (no MarR present) but not here. Not formally confirmed. |
| −61.5 | SoxS | activator | no | Same hypothesis as Rob; PMS doesn't inactivate MarR. |
| −52.5 | CRP | activator | no | Honest miss in CRP-favoring conditions. |
| −40.5 | AcrR | repressor | no | Honest miss. |
| −29 | CpxR | activator | n/a | Excluded as likely annotation error: −29 places a positive regulator deep in the core promoter overlapping the −35 (inconsistent with class-I activator architecture), and no footprint was observed in any Cpx-inducing condition. |
| −18 | MarR | repressor | yes | Site I, recovered. |
| +17 | MarR | repressor | no | Site II, not recovered. |

**Accounting.** 10 annotations → 8 physical sites (marbox collapsed). Excluding Fis (dataset-wide) and CpxR (annotation error) → **6 testable physical sites. 2 recovered** (MarA marbox under salicylate; MarR site I). If marbox paralogs are counted separately: **2/8 regulatory-interaction annotations recovered**.

**References.**


- Martin, R.G. and Rosner, J.L. (1995). *Proc Natl Acad Sci USA* 92:5456–5460. PMID 7777530.
- Alekshun, M.N. and Levy, S.B. (1999). *J Bacteriol* 181:4669–4672. PMID 10419969.


#### mglBp (mglBAC high-affinity galactose transport operon)

Same architecture as galSp: GalR/GalS share a single upstream operator with a class-I CRP activator site closer to the TSS. mglBp2 is an alternative TSS with identical genomic coordinates, treated as redundant.

| Position | TF | Function | Observed? | Reason / notes |
|---|---|---|---|---|
| −59.5 | GalR | repressor | no | W-confidence; mglB operator may have lower intrinsic affinity than the main galE operator (which we do recover). |
| −59.5 | GalS | repressor | no | Same physical site as GalR. |
| −41.5 | CRP | activator | yes | Canonical class-I CRP activator position. |

**Accounting.** Three annotations, two physical sites. **1/2 physical sites recovered.**

#### mhpRp1 (mhpR, 3-(3-hydroxyphenyl)propionate degradation regulator)

| Position | TF | Function | Observed? | Reason / notes |
|---|---|---|---|---|
| −40.5 | CRP | activator | yes | Canonical class-I CRP activator position. |

**Accounting. 1/1 recovered.**

#### mprAp (mprA/emrR, autorepressor of the emrRAB efflux operon)

MprA (= EmrR) is the autorepressor of the *emrRAB* multidrug efflux operon, a MarR-family phenolic-ligand-sensing regulator; binds an imperfect 9-3-9 inverted repeat spanning the core promoter and TSS (Xiong et al. 2000).

| Position | TF | Function | Observed? | Reason / notes |
|---|---|---|---|---|
| −10 | MprA | repressor | yes | Recovered as repressor; under salicylate the footprint disappears, consistent with direct ligand binding to MprA causing dissociation from the promoter (Brooun et al. 1999; Xiong et al. 2000). Disappearance reflects MprA release, not a switch to activator. |

**Accounting. 1/1 recovered.**

**References.**


- Lomovskaya, O., Lewis, K., and Matin, A. (1995). *J Bacteriol* 177:2328–2334. PMID 7730261.
- Brooun, A., Tomashek, J.J., and Lewis, K. (1999). *J Bacteriol* 181:5131–5133. PMID 10438794.
- Xiong, A., Gottman, A., Park, C., Baetens, M., Pandza, S., and Matin, A. (2000). *Antimicrob Agents Chemother* 44:2905–2907. PMID 10991887.


#### ompFp (ompF, outer membrane porin F)

ompFp has an unusually dense set of RegulonDB annotations: within ~60 bp between −108 and −50 there are in-window annotations for IHF, CpxR, CRP, RstA, OmpR (×3 positions), and a second CpxR site. Four annotations (CpxR at −93, CRP at −92.5, RstA at −92, OmpR at −90.5) fall within a 3 bp window and cannot be separated by positional mutational analysis. Several other sites are at the window edge or outside (−108, −121.5, and −176 to −430).

This promoter highlights a general limitation of site-level recovery tallies: when annotated sites overlap within a few base pairs, the mutational footprint contains contributions from multiple TFs that Reg-Seq cannot separate without additional genomic context. We exclude ompFp from the site-level recovery accounting on the grounds that the cluster is not resolvable within the current framework.

ompFp2 is an alternative TSS with identical genomic coordinates, treated as redundant.

**Accounting.** Excluded from the site-level recovery tally.

#### ompRp1 (ompR-envZ operon)

ompRp1, ompRp2, ompRp3, ompRp4 are four annotated TSSs at the same locus with identical binding-site coordinates; we treat ompRp2/3/4 as redundant of ompRp1. Tsui et al. 1991 named the IHF sites IHF-A (−102), IHF-B (−37), IHF-C (+8) and showed by site-specific mutagenesis that only IHF-C is functionally required for in vivo repression of ompB.

| Position | TF | Function | Observed? | Reason / notes |
|---|---|---|---|---|
| −102 | IHF | repressor | no | IHF-A; Tsui et al. 1991 showed inactivating IHF-A (with IHF-B) does not alter IHF-mediated repression — not functionally required. |
| −53 | CRP | repressor | yes (as activator) | Mutational footprint with **activator direction** in cold-shock conditions, not the annotated repressor direction. Matches Huang et al. 1992: same CRP site is repressor on transcripts whose −35 elements overlap the CRP footprint and activator on transcripts whose TSSs lie further downstream. Our reporter reads combined output; activator behavior likely reflects a downstream CRP-activated TSS in the ompB region. Not visible in stationary phase (RpoS-dominated). |
| −37 | IHF | repressor | no | IHF-B; overlaps the σ70 −35 region (mutational signal dominated by RNAP); also shown not functionally required by Tsui et al. 1991. |
| +8 | IHF | repressor | yes | IHF-C; recovered as a clear repressor footprint, consistent with Tsui et al. 1991 site-specific mutagenesis. |

**Accounting.** **2/4 recovered** (IHF-C; CRP as activator on a downstream transcript, biologically validated by Huang et al. 1992 even though direction differs from the ompRp1 annotation). IHF-A and IHF-B are non-essential per Tsui et al. 1991.

**References.**


- Tsui, P., Huang, L., and Freundlich, M. (1991). *J Bacteriol* 173:5800–5807. PMID 1885551.
- Huang, L., Tsui, P., and Freundlich, M. (1992). *J Bacteriol* 174:664–670. PMID 1310090.

##### recNp (recN, SOS regulon, double-strand break repair)

The DnaA sites overlap the LexA sites by ~2.5 bp, forming two overlapping regulatory modules. Wurihan et al. 2018 showed experimentally that DnaA represses recN independently of LexA, with overlapping DnaA-box and LexA-box at the recN promoter. The two annotated LexA sites differ in their match to the CTGT-N8-ACAG consensus (Fernandez De Henestrosa et al. 2000): LexA at −22.5 (CTGTATATAAAACCAG, 1 mismatch CCAG→ACAG); LexA at −0.5 (CTGTACACAATAACAG, perfect palindrome).

| Position | TF | Function | Observed? | Reason / notes |
|---|---|---|---|---|
| −25 | DnaA | repressor | n/a | Excluded as unresolvable: overlaps the −22.5 LexA box by ~2.5 bp; mutations disrupting DnaA also disrupt LexA. No condition specifically turns off DnaA while LexA is bound — total DnaA roughly growth-rate-independent in K-12 (Hansen et al. 1991) — and under SOS-inducing conditions (PMS, H2O2) where LexA is cleaved, no DnaA footprint appears either. |
| −22.5 | LexA | repressor | yes | Recovered broadly across non-SOS conditions; disappears under PMS/H2O2 (LexA cleavage). |
| −4 | DnaA | repressor | n/a | Excluded as unresolvable (see −25). |

| Position | TF | Function | Observed? | Reason / notes |
|---|---|---|---|---|
| −0.5 | LexA | repressor | yes (slow growth only) | Recovered only in galactose, acetate, stationary phase. Counterintuitive given perfect-consensus sequence. Hypothesis: site overlaps TSS / RNAP open-complex region, so LexA occupancy competes with RNAP loading. Under fast growth (frequent RNAP loading) the site is functionally masked; under slow growth, LexA occupies it more of the time. Disappears under SOS conditions. |

**Accounting. 2/2 testable physical sites recovered** (both LexA sites; DnaA annotations excluded as unresolvable due to operator overlap).

**References.**


- Fernandez De Henestrosa, A.R. et al. (2000). *Mol Microbiol* 35:1560–1572. PMID 10760155.
- Wurihan, Gezi, Brambilla, E., Wang, S., Sun, H., Fan, L., Shi, Y., Sclavi, B., and Morigen (2018). *Front Microbiol* 9:1212. PMID 29967594.
- Hansen, F.G., Atlung, T., Braun, R.E., Wright, A., Hughes, P., and Kohiyama, M. (1991). *J Bacteriol* 173:5194–5199. PMID 1860829.


#### relBp (relBE toxin-antitoxin operon)

RelB and RelBE annotations at +1.5 and +13.5 are the same physical sites (RelB alone and the RelB2E heterotrimer bind the same DNA); four physical sites total. Bøggild et al. 2012 showed structurally that relO is optimally bound by two adjacent RelB2E heterotrimers forming a RelE-RelB2-RelB2-RelE heterohexamer overlapping the TSS and −10. Belliveau et al. 2018 also identified the two central sites as the functional operator by Sort-Seq and DNA-affinity MS.

| Position | TF(s) | Function | Observed? | Reason / notes |
|---|---|---|---|---|
| −13.5 | RelBE | repressor | no | Flanking site; not required for the cooperative heterohexamer assembly. |
| +1.5 | RelB + RelBE | repressor | yes | Central relO operator — recovered as repressor. Physical site shared with +13.5. |

| Position | TF(s) | Function | Observed? | Reason / notes |
|---|---|---|---|---|
| +13.5 | RelB + RelBE | repressor | yes | Central relO operator — recovered as repressor. Same architecture as +1.5. |
| +28 | RelBE | repressor | no | Flanking site; not required for cooperative assembly. |

**Accounting.** Six annotations → **four physical sites**. **2/4 physical sites recovered** (the two central relO sites), consistent with structural and prior MPRA evidence that only the central pair forms the functional cooperative operator.

**References.**


- Belliveau, N.M. et al. (2018). *PNAS* 115:E4796–E4805. PMID 29728462.
- Bøggild, A., Sofos, N., Andersen, K.R., Feddersen, A., Easter, A.D., Passmore, L.A., and Brodersen, D.E. (2012). *Structure* 20:1641–1648. PMID 22981948.


#### rspAp (rspAB operon, putative D-mannose dehydrogenase)

YdfH was renamed **RspR** by Shimada et al. 2021 after gSELEX identified the rspAB operon as its single target. Sakihama et al. 2012 originally identified YdfH/RspR as the rspAB repressor by gel shift.

| Position | TF | Function | Observed? | Reason / notes |
|---|---|---|---|---|
| −60.5 | CRP | activator | yes | Canonical class-I CRP activator position. |
| −35 | YdfH (RspR) | repressor | yes | Recovered as repressor. |

**Accounting. 2/2 recovered.**

**References.**


- Sakihama, Y., Mizoguchi, H., Oshima, T., and Ogasawara, N. (2012). *Biosci Biotechnol Biochem* 76:1688–1693. PMID 22972332.
- Shimada, T., Ogasawara, H., Kobayashi, I., Kobayashi, N., and Ishihama, A. (2021). *Front Microbiol* 12:697803. PMID 34220787.


#### sulAp (sulA, SOS-induced cell division inhibitor)

A second RcdA annotation at +47.5 is outside our window and excluded.

| Position | TF | Function | Observed? | Reason / notes |
|---|---|---|---|---|
| −5 | LexA | repressor | yes | Canonical SOS box; sulA is one of the most strongly LexA-repressed genes in the SOS regulon. |

| Position | TF | Function | Observed? | Reason / notes |
|---|---|---|---|---|
| +33.5 | RcdA | repressor | no | Honest miss. Shimada et al. 2012 identified RcdA binding upstream of sulA by DNase-I footprinting after gSELEX. RcdA is primarily implicated in biofilm regulation via csgD; activity at sulA may depend on conditions absent from our panel. |

**Accounting. 1/2 in-window physical sites recovered.**

**References.**


- Shimada, T., Katayama, Y., Kawakita, S., Ogasawara, H., Nakano, M., Yamamoto, K., and Ishihama, A. (2012). *Microbiologyopen* 1:381–394. PMID 23233451.


#### tisBp (tisB, SOS-induced toxin, type I toxin-antitoxin tisB/istR)

| Position | TF | Function | Observed? | Reason / notes |
|---|---|---|---|---|
| −28 | LexA | repressor | yes | Recovered despite W-confidence annotation. |

**Accounting. 1/1 recovered.**

#### tnaCp (tnaLAB operon, tryptophanase, tryptophan degradation)

| Position | TF | Function | Observed? | Reason / notes |
|---|---|---|---|---|
| −102.5 | TorR | activator | n/a | Excluded as condition-not-tested. TorR activation requires both TMAO and anaerobic growth (Bordi et al. 2003); neither is in our panel. Bordi et al. identified TorR-protected regions at tnaLAB by DNase-I footprinting. |
| −59.5 | CRP | activator | yes | Canonical class-I CRP activator; tnaLAB is a classic catabolite-repressed operon. |

| Position | TF | Function | Observed? | Reason / notes |
|---|---|---|---|---|
| −55.5 | TorR | activator | n/a | Excluded (see −102.5). Note: this TorR site overlaps the −59.5 CRP site, so under inducing conditions TorR and CRP would compete for overlapping DNA. |

**Accounting.** Two TorR annotations excluded as condition-not-tested. **1/1 testable site recovered** (CRP).

**References.**


- Bordi, C., Théraulaz, L., Méjean, V., and Jourlin-Castelli, C. (2003). *Mol Microbiol* 48:211–223. PMID 12657056.


#### tolCp1, tolCp2, tolCp3, tolCp4 (tolC, outer membrane channel)

tolC has four promoters with transcription start sites spanning ~70 bp. p1 and p2 (Eguchi et al. 2003) lie upstream of the marbox and are not activated by MarA/SoxS/Rob; p2 is tolC has four promoters spanning ~70 bp. p1 and p2 (Eguchi et al. 2003) lie upstream of the marbox; p2 is activated by EvgAS and PhoPQ. p3 and p4 (Zhang et al. 2008) lie downstream of the marbox and are activated by MarA/SoxS/Rob via a single shared marbox. We built four constructs (each 160 bp) covering this spread of TSSs. The p3 and p4 marbox annotations correspond to the same physical site at genomic position 3178021.5 (Class II / Class I* configurations relative to their respective −10 elements). The p1 and p2 TSSs are 8 bp apart with the same upstream architecture (PhoP box, no marbox influence) — we treat tolCp1/tolCp2 as a single promoter and report from tolCp2.

| Construct | Position | TF | Function | Observed? | Reason / notes |
|---|---|---|---|---|---|
| tolCp2 | −46 | PhoP | activator | yes (1/2 reps) | Recovered in one of two $Mg^{2+}$-starvation replicates. PhoPQ activated by low $Mg^{2+}$. W-confidence and replicate variability consistent with weak but real footprint. |

| Construct | Position | TF | Function | Observed? | Reason / notes |
|---|---|---|---|---|---|
| tolCp3/p4 | shared marbox | Rob | activator | yes (in tolCp2 only) | Recovered via edge effect: p3/p4 −10 sits at downstream edge of tolCp2 window; reporter mRNA initiated from p3/p4 in tolCp2 has only a few bp of native downstream sequence before the barcode, so it escapes the SdsR mRNA destabilization (Parker and Gottesman 2016). Tested under dipyridyl (Rosner et al. 2002). |
| tolCp3/p4 | shared marbox | MarA | activator | no | Honest miss under salicylate. Zhang et al. 2008 reported only 2.0-fold tolC::lacZ activation by salicylate; effect size may fall near our detection threshold. |
| tolCp3/p4 | shared marbox | SoxS | activator | no | Honest miss under PMS. Zhang et al. 2008 reported 2.8-fold activation by paraquat; small effect size. |

**On the tolCp3 and tolCp4 constructs.** No marbox footprint is recovered in either p3 or p4 constructs (even for Rob), despite p3/p4 TSSs being centrally placed. Plausible explanation: the tolCp3/p4 mutagenized inserts extend past the TSSs into native downstream sequence containing the SdsR pairing site (33 nt upstream of RBS; Parker and Gottesman 2016), so reporter mRNA from p3/p4 in those constructs is destabilized by SdsR. The tolCp2 construct ends before this region. Hypothesis, not directly tested.

**Accounting.** Four annotated TFs (PhoP, Rob, MarA, SoxS). **2/4 recovered** (PhoP and Rob). p3/p4 construct results not counted separately — the marbox is the same physical site already assessed via tolCp2.

**References.**


- Aono, R., Tsukagoshi, N., and Yamamoto, M. (1998). *J Bacteriol* 180:938–944. PMID 9473049.
- Eguchi, Y., Oshima, T., Mori, H., Aono, R., Yamamoto, K., Ishihama, A., and Utsumi, R. (2003). *Microbiology* 149:2819–2828. PMID 14523115.
- Parker, A. and Gottesman, S. (2016). *J Bacteriol* 198:1101–1113. PMID 26811318.
- Zhang, A., Rosner, J.L., and Martin, R.G. (2008). *Mol Microbiol* 69:1450–1455. PMID 18673442.


#### uofp (uof-fur operon, iron uptake regulator under oxidative stress)

uofp drives expression of a short upstream ORF (uof) whose translation is coupled to fur translation (Večerek et al. 2007). RegulonDB annotations reflect direct transcriptional activation of uof-fur by OxyR under $H_2O_2$ stress (Zheng et al. 1999, who showed OxyR binds directly at the fur promoter via DNase-I footprinting).

| Position | TF | Function | Observed? | Reason / notes |
|---|---|---|---|---|
| −62 | OxyR | activator | no | Honest miss under $H_2O_2$. Both OxyR positions correspond to a single OxyR tetramer footprint (~45 bp, four ATAGnt elements at 10 bp intervals; Toledano et al. 1994). Despite direct-binding evidence (Zheng et al. 1999) and testing under $H_2O_2$, no mutational footprint detected. Effect size may fall below detection. |
| −61.5 | SoxS | activator | no | Honest miss under PMS. Zheng et al. 1999 explicitly localized SoxS-mediated activation of the fldA-fur locus to the *fldA* promoter, not uof/fur; this annotation likely reflects a low-affinity marbox from genome-wide screens rather than a physiologically relevant site. |
| −40 | OxyR | activator | no | Same OxyR tetramer footprint as −62 (see above). |

**Accounting. 0/3 recovered.**

**References.**


- Toledano, M.B., Kullik, I., Trinh, F., Baird, P.T., Schneider, T.D., and Storz, G. (1994). *Cell* 78:897–909. PMID 8087856.
- Večerek, B., Moll, I., and Bläsi, U. (2007). *EMBO J* 26:965–975. PMID 17268550.
- Zheng, M., Doan, B., Schneider, T.D., and Storz, G. (1999). *J Bacteriol* 181:4639–4643. PMID 10419964.


#### xylAp (xylAB operon, D-xylose isomerase and kinase)

xylAp drives the xylAB metabolic operon. XylR activates transcription in the presence of D-xylose (Song and Park 1997). AraC cross-represses xylAB when arabinose is present (Desai and Rao 2010; Groff et al. 2012). CRP provides catabolite activation. The two XylR positions correspond to the two direct repeats within the ~37 bp XylR operator IA — each XylR dimer contacts one direct repeat (Ni et al. 2013).

| Position | TF | Function | Observed? | Reason / notes |
|---|---|---|---|---|
| −81.5 | CRP | activator | yes | Canonical class-I upstream CRP position; consistent with catabolite activation. |
| −62.5 | XylR | activator | yes | Recovered under xylose. XylR binding requires D-xylose as effector. |
| −48 | AraC | repressor | no | Honest miss. AraC represses xylA only when arabinose-bound. We tested arabinose alone (no xylose) — condition where AraC should be ligand-loaded and active as repressor — but no footprint detected. |
| −41.5 | XylR | activator | yes | Recovered under xylose; second XylR direct repeat in operator IA. |
| −25 | AraC | repressor | no | Honest miss; W-confidence within RNAP-binding region. |

**Accounting.** Five annotations. **3/5 recovered** (CRP, both XylR sites).

**References.**


- Desai, T.A. and Rao, C.V. (2010). *Appl Environ Microbiol* 76:1524–1532. PMID 20023096.
- Groff, D., Benke, P.I., Batth, T.S., Bokinsky, G., Petzold, C.J., Adams, P.D., and Keasling, J.D. (2012). *Appl Environ Microbiol* 78:2221–2229. PMID 22286982.
- Ni, L., Tonthat, N.K., Chinnam, N., and Schumacher, M.A. (2013). *Nucleic Acids Res* 41:1998–2008. PMID 23248008.
- Song, S. and Park, C. (1997). *J Bacteriol* 179:7025–7032. PMID 9371449.

#### xylFp (xylFGHR operon, xylose ABC transporter and regulator)

xylFp drives the xylFGHR operon encoding the high-affinity xylose ABC transporter and XylR. Like xylAB, activated by XylR + D-xylose (Song and Park 1997). The XylR operator IF has two direct repeats in the same configuration as IA (Ni et al. 2013). A CRP annotation at −180.5 is outside our window and excluded.

| Position | TF | Function | Observed? | Reason / notes |
|---|---|---|---|---|
| −82 | Fis | repressor | n/a | Excluded — dataset-wide Fis exclusion (degenerate consensus). |
| −75 | Fis | repressor | n/a | Excluded — dataset-wide Fis exclusion. |
| −61.5 | XylR | activator | yes | Recovered under xylose; first direct repeat of operator IF. |
| −40.5 | XylR | activator | yes | Recovered under xylose; second direct repeat of operator IF. |
| +22 | Fis | repressor | n/a | Excluded — dataset-wide Fis exclusion. |

**Accounting.** Three Fis annotations excluded (dataset-wide). **2/2 testable physical sites recovered** (both XylR sites).

**References.**


- Ni, L., Tonthat, N.K., Chinnam, N., and Schumacher, M.A. (2013). *Nucleic Acids Res* 41:1998–2008. PMID 23248008.
- Song, S. and Park, C. (1997). *J Bacteriol* 179:7025–7032. PMID 9371449.


#### ykgRp (ykgR, small protein of unknown function)

ykgR is a small y-ome gene (47 aa). Hemm et al. 2010 confirmed YkgR protein expression and observed higher levels in minimal glycerol than glucose, consistent with CRP-mediated catabolite activation.

| Position | TF | Function | Observed? | Reason / notes |
|---|---|---|---|---|
| −41.5 | CRP | activator | yes | Recovered under non-glucose conditions; canonical class-II CRP activator position; consistent with Hemm et al. 2010. |

**Accounting. 1/1 recovered.**

**References.**


- Hemm, M.R., Paul, B.J., Miranda-Ríos, J., Zhang, A., Soltanzad, N., and Storz, G. (2010). *J Bacteriol* 192:46–58. PMID 19734316.

#### yncEp (yncE, iron-regulated periplasmic β-propeller protein)

yncE encodes a periplasmic β-propeller protein adjacent to and divergent from yncD (putative TonB-dependent iron transporter). yncE is strongly iron-repressed via Fur (up to 18-fold derepression under iron limitation; McHugh et al. 2003); a Fur-box is present upstream, though not annotated in RegulonDB.

| Position | TF | Function | Observed? | Reason / notes |
|---|---|---|---|---|
| −41.5 | MarA | activator | no | Honest miss under salicylate (physiological MarA inducer). Some activator-like signal in single replicates of other conditions but not reproducible. Annotation likely from marbox overexpression screens (Martin et al. 2008); since panel does include the MarA inducer, counted as genuine miss rather than overexpression-screen exclusion. |

**Accounting. 0/1 recovered.**

**References.**


- McHugh, J.P., Rodríguez-Quiñones, F., Abdul-Tehrani, H., Svistunenko, D.A., Poole, R.K., Cooper, C.E., and Andrews, S.C. (2003). *J Biol Chem* 278:29478–29486. PMID 12746439.


#### yqaEp (yqaE, inner membrane protein of unknown function)

yqaE encodes a small inner membrane protein, confirmed Cpx regulon member (Price and Raivio 2009; Raivio et al. 2013). CpxR binding at the yqaE promoter has been confirmed by EMSA (Vogt et al. 2014).

| Position | TF | Function | Observed? | Reason / notes |
|---|---|---|---|---|
| −48 | CpxR | activator | yes | Recovered under Cpx-inducing conditions. |

**Accounting. 1/1 recovered.**

**References.**


- Price, N.L. and Raivio, T.L. (2009). *J Bacteriol* 191:1798–1815. PMID 19103922.
- Raivio, T.L., Leblanc, S.K.D., and Price, N.L. (2013). *J Bacteriol* 195:2755–2767. PMID 23564175.
- Vogt, S.L., Evans, A.D., Guest, R.L., and Raivio, T.L. (2014). *J Bacteriol* 196:4229–4238. PMID 25246476.

#### znuCp (znuCB operon, zinc ABC transporter)

znuC is the ATPase component of the high-affinity ZnuABC zinc uptake system. znuA and znuCB are divergently transcribed from a shared bidirectional intergenic region with a single Zur binding site repressing both promoters (Patzer and Hantke 1998; Gilston et al. 2014). Under zinc-replete conditions, zinc-loaded Zur binds as a tetramer and blocks RNAP; under zinc limitation, apo-Zur dissociates and both operons derepress.

| Position | TF | Function | Observed? | Reason / notes |
|---|---|---|---|---|
| −27.5 | OxyR | activator | no | Honest miss under $H_2O_2$. W-confidence annotation overlapping RNAP-binding region. Warner and Levy 2012 identified SoxS (not OxyR) as the relevant activator of znuACB under host-relevant conditions in a murine pyelonephritis model — though their gel-shift experiments failed to show direct SoxS binding, suggesting indirect regulation. The OxyR annotation likely reflects low-affinity consensus matching rather than direct physiological regulation. |
| −10 | Zur | repressor | n/a | Excluded as condition-not-tested. Zinc limitation (TPEN or equivalent chelator) is not in our condition panel, so Zur remains zinc-loaded and bound throughout. Mutations disrupting the zur box would derepress only under zinc limitation. |

**Accounting.** Zur excluded as condition-not-tested. **0/1 testable site recovered.**

**References.**


- Gilston, B.A., Wang, S., Marcus, M.D., Canalizo-Hernández, M.A., Swindell, E.P., Xue, Y., Mondragón, A., and O'Halloran, T.V. (2014). *PLOS Biol* 12:e1001987. PMID 25369000.
- Patzer, S.I. and Hantke, K. (1998). *Mol Microbiol* 28:1199–1210. PMID 9680209.
- Sabri, M., Houle, S., and Dozois, C.M. (2009). *Infect Immun* 77:1155–1164. PMID 19103764.
- Warner, D.M. and Levy, S.B. (2012). *J Bacteriol* 194:1177–1185. PMID 22210763.
